# Comparative Analysis of SARS-CoV-2 Antigenicity across Assays and in Human and Animal Model Sera

**DOI:** 10.1101/2023.09.27.559689

**Authors:** Barbara Mühlemann, Samuel H Wilks, Lauren Baracco, Meriem Bekliz, Juan Manuel Carreño, Victor M Corman, Meredith E. Davis-Gardner, Wanwisa Dejnirattisai, Michael S Diamond, Daniel C. Douek, Christian Drosten, Isabella Eckerle, Venkata-Viswanadh Edara, Madison Ellis, Ron A M Fouchier, Matthew Frieman, Sucheta Godbole, Bart Haagmans, Peter J Halfmann, Amy R Henry, Terry C Jones, Leah C Katzelnick, Yoshihiro Kawaoka, Janine Kimpel, Florian Krammer, Lilin Lai, Chang Liu, Sabrina Lusvarghi, Benjamin Meyer, Juthathip Mongkolsapaya, David C Montefiori, Anna Mykytyn, Antonia Netzl, Simon Pollett, Annika Rössler, Gavin R Screaton, Xiaoying Shen, Alex Sigal, Viviana Simon, Rahul Subramanian, Piyada Supasa, Mehul Suthar, Sina Türeli, Wei Wang, Carol D Weiss, Derek J Smith

## Abstract

The antigenic evolution of SARS-CoV-2 requires ongoing monitoring to judge the immune escape of newly arising variants. A surveillance system necessitates an understanding of differences in neutralization titers measured in different assays and using human and animal sera. We compared 18 datasets generated using human, hamster, and mouse sera, and six different neutralization assays. Titer magnitude was lowest in human, intermediate in hamster, and highest in mouse sera. Fold change, immunodominance patterns and antigenic maps were similar among sera. Most assays yielded similar results, except for differences in fold change in cytopathic effect assays. Not enough data was available for conclusively judging mouse sera, but hamster sera were a consistent surrogate for human first-infection sera.

## Introduction

The assessment of antigenic differences among SARS-CoV-2 variants is of critical importance for interpreting the genetic evolution of the virus and for judging the need for vaccine updates. Understanding the comparability of data generated in different laboratories using different neutralization assays and human and animal sera would create a solid foundation for a surveillance system that combines data from laboratories throughout the world.

Multiple laboratories have tested a wide range of variants against primary infection or vaccination sera from species including humans, hamsters, and mice, using a variety of neutralization assays (*1–12*). In the early phases of the pandemic, it was not clear which assay(s) would measure the most relevant antigenic differences among variants for assessing protection in humans, and there was value in multiple laboratories using different methods. Comparisons of different neutralization assays in the same laboratory have shown that titers between assays broadly correlate (*4*, *13–15*). A rough correspondence has also been shown between titers generated by the same laboratory using authentic SARS-CoV-2 and vesicular stomatitis virus (VSV) pseudotypes using plaque reduction neutralization tests (PRNT) on Vero E6 and Calu-3 cells (*4*). However, differences in fold change have been observed for BA.1, BA.2, BA.2.12.1, and BA.4/BA.5 VSV pseudotypes titrated in Vero E6/TMPRSS2 and HEK293T/ACE2 cells (*16*). Further, neutralization titers measured against the B.1 variant using lentivirus pseudotype neutralization assays in the same sera in two clinically approved laboratories were higher in the assay that used TMPRSS2 overexpressing cells (*17*). Using the first available neutralization data of the Omicron BA.1 variant drawn from 23 laboratories, Netzl et al., 2022 found that pseudovirus-based assays measure two or three times higher geometric mean neutralization titers (GMT) against the ancestral and BA.1 variants than authentic virus assays in sera from vaccinees immunized with two or three times, respectively, but observed no substantial difference in fold change between pseudovirus and live virus assays (*18*).

Human sera are the gold standard for assessing antigenic differences most relevant for escape from population immunity. Sera from individuals with first infections or vaccinations with different variants provide baseline information about antigenic relationships between SARS-CoV-2 variants (*3*, *5*, *8*, *12*, *19*, *20*), to which antigenic distances measured in sera with more diverse or unknown infection histories can be compared (*21*). Data from titrations of single-variant exposure sera against different variants can be described using antigenic cartography (*22*), providing a visualization of antigenic relationships between variants (*3*, *4*, *7*, *8*, *11*, *12*). However, with a large fraction of the population either vaccinated or previously infected, finding individuals with first infections or serum donors with known infection histories is now no longer possible (other than sera from young children, which is often difficult to obtain in large enough volumes, and identification of the infecting variant is complicated by co-circulating variants, reduced genomic surveillance, and possible earlier asymptomatic infections or maternal antibody transfer). An alternative is to use sera from experimentally infected animals, which allows for the exact control of the infecting variant. In that case, it is necessary to determine whether animal model sera accurately reflect human first-infection sera with regard to their ability to neutralize different SARS-CoV-2 variants.

To better understand variability of neutralization titers generated in different laboratories using different sera and assays, we analyze data from 18 studies of antigenic differences among SARS-CoV-2 variants. Each study tested at least five variants against human, hamster, or mouse primary infection or vaccination sera raised against at least two different virus variants. The 18 datasets were made with six different neutralization assays. We sought to answer two questions: (i) how well do animal model sera replicate patterns of reactivity in human sera, and (ii) how do the various neutralization assays compare to each other. Datasets are compared according to four parameters: titer magnitude, fold change from the homologous variant, patterns of immunodominance, and the antigenic maps constructed from these datasets. We also investigated how well a combined antigenic map constructed from a combination of the datasets corresponds to the antigenic maps constructed from each individual dataset.

## Results

We compared 18 datasets from 14 different laboratories (Table 1, Table S1, Supplementary Text, figs. S1-S25). The datasets were generated and shared within the National Institute of Health SARS-CoV-2 Assessment of Viral Evolution (NIH SAVE) consortium (*23*), and by outside collaborators. Datasets were generated using human (n=8), hamster (n=8), and mouse (n=2) sera, and six different neutralization assays (lentivirus pseudotype neutralization (n=3) (LV-PV-neut), VSV pseudotype neutralization (n=2) (VSV-PV-neut), focus reduction neutralization test (n=6) (FRNT), PRNT (n=3), microneutralization (n=1) (Microneut), and cytopathic effect / limiting dilution assays (n=3) (CPE)). Two datasets were generated using sera pooled from multiple individuals (Maryland (mouse sera) and Madison (pooled) (hamster sera)). The datasets contain between five and 21 variants and two to 13 groups of sera raised by infection or vaccination with different variants. For the comparisons, we considered titrations generated using six different serum groups (4 weeks post mRNA-1273 vaccination and convalescent D614G, B.1.1.7, B.1.351, P.1, and B.1.617.2 sera, Table S1) and 23 different variants that were titrated in at least two datasets.

**Table 1:**
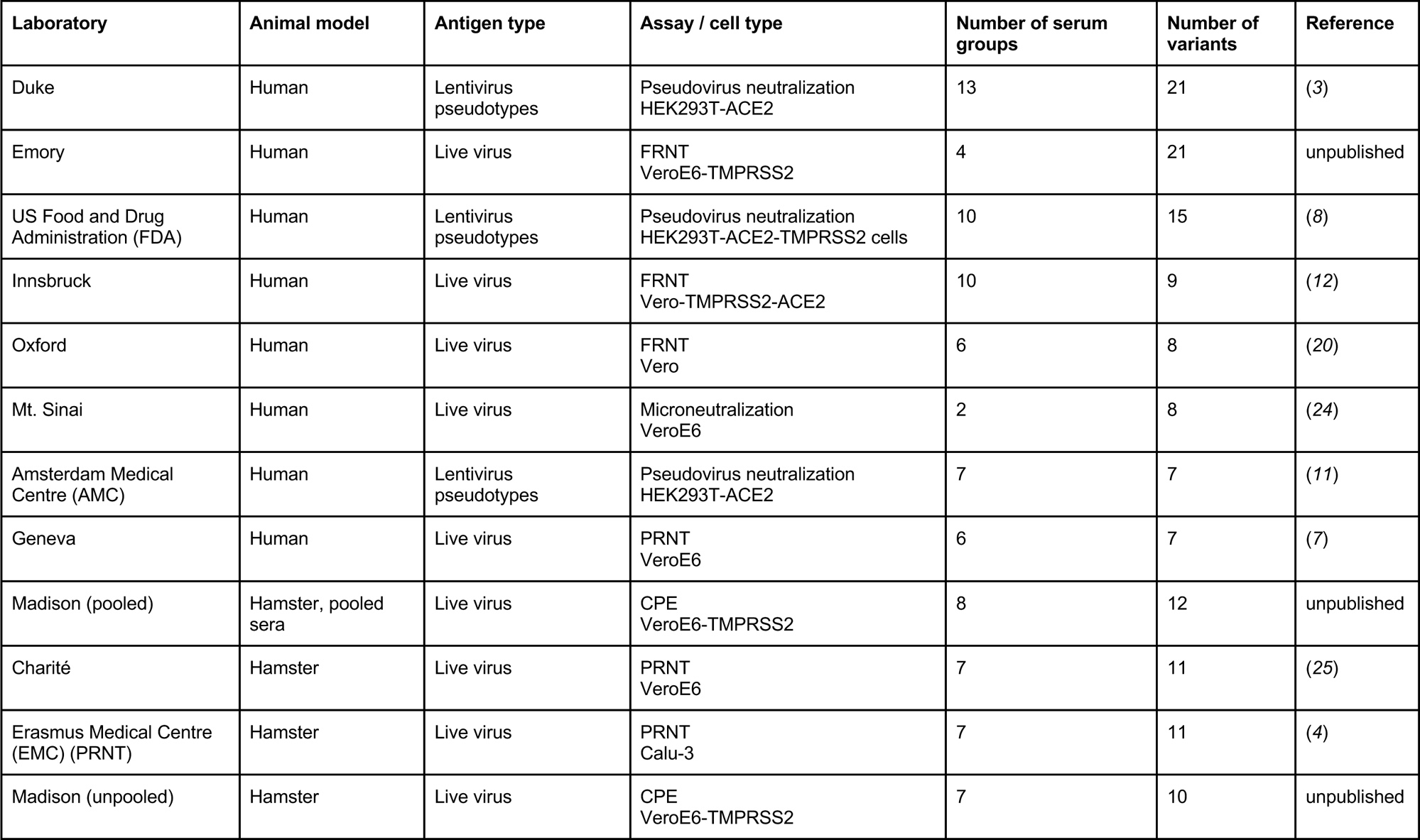
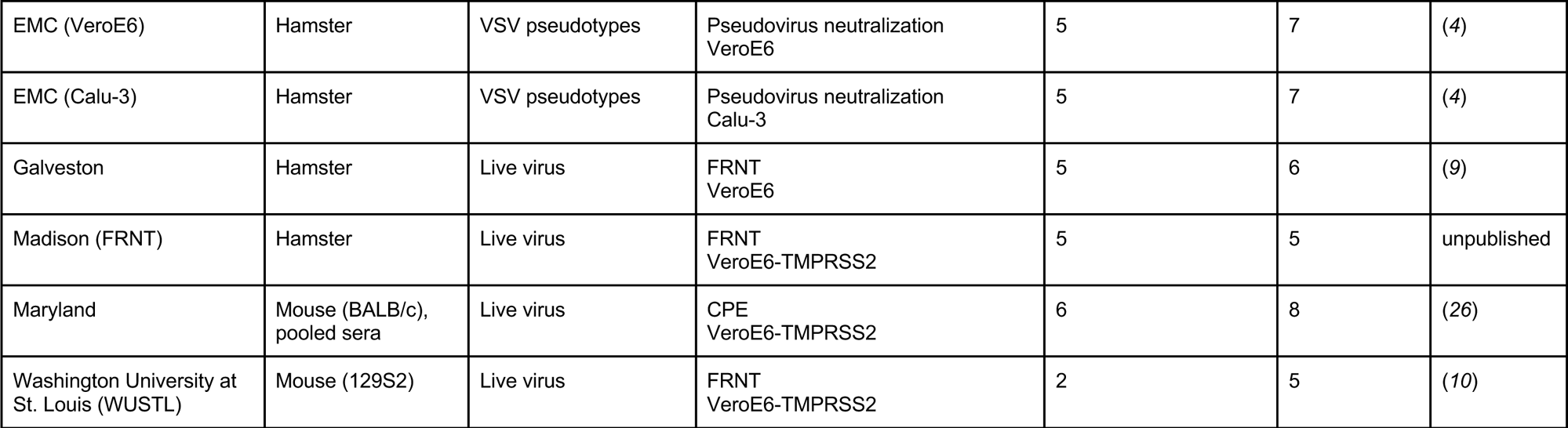
Overview of datasets used in this study. For detailed information on animal model sera and variants, see Table S1, Supplementary Text, figs. S1-S25.

| Laboratory | Animal model | Antigen type | Assay / cell type | Number of serum groups | Number of variants | Reference |
| --- | --- | --- | --- | --- | --- | --- |
| Duke | Human | Lentivirus pseudotypes | Pseudovirus neutralization HEK293T-ACE2 | 13 | 21 | (3) |
| Emory | Human | Live virus | FRNT VeroE6-TMPRSS2 | 4 | 21 | unpublished |
| US Food and Drug Administration (FDA) | Human | Lentivirus pseudotypes | Pseudovirus neutralization HEK293T-ACE2-TMPRSS2 cells | 10 | 15 | (8) |
| Innsbruck | Human | Live virus | FRNT Vero-TMPRSS2-ACE2 | 10 | 9 | (12) |
| Oxford | Human | Live virus | FRNT Vero | 6 | 8 | (20) |
| Mt. Sinai | Human | Live virus | Microneutralization VeroE6 | 2 | 8 | (24) |
| Amsterdam Medical Centre (AMC) | Human | Lentivirus pseudotypes | Pseudovirus neutralization HEK293T-ACE2 | 7 | 7 | (11) |
| Geneva | Human | Live virus | PRNT VeroE6 | 6 | 7 | (7) |
| Madison (pooled) | Hamster, pooled sera | Live virus | CPE VeroE6-TMPRSS2 | 8 | 12 | unpublished |
| Charité | Hamster | Live virus | PRNT VeroE6 | 7 | 11 | (25) |
| Erasmus Medical Centre (EMC) (PRNT) | Hamster | Live virus | PRNT Calu-3 | 7 | 11 | (4) |
| Madison (unpooled) | Hamster | Live virus | CPE VeroE6-TMPRSS2 | 7 | 10 | unpublished |
| EMC (VeroE6) | Hamster | VSV pseudotypes | Pseudovirus neutralization<br>VeroE6 | 5 | 7 | (4) |
| EMC (Calu-3) | Hamster | VSV pseudotypes | Pseudovirus neutralization<br>Calu-3 | 5 | 7 | (4) |
| Galveston | Hamster | Live virus | FRNT<br>VeroE6 | 5 | 6 | (9) |
| Madison (FRNT) | Hamster | Live virus | FRNT<br>VeroE6-TMPRSS2 | 5 | 5 | unpublished |
| Maryland | Mouse (BALB/c),<br>pooled sera | Live virus | CPE<br>VeroE6-TMPRSS2 | 6 | 8 | (26) |
| Washington University at<br>St. Louis (WUSTL) | Mouse (129S2) | Live virus | FRNT<br>VeroE6-TMPRSS2 | 2 | 5 | (10) |

### Titer magnitude

Titer magnitude can vary between individual sera that show otherwise similar patterns of reactivity. We therefore investigated overall differences in titer magnitude between the human, hamster, and mouse sera. Titers were generally lowest for human sera, and highest for mouse sera (∼6.7-fold higher than human titers), with those for hamster sera being intermediate (∼3.1-fold higher than human titers) (Fig. 1A, 1B, figs. S26-S28, S29A-B). The effect cannot be explained by differences in assays used (Fig. 1B, figs. S28, S29A,C).

**Figure 1:**
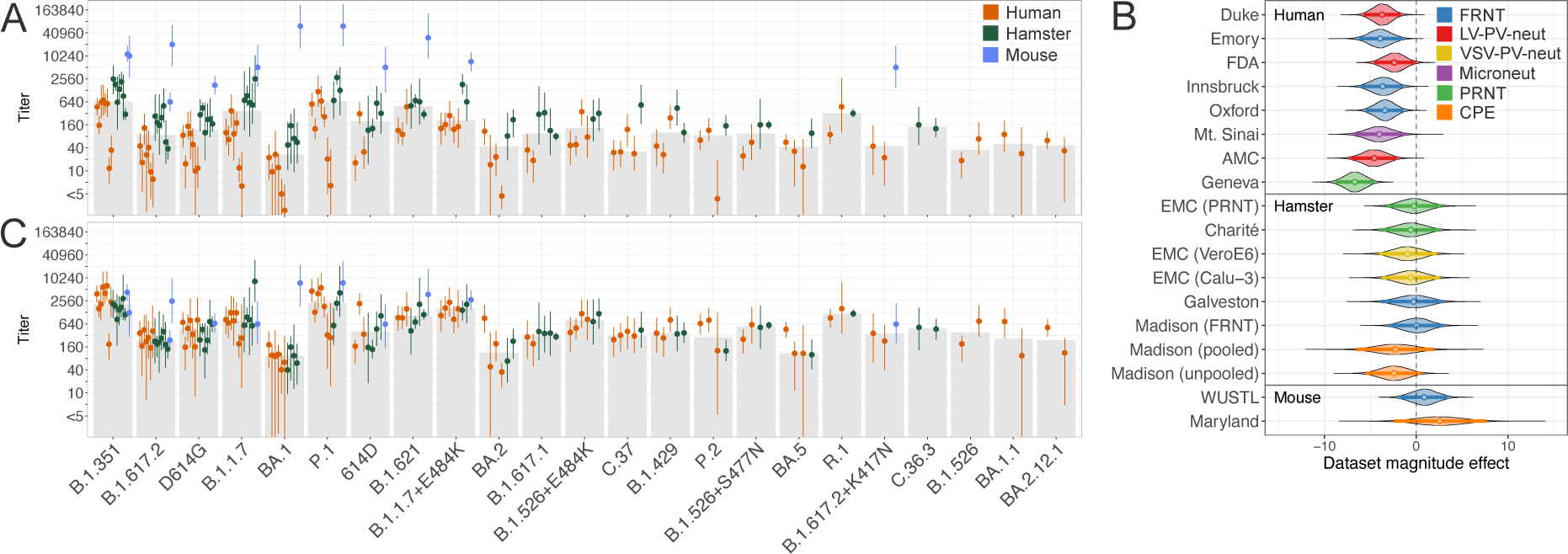
Comparison of titers between datasets as exemplified by the B.1.351 convalescent serum group. (**A**) Raw titers for the B.1.351 convalescent serum group. (**B**) Posterior distribution of the dataset magnitude effect. Datasets are grouped by animal model on the y-axis and coloured by assay (blue: FRNT, red: LV-PV-neut, yellow: VSV-PV-neut, purple: Microneut, green: PRNT, orange: CPE). Vertical bars show the 95% highest posterior density interval, each with a colored dot denoting the mean. (**C**) Titers after adjusting for dataset magnitude effects. In **A** and **C**, each dot corresponds to the GMT of a variant titrated against all B.1.351 sera in a particular dataset, GMTs in panel **C** were calculated from titers adjusted for dataset magnitude effect. Dots are colored by the animal model (red: human, green: hamster, blue: mouse). The gray bar heights indicate the median of the GMTs of the individual datasets. Equivalent figures for the other five serum groups can be found in figs. S27 (titer magnitude) and S31 (titer variability).

To control for the difference in magnitude among the raw titers (Fig. 1A, figs. S26-S28), we adjusted titrations for systematic differences in titer magnitude between datasets. To do so, we modeled each titer as a combination of the overall geometric mean titer for a variant and serum group, a reactivity bias of each serum to account for individuals generating an immune response leading to generally higher or lower titers, and a per-dataset magnitude effect (Materials and Methods). After adjusting each titer by the estimated per-dataset magnitude effect, there is no systematic difference in variation around a common GMT in titrations generated using human and hamster sera, with higher variability in the two mouse sera datasets (Fig. 1C, figs. S30, S31, S33A) and no difference in variation using different assays (figs. S32, S33B).

Therefore, these data indicate that after adjusting for titer magnitude, there is no evidence for systematic differences among hamster, mouse or human sera. Indeed, due to their higher titers, the dynamic range of measurement is greater before reaching the limit of detection of the assay in the hamster and mouse sera, making them more useful than human sera for measuring differences between antigenically diverse variants.

### Fold change

The fold change in neutralization between different variants titrated against the same serum gives a representation of the antigenic distance between the variants, irrespective of titer magnitude differences. We investigated whether the fold change measured from the homologous variant to other variants differed systematically by animal model or assay (Fig. 2, figs. S34-S38). Overall, the datasets roughly agree in the rank order of fold change measured for the different variants, with the exception of the Maryland and Madison (pooled) datasets, both generated using a CPE assay (Fig. 2, figs. S34-S35, S39). The amount of fold change differed between datasets. In particular, datasets generated using CPE assays measured less fold change than other datasets, whereas datasets generated using lentivirus pseudotype neutralization assays showed greater fold changes (Fig. 2, fig. S35, S40, S41).The amount of fold change measured using human, hamster, or mouse sera is similar, although insignificantly higher for the human datasets (fig. S41). Consequently, we find that the hamster model corresponds well with human sera with regards to fold change, with not enough data for conclusions about sera from the mouse models.

**Figure 2:**
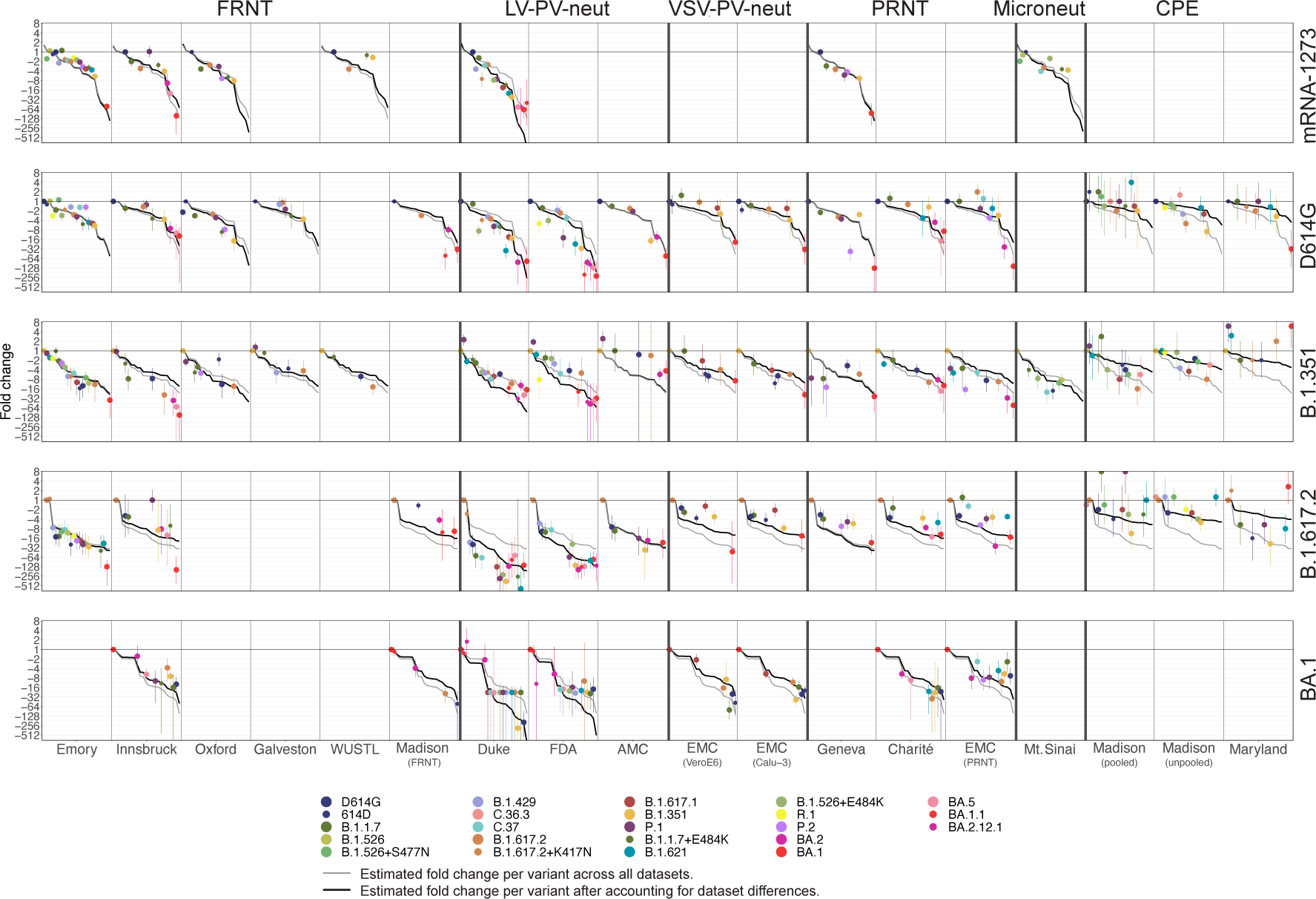
Fold change in the mRNA-1273, D614G, B.1.351, and B.1.617.2 convalescent serum groups measured in the 18 datasets. Dots show the mean estimated fold change for each variant and the bars show the estimated 95% highest posterior density intervals, with the colors corresponding to the variant. Datasets are grouped on the x-axis with bold vertical lines separating the FRNT, LV-PV-neut, VSV-PV-neut, PRNT, Microneut, and CPE datasets. The light gray line in each panel indicates the estimated fold change per variant in a particular serum group calculated across all datasets, in descending order. The black line shows the estimated fold change per variant in descending order after accounting for differences in fold change between datasets (Material and Methods). Variants are ordered on the x-axis by decreasing estimated fold change, calculated in a particular serum group across all datasets. Equivalent figures that include the B.1.1.7 and P.1 convalescent serum groups, as well as ordered by animal model sera are shown in figs. S30 and S31, respectively. Figures split by variant are shown in figures S36 (by dataset), S37 (by animal model), and S38 (by assay).

### Immunodominance patterns

Several studies have found evidence for variation in the sensitivity of sera to substitutions at different positions in the spike protein, consistent with changes in the immunodominance of different sites depending on the infecting variant (*3*, *27*, *28*). For example, sera raised against the ancestral variants are sensitive to substitutions at position 484 yet relatively insensitive to substitutions at position 501, whereas B.1.351 convalescent sera show an opposite pattern, with substitutions at position 484 having little effect on reactivity and changes at position 501 having more effect (*3*, *27*, *28*). There is increasing awareness that such changes in immunodominance may affect the epitopes preferentially targeted by an individual’s immune response, and hence its level of protection against subsequently circulating variants (*3*). We investigated whether consistent patterns of immunodominance could be observed in the datasets analyzed here with regards to sensitivity to substitutions at positions 484 and 501 already described for human sera.

Figure 3 shows fold differences between pairs of variants differing either at position 484 (E to K, panel A) or 501 (N to Y, panel B) titrated against six groups of sera. Figures S42-S48 show the same data split by animal model, assay, and cell type. In general, we find evidence that antibodies from mRNA-1273-immunized and D614G and B.1.1.7 convalescent sera are escaped by substitutions at position 484 (no mouse sera were available for this comparison), whereas neutralization by P.1 convalescent sera is not affected by the E484K substitution (Fig. 3A, fig. S43, S44). However, we find larger differences between datasets in the effect of the E484K substitution in B.1.351 convalescent sera, where the FDA, Emory, and Innsbruck datasets see a strong increase of titers with presence of E484K, in contrast to other datasets (Fig. 3A, figs. S43, 44). The FDA, Emory, and Innsbruck datasets were all made with TMPRSS2 overexpressing cell lines, possibly suggesting an effect of TMPRSS2 overexpression on the reactivity to the E484K substitution in B.1.351 sera (figs. S47, Table 1). However, the same pattern is not present in the Madison (pooled) and Madison (unpooled) datasets, which were generated using CPE assays and TMPRSS2 overexpressing VeroE6 cells, and is also absent in the EMC (PRNT) dataset, which was generated using Calu-3 cells that naturally express TMPRSS2.

**Figure 3:**
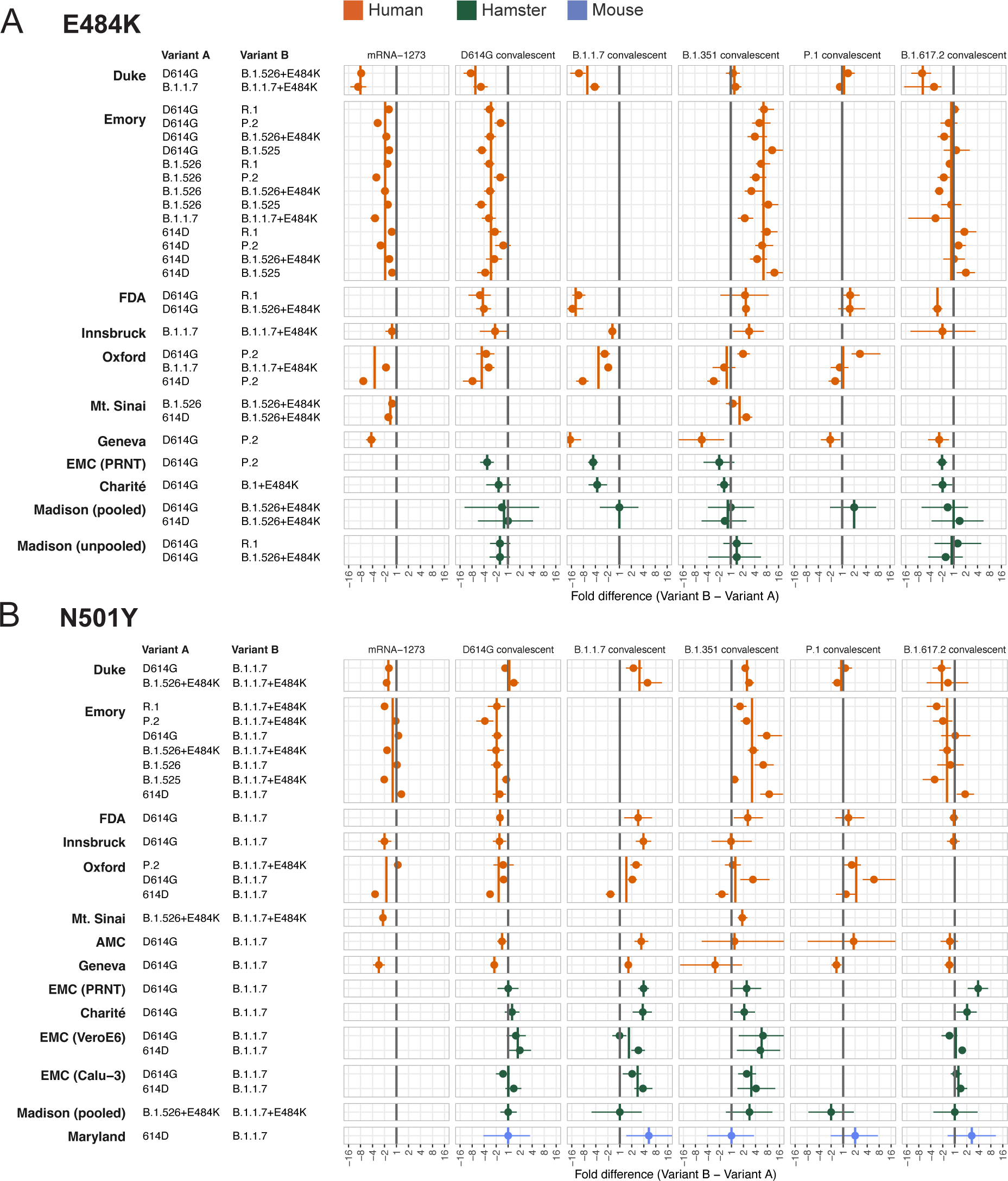
Sensitivity of different serum groups to the E484K and N501Y substitutions. Each row shows the average fold difference in titer between the two variants on the right, which differ by only the E484K (**A**) or the N501Y (**B**) substitution in the receptor binding domain (RBD). A positive fold difference corresponds to higher titers against the variant with the substitution, while a negative fold difference corresponds to higher titers against the variant without the substitution. Symbols and ranges correspond to the average fold difference and 95% highest posterior density intervals, calculated as described in the Methods. Data are colored by animal model (red: human, green: hamster, blue: mouse). Vertical lines indicate the average fold difference in each dataset, colored by animal model (red: human, green: hamster, blue: mouse). The gray line indicates no difference in titers between variants. Figure S42 shows the same figure split by assay.

When considering effects of the N501Y substitution (Fig. 3B, fig. S45), we find that on average, B.1.1.7 and B.1.351 convalescent sera are sensitive to the N501Y substitution in both human and hamster sera, whereas mRNA-1273, D614G, P.1, and B.1.617.2 convalescent sera are largely insensitive. There were no differences relating to the assay or the cell type that was used (figs. S42, S46, S48).

Generally, our findings of the reactivity of different serum groups correspond to that found in Wilks et al. (*3*), however, while Wilks et al., 2022 found evidence for spike positions other than 484 and 501 being preferentially targeted depending on first-infecting variant (*3*), we do not have sufficient data to test these here. Overall, sera from different animal models and titrated in different assays exhibit broadly similar patterns of immunodominance, with the exception of differences in the reactivity of the E484K substitution in B.1.351 between datasets, not directly attributable to the animal model, assay, or cell type used.

### Antigenic cartography

Antigenic maps provide a visual summary of titration data and can highlight patterns of reactivity to different serum groups not easily identified from titer tables (*22*). They estimate the base antigenic distances among variants, taking fold change compared to the maximum titer of each serum as input. This makes antigenic maps insensitive to differences in titer magnitude between datasets. For accurate placement in the antigenic map, each variant and serum should be determined by at least three titrations. This ‘geometric averaging’ of the data can mitigate the impact of noisy assay measurement, making the maps robust to some degree of titration error (*22*). Variants that are closely related antigenically and that have been titrated against a common set of sera will be positioned proximally in the map. Antigenic maps thus allow the easy identification of candidate vaccine variants that most closely represent the currently circulating diversity. We investigated whether antigenic maps constructed from the 18 datasets analyzed here show broadly similar topologies with regards to the placement of the main variants.

All maps show a similar topology of the ancestral, B.1.351, B.1.617.2 and Omicron BA.1 variants (Fig. 4A), with the ancestral variant occupying a central position, B.1.351 positioned towards the top, B.1.617.2 towards the bottom, and BA.1 furthest from D614G and mRNA-1273 vaccine sera, towards the right. Where present, Omicron BA.5 was consistently placed at the top of Omicron BA.1. Three maps (Madison (pooled), Madison (unpooled), and Maryland, third row of Fig. 4A) constructed from titers generated using CPE assays, differ more substantially from the other maps – this was to some degree anticipated given that these datasets exhibit discrepancies in rank order and amount of fold change between variants in at least one serum group (Fig. 2, fig. S34-S35, S40) compared to the other datasets.

**Figure 4:**
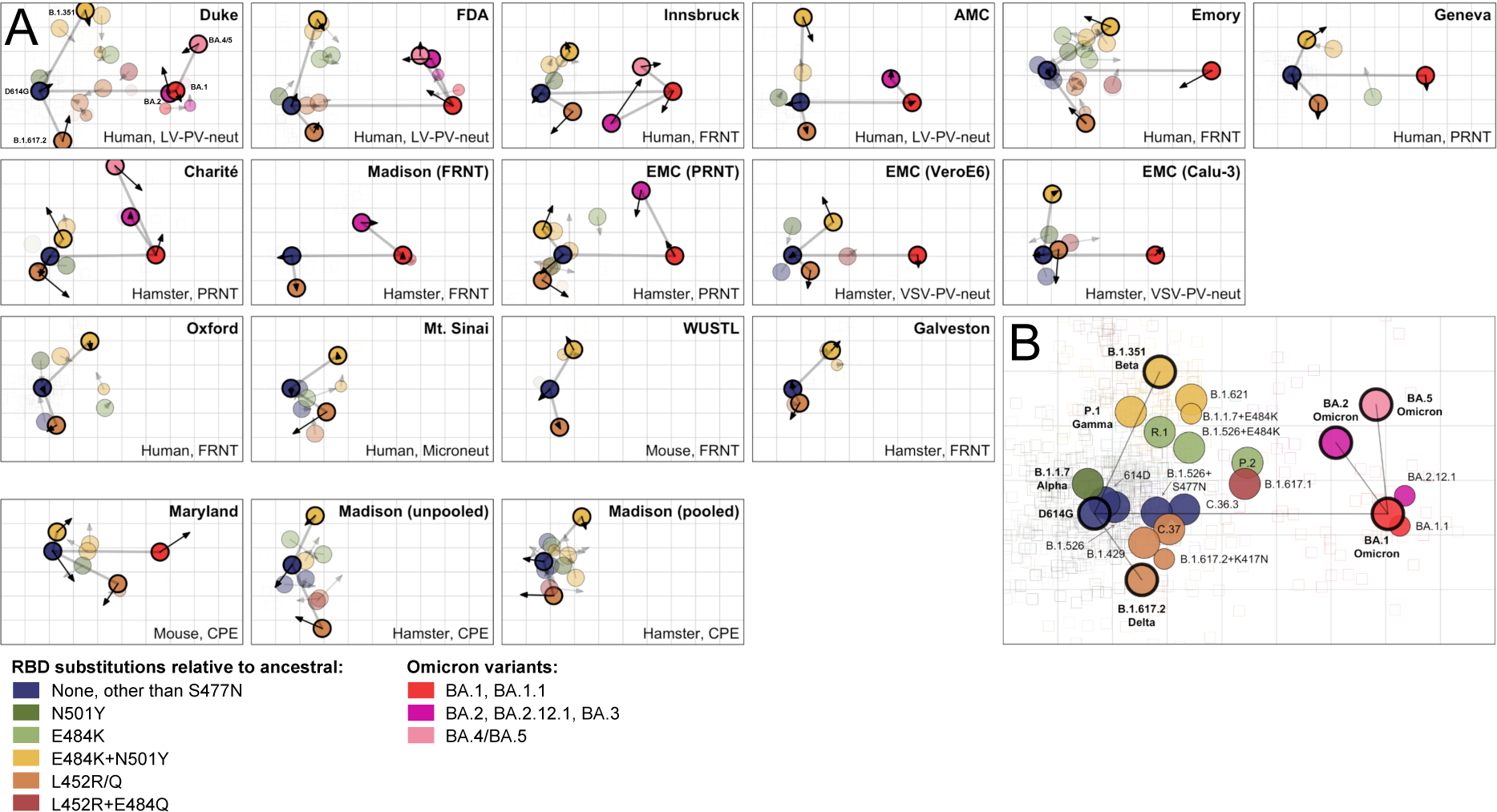
Antigenic map comparison. (**A**) Antigenic maps for each of the 18 datasets. Arrows point to the position of each variant in the merged map shown in panel B. Data for the maps in the bottom row were generated using CPE assays. (**B**) Antigenic map constructed from merging all titers of the 18 datasets. D614G, B.1.351, B.1.617.2, BA.1, BA.2 and BA.4/5 are highlighted. A version of this map colored by variant is shown in fig. S53. In both panels, variants are colored by substitutions in the RBD. Blue: no substitutions in RBD relative to the ancestral variant, except S477N. Includes ancestral (D614G and 614D), A.23.1, B.1.526, and B.1.526+S477N variants. Dark green: variants with only the N501Y substitution in the RBD. Includes B.1.1.7 and D614G+N501Y variants. Light green: variants with only the E484K substitution in the RBD. Includes B.1.525, R.1, P.2, and B.1.526+E484K variants. Yellow: variants with the E484K and N501Y substitutions in the RBD. Includes P.1 (+K417T), B.1.351 (+K417N), B.1.1.7+E484K, and B.1.621 (+R346K) variants. Orange: variants with the L452R (or Q, in the case of C.37) substitution in the RBD: Includes C.36.3, C.37 (+F490S), B.1.429, B.1.617.2 (+T478K), B.1.617.2+K417N (+T478K), AY.4.2 (+T478K), AY.1+K417N (+T478K), and AY.2+K417N (+T478K) variants. Dark red: variants with L452R and E484Q: includes B.1.617.1 (+T478K), B.1.630 (+T478K), and AY.3+E484Q (+T478K) variants. Bright red: Omicron BA.1 and BA.1.1 variants. Magenta: Omicron BA.2, BA.2.12.1, and BA.3 variants. Light pink: Omicron BA.4 and BA.5 variants.

We considered differences in the relative spread of the pre-Omicron variants (as exemplified by the distance between the D614G, B.1.351, and B.1.617.2 variants), and the Omicron variants (BA.1 and BA.2). Antigenic maps made using PRNT assays and hamster sera (Charité, EMC) show a tighter clustering of the pre-Omicron variants and a looser clustering of the Omicron variants, whereas maps made using human sera and lentivirus pseudotypes (Duke, FDA, AMC) show the opposite (Fig. 4, fig. S50). The tighter clustering of the Omicron variants in the Duke, FDA, and AMC datasets may be related to the smaller number of Omicron sera (Duke: n=5, FDA: n=2, AMC: n=1), which in these datasets also measure less fold change between BA.1 and BA.2 than other datasets. Given the longer duration of the pandemic at the time of the circulation of BA.1, these sera may also be more likely to be affected by undetected earlier infections. This, combined with the small number of Omicron sera, may lead to lower confidence in the positioning of those variants in the antigenic maps (fig. S51) and possibly cause the tighter clustering of Omicron variants in those datasets. The larger distances between the pre-Omicron variants in the lentivirus pseudotype assay datasets is supported by the larger fold change measured for those datasets (Fig. 2, fig. S52). Overall, the largely similar relative position of variants in the different antigenic maps suggests that hamster sera can provide an adequate surrogate for human first-infection sera, but that the choice of assay may affect the relative distances between variants.

A global monitoring system of SARS-CoV-2 antigenic variability will benefit if titration data from different laboratories, animal model sera, and assays can be combined to give a reliable consensus representation of SARS-CoV-2 antigenic evolution. As distances in an antigenic map are insensitive to overall titer magnitude, datasets with differences in titer magnitude can be compared and merged. We therefore investigated how well an antigenic map constructed from all titrations from the 18 datasets reflects the antigenic map positions in each individual dataset. We merged datasets on a per-variant basis, where we treated the variants as equal across datasets, and each serum was considered individually. Although a two-dimensional antigenic map is easiest to interpret visually, antigenic maps can be constructed in different dimensions, with lower stress in higher dimensions. Under cross-validation, we found that the merged map (Fig. 4B, fig. S53) provides a good fit to the data in two dimensions (figs. S54, S55). The map was robust to assay noise (fig. S56) and missing titers (figs. S57-S59) and to the exclusion of individual datasets (figs. S60, S61), although the merged map systematically underestimates log_2_ titers by ∼1.66 (fig S58).

The merged map shows a similar overall topology to the individual antigenic maps (Fig. 4A, figs. S62-S64), with the exception of the antigenic maps made from CPE assay datasets. In agreement with observations elsewhere (*3*), pre-Omicron variants placed to the right of the ancestral variant have substitutions at position 484 (light green, yellow, dark red), with variants in the top right additionally having the N501Y substitution (yellow). Variants towards the bottom-left area have the L452R substitution (orange). Omicron variants are placed furthest to the right of the ancestral variant, with BA.2 and BA.5 placed towards the top of BA.1.

## Discussion

We have constructed a statistical framework to compare SARS-CoV-2 neutralization titers and applied it to data from 18 laboratories, investigating differences in titer magnitude and variability, fold change, immunodominance patterns, and antigenic interrelatedness using antigenic cartography. Titers in the human datasets were generally lowest, followed by the hamster datasets, with the highest titers in the datasets generated using mouse sera. We did not find systematic differences in titer magnitude related to the assay used. The higher titers for mouse and hamster sera may stem from a higher inoculum dose used for raising animal sera compared to natural infections or vaccinations in humans. As a proportion of body weight, the vaccine dose for mice is much higher than for humans. The dose used for hamster infections was likely also higher than that estimated in human-to-human transmission. In addition, animals were boosted in one mouse and one hamster dataset after the initial infection or vaccination (Charité and WUSTL), again possibly leading to higher titers. Although not investigated due to the absence of metadata, differences between human study populations (such as age, co-morbidities and disease severity) as well as the timing of serum collection will affect titer magnitude. For example, in the Galveston dataset, hamster sera were taken at days 14, 28, and 45 post infection, and homologous titers increased ∼2.32-fold between day 14 and 45 (*9*). As we find that the rank order of fold change is similar between different animal model sera, the higher titers in non-human sera are not necessarily disadvantageous, as they may allow the characterisation of antigenically diverse variants without reactivity falling below the limit of detection of the assay. Dataset-specific differences in titer magnitude can be effectively adjusted for, and the remaining titer variability is similar between the different human and animal sera and neutralization assays with the exception of the two mouse datasets, which are more variable, possibly due to the smaller amount of data.

We did not find evidence for systematic differences relating to human and animal model sera with regards to amount of fold change or rank order of variants. However, we found that datasets constructed using CPE assays measure less fold change with different rank order of variants compared to other assays, whereas lentivirus pseudotype datasets tend to measure larger fold changes but rank fold changes similarly. Likewise, antigenic map topology is similar for human, hamster, and mouse sera and for different assays, with the exception of maps constructed from CPE assay data that show a different arrangement of variants, and maps generated using lentivirus pseudotype assay data that show a larger spread of the pre-Omicron variants. Larger fold change and spread of the pre-Omicron variants in the lentivirus pseudotype assay datasets may be due to higher assay sensitivity to RBD targeting antibodies, especially since the three datasets used ACE2 overexpressing cell lines. The level of ACE2 expression has been shown to increase how much RBD targeting antibodies contribute to neutralization (*29*). Furthermore, changes to spike folding, cleavage, and density, may lead to differences compared to live virus assays (*30*, *31*). Since the fold changes measured in the lentivirus pseudotype assay datasets correspond well with the other assays (excluding the CPE assay) modulo a scale factor, differences to other datasets can be adjusted for by a linear scale factor as was done in (*3*). The different rank order and amount of fold change in CPE assays may be because those assays measure neutralization across multiple replication cycles, and titers in these assays correspond to a different endpoint, requiring ∼99% of the initial inoculum to be inhibited by antibodies, compared to a ∼50% or ∼90% inhibition measured in corresponding PRNT, FRNT, and single-cycle pseudovirus neutralization assays (*32*). The differences in rank order of fold change and antigenic map topology in the CPE assays, and the relative correspondence among the other datasets suggest that CPE assay datasets may not be suitable for long-term routine antigenic characterisation of SARS-CoV-2.

We find that hamster sera mostly recapitulate the immunodominance patterns seen in human sera for E484K and N501Y, with not enough data yet to tell for mouse sera. The variation in reactivity to the E484K substitution in B.1.351 sera between different datasets is not directly attributable to the animal model, assay, or cell type, showing that depending on the dataset, different conclusions about immunodominance patterns with regards to the E484K substitution may be reached. Head-to-head comparisons of the same sera and variants in different assays and different laboratories are required to further elucidate possible drivers of this effect.

The merged antigenic map generated from titrations from all datasets broadly replicates patterns observed in the individual maps. Similarly, Netzl et al., 2022, found that an antigenic map constructed from GMTs extracted from 23 studies was broadly similar to the map presented in Wilks et al., 2022 (*18*). This provides evidence that data from a variety of sources can be combined to provide a general overview of the evolution of SARS-CoV-2, which can be helpful for a surveillance system, especially when datasets from individual laboratories may be incomplete on their own.

The 18 datasets considered here are heterogeneous with regards to the type of sera and neutralization assays used, which does not allow us to elucidate all possible causes of the patterns observed, as not all types of sera have been titrated in each assay. Further, due to absence of data, we were only able to compare immunodominance patterns for the E484K and N501Y substitutions and the pre-Omicron serum groups. Finally, because of the very limited number of human first-infection sera to variants arising after Omicron BA.1, we limited our analyses to variants up to BA.5. Therefore, we were not able to compare serum reactivities and antigenic differences of later Omicron variants, which are antigenically more diverse than pre-Omicron variants in hamster sera (*25*, *33*) and for which there may be greater variability between first-infection animal sera and assays.

Despite these limitations, the results suggest that all assays performed similarly, except for the CPE assay, and the differences present between human and animal sera and among assays can be accounted for by linear scale factors. Hamster sera can therefore serve as a useful substitute for human first-infection sera, with more data needed to determine the same for mouse sera. To advance SARS-CoV-2 antigenic surveillance and address the limitations discussed here, two key areas of ongoing investigation are crucial. First, within-laboratory comparisons of neutralization titers when using the same sera in different assays, as well as different animal model sera using the same assay. Second, upon circulation of major new variants, sourcing of first-infection human sera, to continually monitor the validity of the animal model sera as a surrogate system. These results, combined with the statistical framework established here for the ongoing comparative analysis SARS-CoV-2 neutralization data from a network of collaborating laboratories, are key components of a coordinated global surveillance system for monitoring SARS-CoV-2 antigenic variation.

## Acknowledgements

We thank the authors of van Der Straten et al., 2022 (*11*) and Liu et al., 2022 (*9*) for kindly sharing their datasets. Human sera panels for the Emory dataset were kindly provided by Nadine Rouphael, Jens Wrammert, Rafick Sekaly, John Roback, Jesse J. Waggoner, Frances Eun-Hyung Lee, Ignacio Sanz, Grace Mantus, Lindsay E. Nyhoff, Connie Wei, Linda Roback, Shannon Bonds, Safiyyah Rasheed.

The FDA lab sera results were derived from the Epidemiology, Immunology, and Clinical Characteristics of Emerging Infectious Diseases with Pandemic Potential (EPICC) (*8*) cohort studies for which the authors acknowledge the contributions of Nusrat J Epsi, Anthony C Fries, Stephanie A Richard, David A Lindholm, Katrin Mende, Evan C Ewers, Derek T Larson, Rhonda E Colombo, Christopher J Colombo, Julia S Rozman, Alfred Smith, Tahaniyat Lalani, Catherine M Berjohn, Ryan C Maves, Milissa U Jones, Rupal Mody, Nikhil Huprikar, Jeffrey Livezey, David Saunders, Anuradha Ganesan, Mark P Simons, Christopher C Broder, David R Tribble, Eric D Laing, Brian K Agan, and Timothy H Burgess.

## Funding

SHW, ST, AN, TCJ, DJS: supported by the NIH NIAID Centers of Excellence for Influenza Research and Response (CEIRR) contract 75N93021C00014 as part of the SAVE program.

AN was supported by the Gates Cambridge Trust.

BM, VMC, CD, TCJ: Funded by German Federal Ministry of Education and Research through project DZIF (8040701710 and 8064701703) and VARIpath (01KI2021), and the German Federal Ministry of Health through project SeroVarCoV.

CD: ECDC project Aurorae (NP/21/2021/DPR/25121) and EU Hera project Durable (101102733).

VMC: Participant in the BIH-Charité Clinician Scientist Program funded by Charité - Universitätsmedizin Berlin and the Berlin Institute of Health.

MSD: National Institutes of Health R01 AI157155 and NIAID Centers of Excellence for Influenza Research and Response (CEIRR) contracts 75N93021C00014 and 75N93019C00051.

LCK: Supported in part by the Intramural Research Program of the National Institute of Allergy and Infectious Diseases.

RS: Appointment to the NIAID Emerging Leaders in Data Science Research Participation Program. The Emerging Leaders in Data Science Research Participation Program is administered by the Oak Ridge Institute for Science and Education (ORISE) through an interagency agreement between the U.S. Department of Energy (DOE) and NIAID. ORISE is managed by Oak Ridge Associated Universities (ORAU) under DOE contract number DE-SC0014664.

YK: Acknowledges support from a Research Program on Emerging and Reemerging Infectious Diseases (JP21fk0108552 and JP21fk0108615), a Project Promoting Support for Drug Discovery (JP21nf0101632), the Japan Program for Infectious Diseases Research and Infrastructure (JP22wm0125002), and the University of Tokyo Pandemic Preparedness, Infection and Advanced Research Center (UTOPIA) grant (JP223fa627001) from the Japan Agency for Medical Research and Development.

GRS: The Chinese Academy of Medical Sciences (CAMS) Innovation Fund for Medical Science (CIFMS), China (2018-I2M-2-002), Schmidt Futures, and Red Avenue Foundation. The Wellcome Centre for Human Genetics is supported by the Wellcome Trust (090532/Z/09/Z).

MS, VVE, MD-G, LL, ME, DCD: Supported in part by grants (NIH P51OD011132, 3U19AI057266-17S1, HHSN272201400004C, NIH/NIAID CEIRR under contract 75N93021C00017 to Emory University) from the National Institute of Allergy and Infectious Diseases (NIAID), National Institutes of Health (NIH), Emory Executive Vice President for Health Affairs Synergy Fund award, the Pediatric Research Alliance Center for Childhood Infections and Vaccines and Children’s Healthcare of Atlanta, COVID-Catalyst-I3 Funds from the Woodruff Health Sciences Center and Emory School of Medicine, Woodruff Health Sciences Center 2020 COVID-19 CURE Award. Funders played no role in the design and conduct of the study; collection, management, analysis, and interpretation of the data; preparation, review, or approval of the manuscript; and decision to submit the manuscript for publication.

DCD, ARH, SG: Funded in part by the intramural program of the National Institutes of Health.

AS: The cohort was supported by the Bill and Melinda Gates Foundation award INV-018944 (to A.S.). SP: Was supported by awards from the Defense Health Program (HU00012020067) and the National Institute of Allergy and Infectious Disease (HU00011920111). The EPICC protocol was executed by the Infectious Disease Clinical Research Program (IDCRP), a Department of Defense (DoD) program executed by the Uniformed Services University of the Health Sciences (USUHS) through a cooperative agreement by the Henry M. Jackson Foundation for the Advancement of Military Medicine, Inc. (HJF).

JK: NIH NIAID Centers of Excellence for Influenza Research and Response (CEIRR) contract 75N93021C00014 as part of the SAVE program, European Union’s Horizon 2020 research and innovation program under grant agreement No. 101016174, and the Austrian Science Fund (FWF) with the project number P35159-B.

## Author contributions

Conceptualization: BM, SHW, DJS

Formal Analysis: BM, SHW, AN, ST

Investigation: BM, SHW

Methodology: BM, SHW

Software: BM, SHW

Validation: BM, SHW

Visualization: BM, SHW

Writing - original draft: BM, SHW, DJS

Funding acquisition: DJS, CD, TCJ

Supervision: DJS

Resources: LB, MB, YMC, VMC, MED-G, WD, MSD, DCD, CD, IE, V-VE, ME, RAMF, MF, SG, BH, PJH, ARH, TCJ, LCK, YK, JK, FK, LL, CL, SL, BMe, JM, AM, SP, AR, GRS, AS, VS, RS, PS, MS, WW, CDW

All authors contributed to Data curation and Writing - review & editing

## Disclaimer

The views expressed are those of the authors and do not reflect the official policy of the USUHS, Department of the Army, Department of the Navy, the Department of the Air Force, the Department of Defense or the U.S. Government and the Henry M. Jackson Foundation for the Advancement of Military Medicine, Inc. (HJF). The investigators have adhered to the policies for protection of human subjects as prescribed in 45 CFR 46. SL, WW, and CDW are US Government employees. Title 17 U.S.C. §105 provides that ‘Copyright protection under this title is not available for any work of the United States Government.’ Title 17 U.S.C. §101 defines a U.S. Government work as a work prepared by a employee of the U.S. Government as part of that person’s official duties.

