## Supplementary material for "Comparative Analysis of SARS-CoV-2 Antigenicity across Assays and in Human and Animal Model Sera"

### Materials and Methods

#### Description of datasets

We analyzed 18 datasets shared by collaborating laboratories. Dataset details are given in Table 1, Table S1, Supplementary Text, and figs. S1-S25. Datasets are named after the institution of the Principal Investigator, with the assay used as an additional indicator, if a laboratory contributed multiple datasets. For the titer comparison, we compared titers from the following, most-commonly used, serum groups: mRNA-1273 (double vaccinated), D614G convalescent (this includes any sera labeled as 'wild-type sera', as well as from animals infected with prototype variants with D614G, or 614D (for datasets where only prototype virus with 614D were used for infection)); B.1.1.7 convalescent / infected, B.1.351 convalescent / infected (including mRNA-1273.351 sera in the Washington University in St. Louis (WUSTL) dataset); P.1 convalescent / infected; B.1.617.2 convalescent / infected; and BA.1 convalescent / infected. We only compared titers of the 23 variants titrated in at least two datasets (B.1.351, B.1.617.2, D614G, B.1.1.7, BA.1, P.1, 614D, B.1.621, B.1.1.7+E484K, BA.2, B.1.617.1, B.1.526+E484K, C.37, B.1.429, P.2, B.1.526+S477N, BA.5, R.1, B.1.617.2+K417N, C.36.3, B.1.526, BA.1.1, BA.2.12.1).

##### Removal of outlier sera

The datasets generated using human sera that we included here included sera from individuals with no known previous infections or vaccinations. Nevertheless, it is possible that previous infections were undetectable, asymptomatic, or unreported. Therefore, following Wilks et al., 2022 (*3*), we attempted to identify and exclude human sera with reactivity patterns indicative of possible previous infections from subsequent analyses. We used two criteria for identifying outliers: first, any serum where a titer measured against any variant was >2-fold higher than the titer measured against the homologous variant for that serum group. Second, we visually inspected serum reactivity profiles and any serum whose reactivity profile differed from the general trend of the other sera in the same serum group was considered an outlier and excluded from the study.

**Duke**

The 'Duke' dataset was provided by David Montefiori and Shaunna Shen from Duke University. The dataset and antigenic map were previously published by Wilks et al., 2022 (*3*), where detailed information about map construction and sensitivity analyses can be found. The full map is shown in fig. S1. Outlier removal was performed in Wilks et al., 2022 (*3*), resulting in the removal of 2 of 15 D614G convalescent sera, 1 of 14 B.1.1.7 convalescent sera, 2 of 32 mRNA-1273 sera, 3 of 19 B.1.351 convalescent sera, 4 of 17 P.1 convalescent sera, 4 of 28 B.1.617.2 convalescent sera, and 3 of 7 BA.1 convalescent sera (see fig. S1 in (*3*)).

**Emory**

The 'Emory' dataset was provided by Prof. Mehul Suthar from Emory University. The assay is described in Edara et al., 2022 (*34*). The reactivity of the A.23.1, B.1.617.2, and B.1.617.2+K417N variants was lowered by 1 on the $\log_2$ scale, as these variants displayed disproportionally high titers in all serum groups tested. The full map and sensitivity analyses are shown in fig. S2. The map was constructed using 1000 optimisations. Outlier removal resulted in the removal of 1 of 17 D614G convalescent sera, 1 of 16 B.1.351 convalescent sera, and 9 of 14 B.1.617.2 convalescent sera (fig. S3).

**FDA**

The Food and Drug Administration (FDA) dataset was originally published in (*8*), where detailed information about map construction and sensitivity analyses can be found. The full map after removing outlier sera is shown in fig. S4. Outlier removal resulted in the removal of 4 of 10 B.1.1.7 convalescent sera, 2 of 10 C.37 convalescent sera, and 3 of 25 D614G convalescent sera (fig. S5).

**Innsbruck**

The ‘Innsbruck’ dataset was provided by Dr. Janine Kimpel and Dr. Annika Rössler from the Medical University of Innsbruck. The antigenic map was originally published in Rössler et al., 2022 (*12*), where detailed information about map construction and sensitivity analyses can be found. The full map after removing outlier sera is shown in fig. S6. Outlier removal resulted in the removal of 1 of 5 B.1.617.2 convalescent sera, 1 of 10 AstraZeneca vaccine sera, 2 of 9 BA.2 convalescent sera, and 1 of 10 D614G convalescent sera (fig. S7).

**Oxford**

The ‘Oxford’ dataset was provided by Prof. Gavin Screaton at Oxford University. The raw data for this map was originally published in (*35*). The full map after removing outlier sera and sensitivity analyses is shown in fig. S8. The map was constructed using 1000 optimizations. Outlier removal resulted in the removal of 2 of 34 D614G convalescent sera, 2 of 18 B.1.1.7 convalescent sera, 4 of 14 P.1 convalescent sera, and 1 of 14 B.1.351 convalescent sera (fig. S9).

**Mt. Sinai**

The ‘Mt. Sinai’ dataset was provided by Prof. Florian Krammer and Prof. Viviana Simon at the Icahn School of Medicine at Mount Sinai in New York. The titrations against the 2x mRNA-1273 vaccine sera were published in Carreño et al., 2021 (*24*). B.1.351 sera were provided by Prof. Alex Sigal and were taken on average 28 days post-diagnosis (min: 19, max: 46). The full map and sensitivity analyses are shown in fig. S10. The map was constructed using 1000 optimizations. Outlier removal resulted in the removal of 3 of 30 mRNA-1273 sera (fig. S11).

**AMC**

The Amsterdam Medical Centre (AMC) dataset was originally published in (*11*), where detailed information about map construction and sensitivity analyses can be found. The full map after removing outlier sera is shown in fig. S12. Outlier removal resulted in the removal of 1 of 4 B.1.351 convalescent sera, 1 of 3 P.1 convalescent sera, and the single BA.1 convalescent serum (fig. S13).

**Geneva**

The 'Geneva' dataset was originally published in (*7*), where detailed information about map construction and sensitivity analyses can be found. The full map is shown in fig. S14. No outlier sera were identified (fig. S15).

**Madison (pooled)**

The 'Madison (pooled)' dataset was provided by Prof. Yoshihiro Kawaoka and Prof. Peter Halfmann at the University of Wisconsin-Madison. The sera used in this map were pooled from 3 hamsters. The data was generated using a CPE/limiting dilution assay described in Chiba et al., 2021 (*36*). The full map and sensitivity analyses are shown in fig. S16. The map was made with 1000 optimizations. The reactivity of B.1.1.7 and P.1 was lowered by 3 on the log scale, as these variants displayed disproportionally high titers in all serum groups tested. For the analyses of immunodominance switches shown in Fig. 3, pairwise comparisons involving B.1.1.7 were left out due to the higher reactivity of that variant.

**Charité**

The ‘Charité’ dataset was provided by Prof. Christian Drosten and Dr. Victor Corman at the Institute of Virology at Charité – Universitätsmedizin Berlin and is available as Mühlemann et al., 2023 (*25*). The full map is shown in fig. S17.

**EMC (PRNT)**

The Erasmus Medical Centre (EMC) (PRNT) dataset was provided by Prof. Bart Haagmans and Anna Mykytyn from the Erasmus Medical Centre in Rotterdam, Netherlands. The antigenic map was originally published by Mykytyn et al., 2022 (*4*), as Figure 4I. For the analyses presented here, we excluded the P.1 sera from the map and titer analyses, as their pattern of reactivity closely resembled the pattern of sera raised against B.1.617.2. The map used here was constructed using 1000 optimisations. Sensitivity analyses of the map can be found in Mykytyn et al., 2022 (*4*). The full map is shown in fig. S18.

**Madison (unpooled)**

The ‘Madison (unpooled)’ dataset was provided by Prof. Yoshihiro Kawaoka and Prof. Peter Halfmann at the University of Wisconsin-Madison. The data was generated using a CPE/limiting dilution assay described in Chiba et al., 2021 (*36*). The full map and sensitivity analyses are shown in fig. S19. The map was made with 500 optimizations.

**EMC (VeroE6)**

The ‘EMC (VeroE6)’ dataset was provided by Prof. Bart Haagmans and Anna Mykytyn. The antigenic map was originally published by Mykytyn et al., 2022 (*4*), as Figure 3A. The dataset used the same sera as the EMC (PRNT) dataset but titrations were done using a VSV pseudotype neutralization assay on VeroE6 cells. For the analyses presented here, we excluded the P.1, P.2, and B.1.621 sera from the map and titer analyses, as their homologous variants were not titrated. The map used here was constructed using 1000 optimisations. Sensitivity analyses of the map can be found in Mykytyn et al., 2022 (*4*). The full map is shown in fig. S20.

**EMC (Calu-3)**

The ‘EMC (Calu-3)’ dataset was provided by Prof. Bart Haagmans and Anna Mykytyn. The antigenic map was originally published by Mykytyn et al., 2022 (*4*), as Figure 3B. The dataset used the same sera as the EMC (PRNT) dataset but titrations were done using a VSV pseudotype neutralization assay on Calu-3 cells. For the analyses presented here, we excluded the P.1, P.2, and B.1.621 sera from the map and titer analyses, as their homologous variants were not titrated. The map used here was constructed using 1000 optimisations. Sensitivity analyses of the map can be found in Mykytyn et al., 2022 (*4*). The full map is shown in fig. S21.

**Galveston**

The ‘Galveston’ dataset was provided by Pei-Yong Shi at the University of Texas Medical Branch in Galveston, TX. The raw data was originally published in (*9*), and corresponds to the titrations done with hamster sera taken 28 days post infection (published in Supplementary Table 2 of (*9*)). The full map and sensitivity analyses are shown in fig. S22. The map was made with 1000 optimizations.

**Madison (FRNT)**

The ‘Madison (FRNT)’ dataset was provided by Prof. Yoshihiro Kawaoka and Prof. Peter Halfmann at the University of Wisconsin-Madison. The FRNT used to determine the neutralization titers is described in Takashita et al., 2022 (*37*). The full map and sensitivity analyses are shown in fig. S23. The map was made with 1000 optimizations.

**Maryland**

The ‘Maryland’ dataset was provided by Prof. Matthew Frieman at the University of Maryland and was originally published as Figure 6B and supplementary fig. S4 in (*26*). The sera used in this map were pooled sera from 20 BALB/c mice. The full map and sensitivity analyses are shown in fig. S24. The map was made with 1000 optimizations.

**WUSTL**

The Washington University in St. Louis (WUSTL) dataset was provided by Michael Diamond at Washington University School of Medicine. The antigenic map was originally published in Ying et al., 2021 (*10*). The full map and sensitivity analyses are shown in fig. S25. The map was made with 1000 optimizations.

#### Titer analyses

GMTs accounting for non-detectable titers were inferred using the “titertools” package in R (*38*), using the “HDI” option for estimation of the GMT and 95% highest posterior density intervals. The prior for the mean of the GMT was a normal distribution with mean 0 and standard deviation 100 and the prior for the standard deviation of the GMT was an inverse gamma distribution with a shape parameter of 2 and a scale parameter of 0.75.

Differences in titer magnitude between datasets were modeled such that the logged titer measured (here and elsewhere log always refers to $\log_2$) for variant i and serum j in dataset m is given by:

$$logtiter_{ijm} = serumGroupGMT_{iJ} + serumEffect_j + datasetMagnitudeEffect_m + \varepsilon_{ij}$$

Equation 1.

Where serumGroupGMT corresponds to the average logged titer of antigen i against serum group J, serumEffect is the serum reactivity bias of serum j and datasetMagnitudeEffect is the dataset magnitude effect of dataset m. ε is independently and normally distributed $\log_2$ titer noise which is assumed to have a standard deviation that is separately estimated for each dataset. Priors for this standard deviation parameters were calculated from an inverse gamma distribution with shape=3, and scale=1.5. The following prior distributions were used: serumGroupGMT: N(7, 20). serumEffect: N(0, 6). datasetMagnitudeEffect: N(0, 6). Raw titers were adjusted by the estimated dataset reactivity effect. The adjusted titers were used for the analyses of fold change and immunodominance effects.

Differences in titer magnitude between animal model sera and assay was modeled such that the logged titer measured for variant i and serum j in dataset m is given by:

$$logtiter_{ijm} = serumGroupGMT_{iJ} + serumEffect_j + animalMagnitudeEffect_a + assayMagnitudeEffect_b + \varepsilon_{ij}$$

Equation 2.

Where serumGroupGMT corresponds to the average logged titer of antigen i against serum group J, serumEffect is the serum reactivity bias of serum j, animalMagnitudeEffect is the magnitude effect for the animal model a (one of human, hamster or mouse), assayMagnitudeEffect is the magnitude effect of the assay b (one of FRNT, LV-PV-Neut, VSV-PV-Neut, Microneut, PRNT or CPE), and ε is independently and normally distributed log titer noise. The following prior distributions were used: serumGroupGMT: N(7, 20). serumEffect: N(0, 6). animalMagnitudeEffect: N(0, 6). assayMagnitudeEffect: N(0, 6).

In addition to evaluating the effect of animal model and assay on titer magnitude, we also considered whether the neutralization cut-off may affect titer magnitude, as six datasets used a neutralization cut-off other than 50% reduction of plaques. We found no influence of neutralization cut-off on titer magnitude (fig. S29D,E).

Differences in variability between datasets after adjusting for dataset magnitude were estimated by introducing an additional term into equation 1 to account for the extent to which the serum group mean against each antigen deviated for each dataset from the overall dataset average, with individual log titers described as:

$$logtiter_{ijm} = serumGroupGMT_{iJ} + serumEffect_j + datasetMagnitudeEffect_m + serumGroupGMTResidualDeviation_m$$

Equation 3.

In this formulation all serumGroupGMTResidualDeviation parameters were drawn from a normal distribution with mean 0 and a standard deviation parameter that was separately estimated for each dataset, which forms the “dataset variability effect” shown in fig. S33. The following additional prior distributions were used: serumGroupGMT: N(7, 20). serumEffect: N(0, 6). datasetMagnitudeEffect: N(0, 6).

Fitting was performed using the “cmdstanr” interface to Stan.

### Fold change modeling

Fold change from homologous variants accounting for non-detectable titers was inferred using the “titertools” package in R (*38*), using the “HDI” option, as described in Wilks et al., 2022 (*3*). The prior for the mean of the fold change was a normal distribution with mean 0 and standard deviation 100 and the prior for the standard deviation of the fold change was an inverse gamma distribution with a shape parameter of 2 and a scale parameter of 0.75.

We estimated how the magnitude of the fold change varies by dataset. The fold change from homologous measured for variant i and serum j in serum group J in dataset m is given by:

$$predictedFoldchange_{ijm} = agFoldDrops_{iJ} * datasetSlope_{m} + \varepsilon_{ij}$$

Equation 4.

Where agFoldDrops correspond to the average fold change from the homologous variant measured for antigen i in serum group J across all datasets, and the datasetSlope is the slope specific to each dataset, m. ε is independently and normally distributed log titer noise. The following prior distributions were used: datasetSlope: N(1, 10). agFoldDrops: N(-1, 3).

We also estimated the effect of the animal model and the assay used on the magnitude of the fold change. In that case, the logged titer measured for variant i and serum j in serum group J in dataset m is given by:

$$predictedFoldchange_{ijm} = agFoldDrops_{iJ} * animalSlope_{a} * assaySlope_{b} + \varepsilon_{ij}$$

Equation 5.

Where agFoldDrops correspond to the average fold change measured for antigen i in serum group J across all datasets, and the animalSlopeEffect and assaySlopeEffect are the slope specific to each animal model and each assay. ε is independently and normally distributed log titer noise. The following prior distributions were used: assaySlope: N(1, 10). animalSlope: N(1, 10). agFoldDrops: N(-1, 3).

Fitting was performed using the “cmdstanr” interface to Stan.

#### Antigenic cartography

Antigenic maps were constructed using the “Racmacs” package (*39*) in R using 1000 optimizations and the minimum column basis parameter set to “none”. All maps were evaluated for fit in two dimensions and correspondence of map distances and table distances when not already performed in the original publication. For assessing dimensionality of each individual map, the root-mean-squared-error (RMSE) of detectable titers in 1 to 4 dimensions was compared to known titers. Per dimension, 1000 repeats were performed wherein a map was constructed from 90% of all titers. For each run, the RMSE is calculated from the differences between the titers predicted from the map and the known titers on the $\log_2$ scale. As the orientation of the map in the x and y direction is free, maps were oriented on the x-axis such that the line from the ancestral variant (either D614G or 614D) to Omicron BA.1 is horizontal, with Omicron BA.1 on the right, and on the y-axis such that B.1.351 is at the top. Maps without Omicron BA.1 were oriented such that the ancestral variant (D614G or 614D) is on the left, the B.1.351 variant at the top and the B.1.617.2 variant at the bottom.

The merged antigenic map was constructed by merging on a per-variant basis, where we considered the variants to be equal across datasets, and each serum in each dataset was considered individually. The merged map was constructed using 5000 optimizations, with the minimum column basis parameter set to “none”.

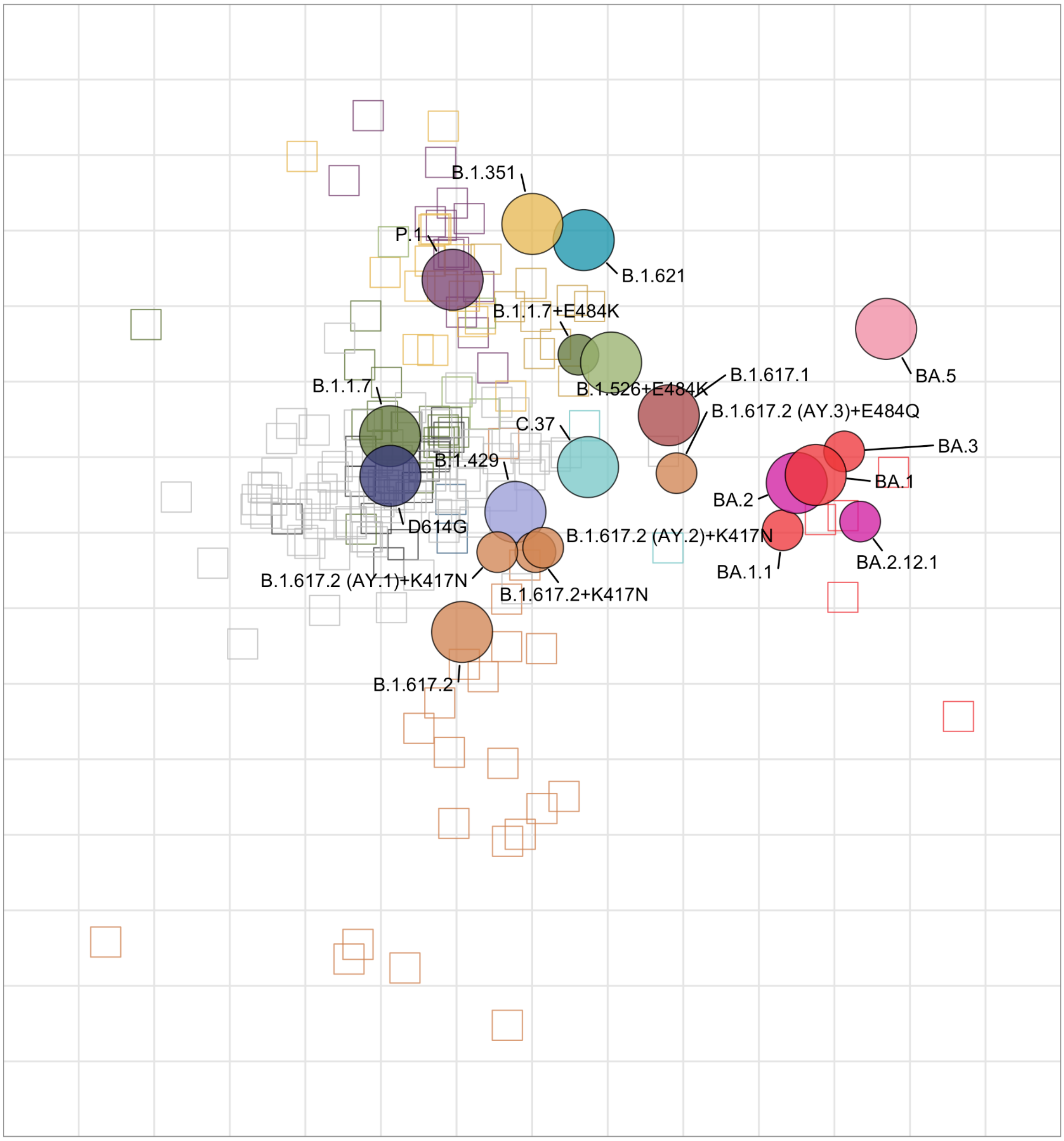


**Figure S1: Detailed figure of the Duke antigenic map.** Sensitivity analyses can be found in Wilks et al., 2022 (*3*).

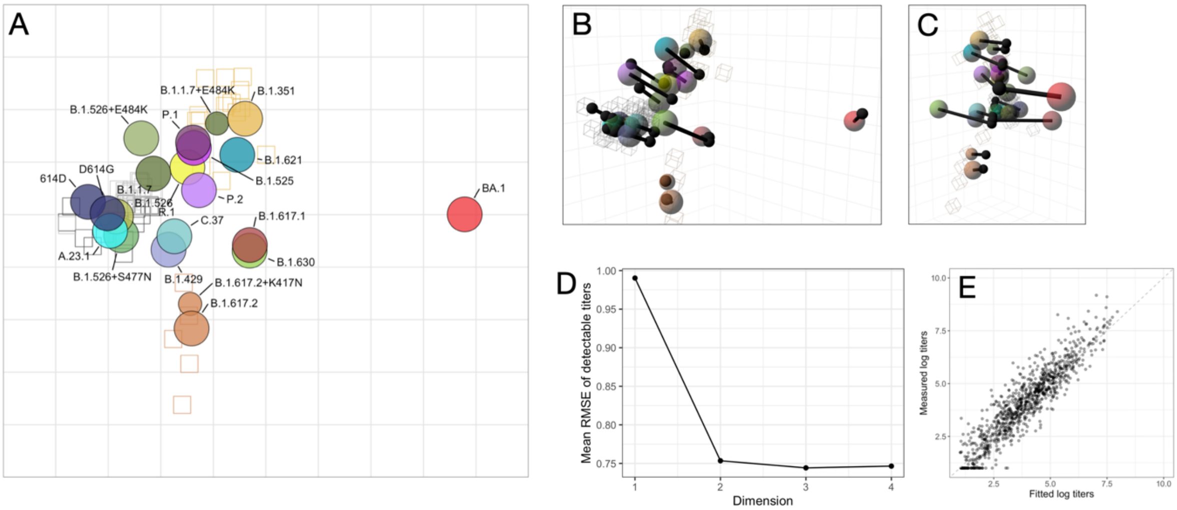


**Figure S2: Emory antigenic map.** A) Detailed labeled figure of the Emory antigenic map. B) Emory antigenic map optimized in three dimensions, 'side' view. C) Emory antigenic map optimized in three dimensions, 'front' view. D) Dimensionality test for Emory antigenic map. Three-dimensional maps were constructed with 500 optimisations. Dimensionality tests were performed with 500 replicates per dimension and 500 optimisations. E) Scatter plot of fitted titers and measured titers. The fitted titers are determined from the distances in the antigenic map. Titers are plotted on the $\log_2$ scale. The dashed gray line is the line of best fit. Non-detectable measured titers were set to 1. In B and C, the black lines point to the position of the variant in the two dimensional map. All analyses were performed after removing outlier sera.

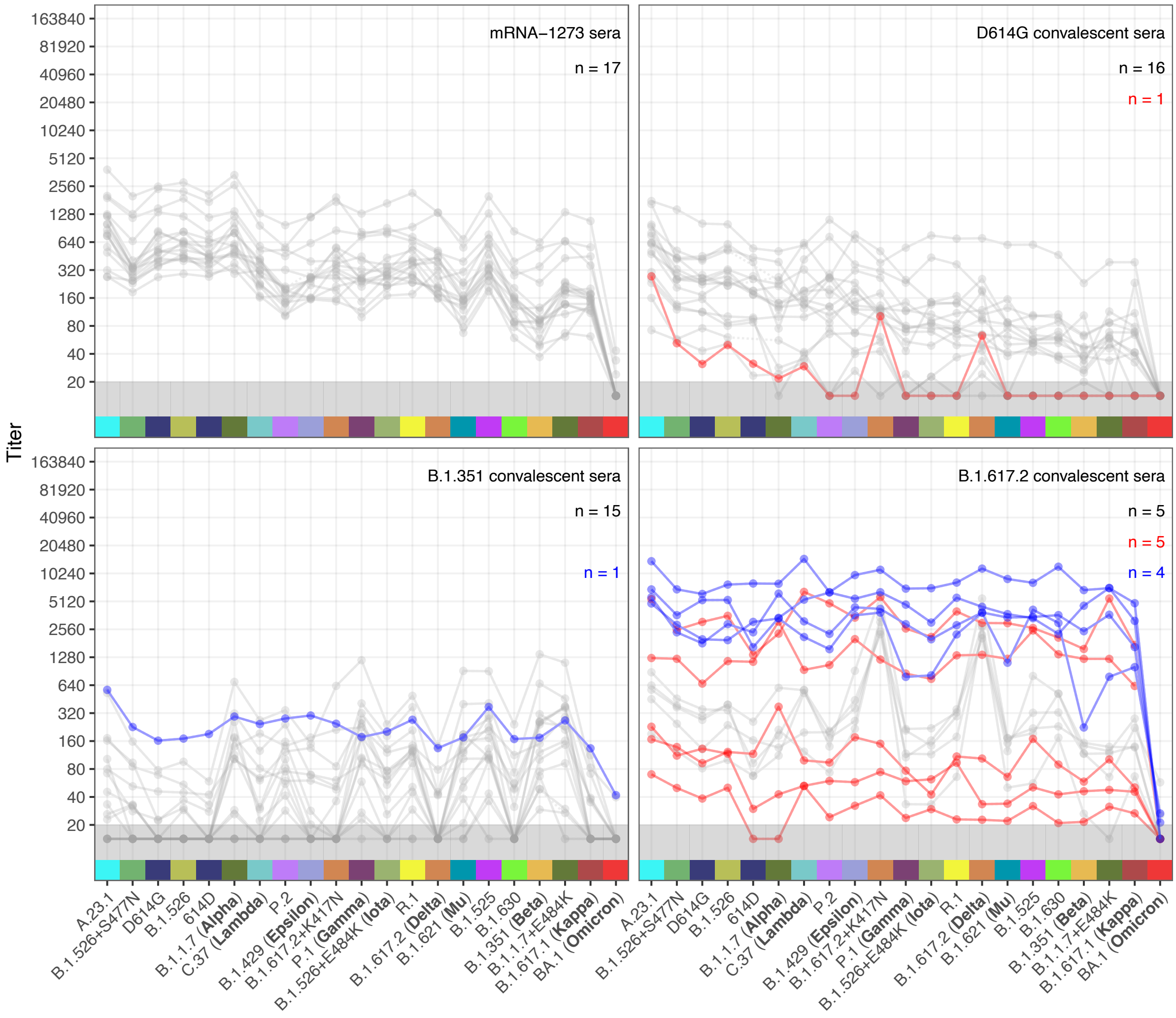


**Figure S3: Titerplots for the Emory dataset, with outlier titers highlighted.** Sera excluded based on the presence of a titer that was >2-fold higher than the titer against the homologous variant for their serum group are shown in red, with the number of excluded sera given in red numbers at the top right. The A.23.1 and B.1.525 variants were not considered when determining the titers >2-fold higher than homologous. Sera excluded based on visual inspection of titer patterns are highlighted in blue, with the number of excluded sera given in red in the top right. Total number of non-excluded sera are shown in black in the top right. Points in the gray region at the bottom of the plots show titers and GMTs that fell below the detection threshold of 20.

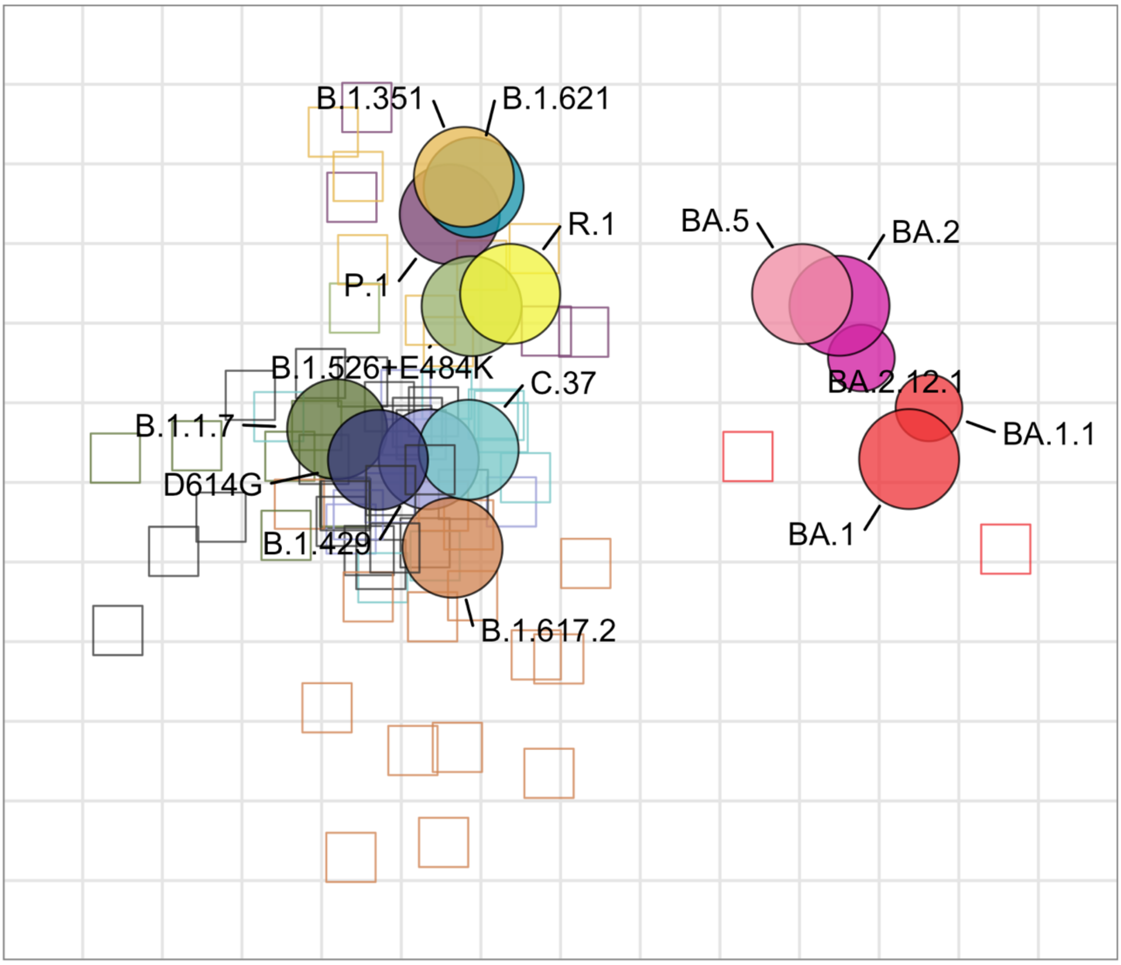


**Figure S4: Detailed figure of the FDA antigenic map.** Sensitivity analyses can be found in Wang et al., 2022 (*8*). The map is shown after removing outlier sera.

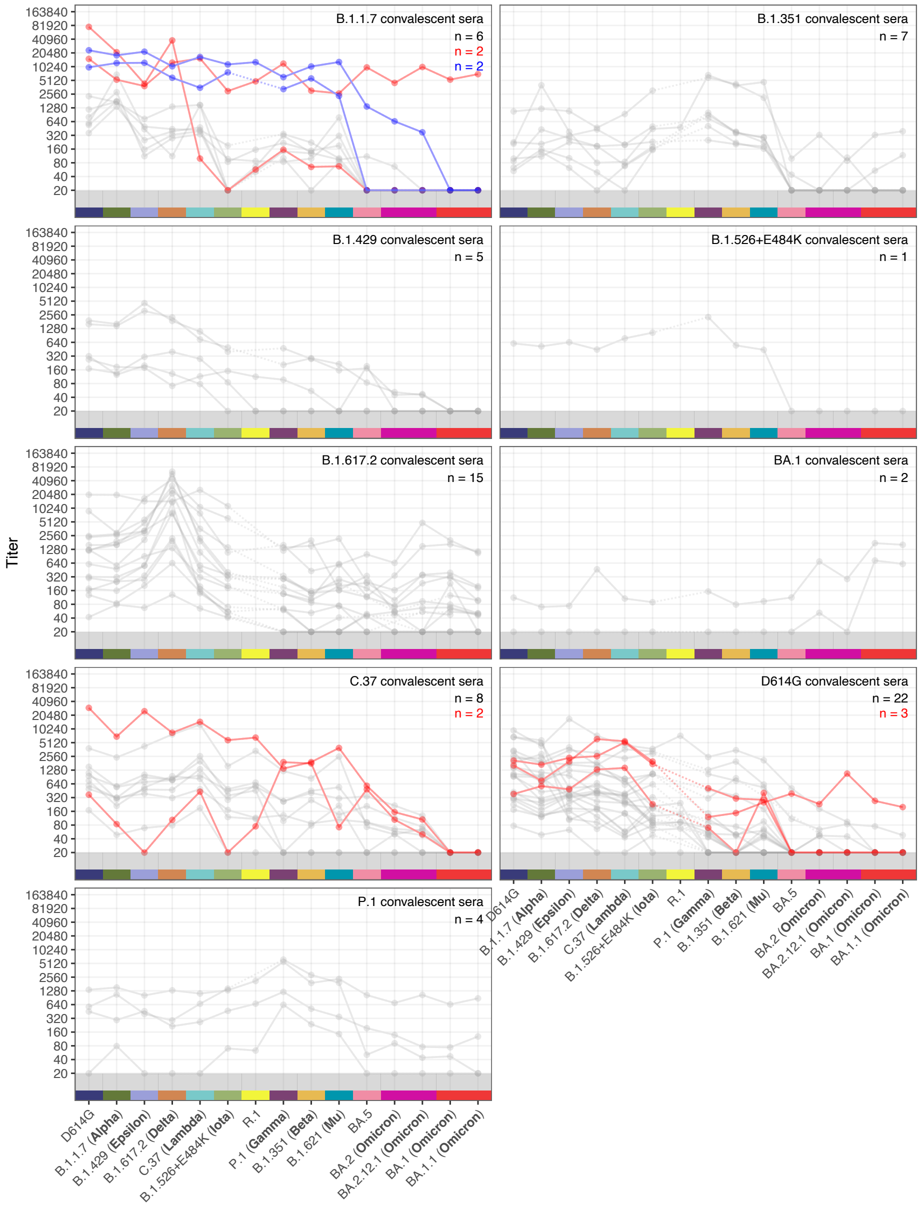


**Figure S5: Titerplots for the FDA dataset, with outlier titers highlighted.** Sera excluded based on the presence of a titer that was >2-fold higher than the titer against the homologous variant for their serum group are shown in red, with the number of excluded sera given in red numbers at the top right. The P.1 variant was not considered when determining the titers >2-fold higher than homologous. Sera excluded based on visual inspection of titer patterns are highlighted in blue, with the number of excluded sera given in red in the top right. Total number of non-excluded sera are shown in black in the top right. Points in the gray region at the bottom of the plots show titers and GMTs that fell below the detection threshold of 20.

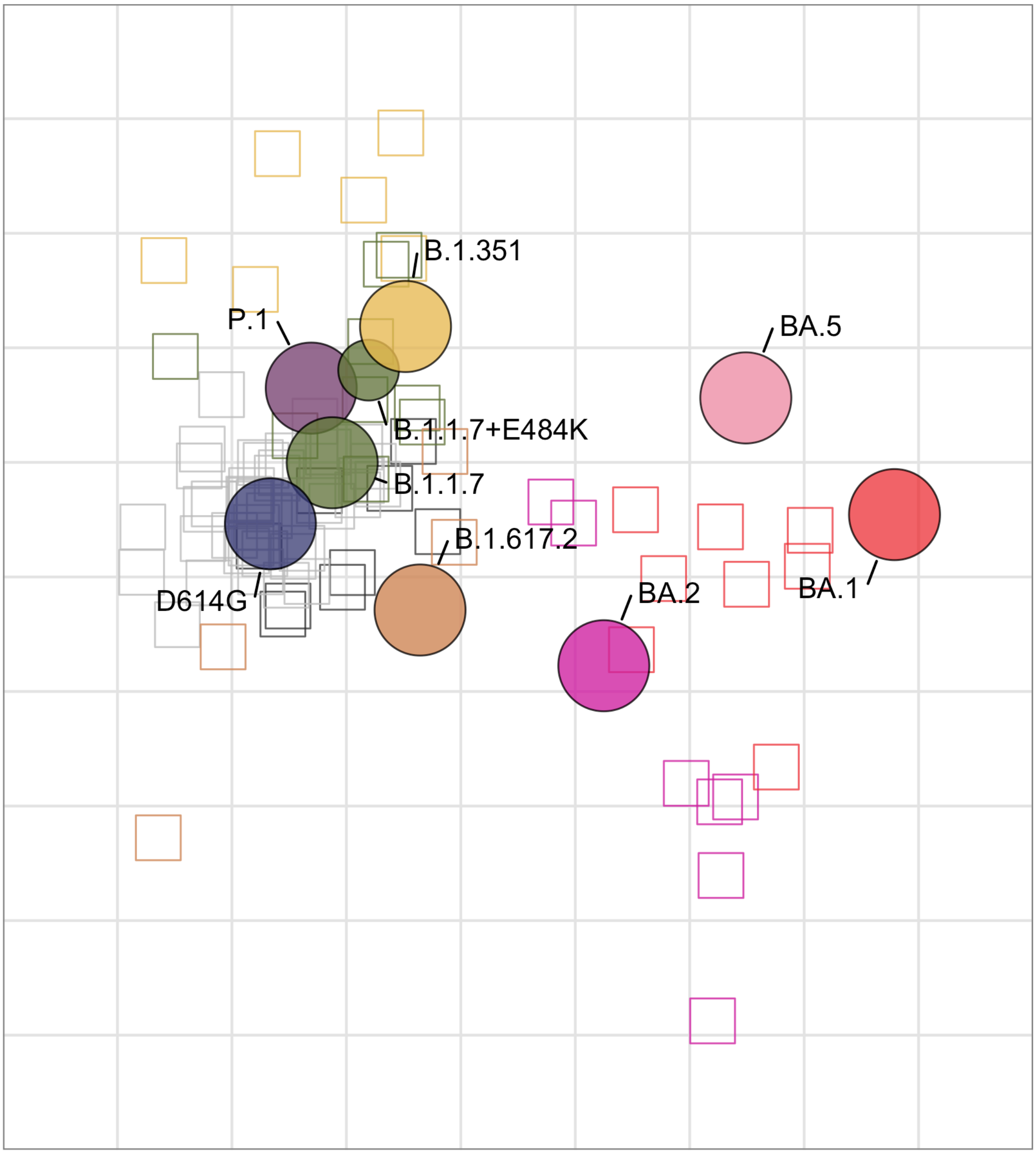


**Figure S6: Detailed figure of the Innsbruck antigenic map.** Sensitivity analyses can be found in Rössler et al., 2023 (*12*). The map is shown after removing outlier sera.

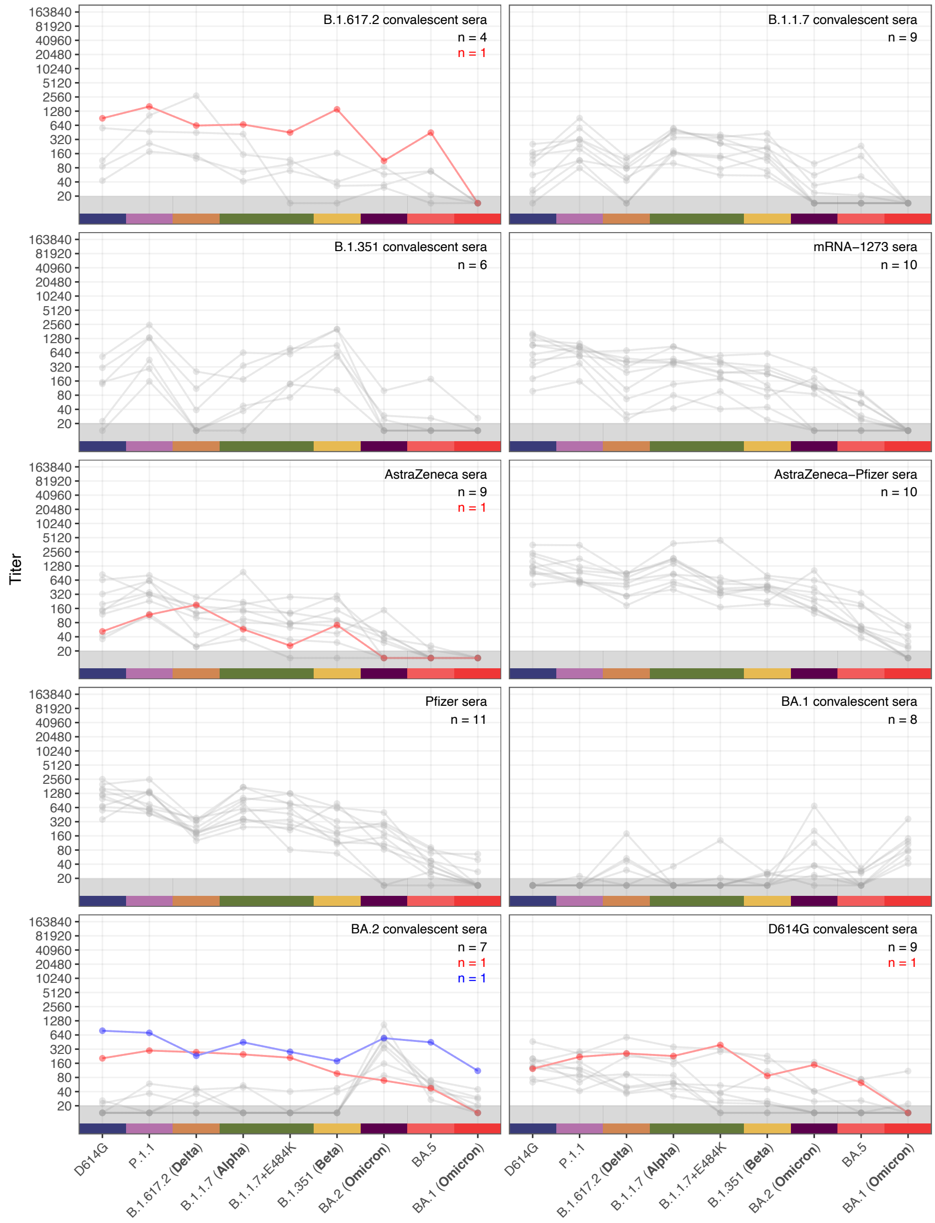


**Figure S7: Titerplots for the Innsbruck dataset, with outlier titers highlighted.** Sera excluded based on the presence of a titer that was >2-fold higher than the titer against the homologous variant for their serum group are shown in red, with the number of excluded sera given in red numbers at the top right. The P.1 and BA.2 variants were not considered when determining the titers >2-fold higher than homologous. Sera excluded based on visual inspection of titer patterns are highlighted in blue, with the number of excluded sera given in red in the top right. Total number of non-excluded sera are shown in black in the top right. Points in the gray region at the bottom of the plots show titers and GMTs that fell below the detection threshold of 20.

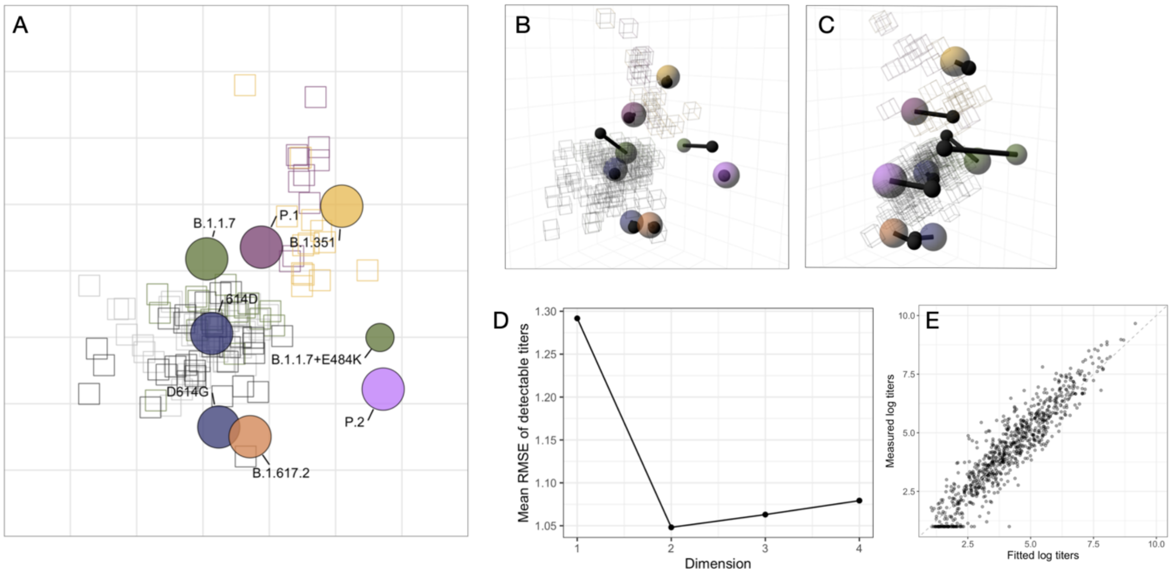


**Figure S8: Oxford antigenic map.** A) Detailed labeled figure of the Oxford antigenic map. B) Oxford antigenic map optimized in three dimensions, 'side' view. C) Oxford antigenic map optimized in three dimensions, 'front' view. D) Dimensionality test for 'Oxford' antigenic map. Three-dimensional maps were constructed with 500 optimisations. Dimensionality tests were performed with 500 replicates per dimension and 500 optimisations. E) Scatter plot of fitted titers and measured titers. The fitted titers are determined from the distances in the antigenic map. Titers are plotted on the $\log_2$ scale. The dashed gray line is the line of best fit. Non-detectable measured titers were set to 1. In B and C, the black lines point to the position of the variant in the two dimensional map. All analyses were performed after removing outlier sera.

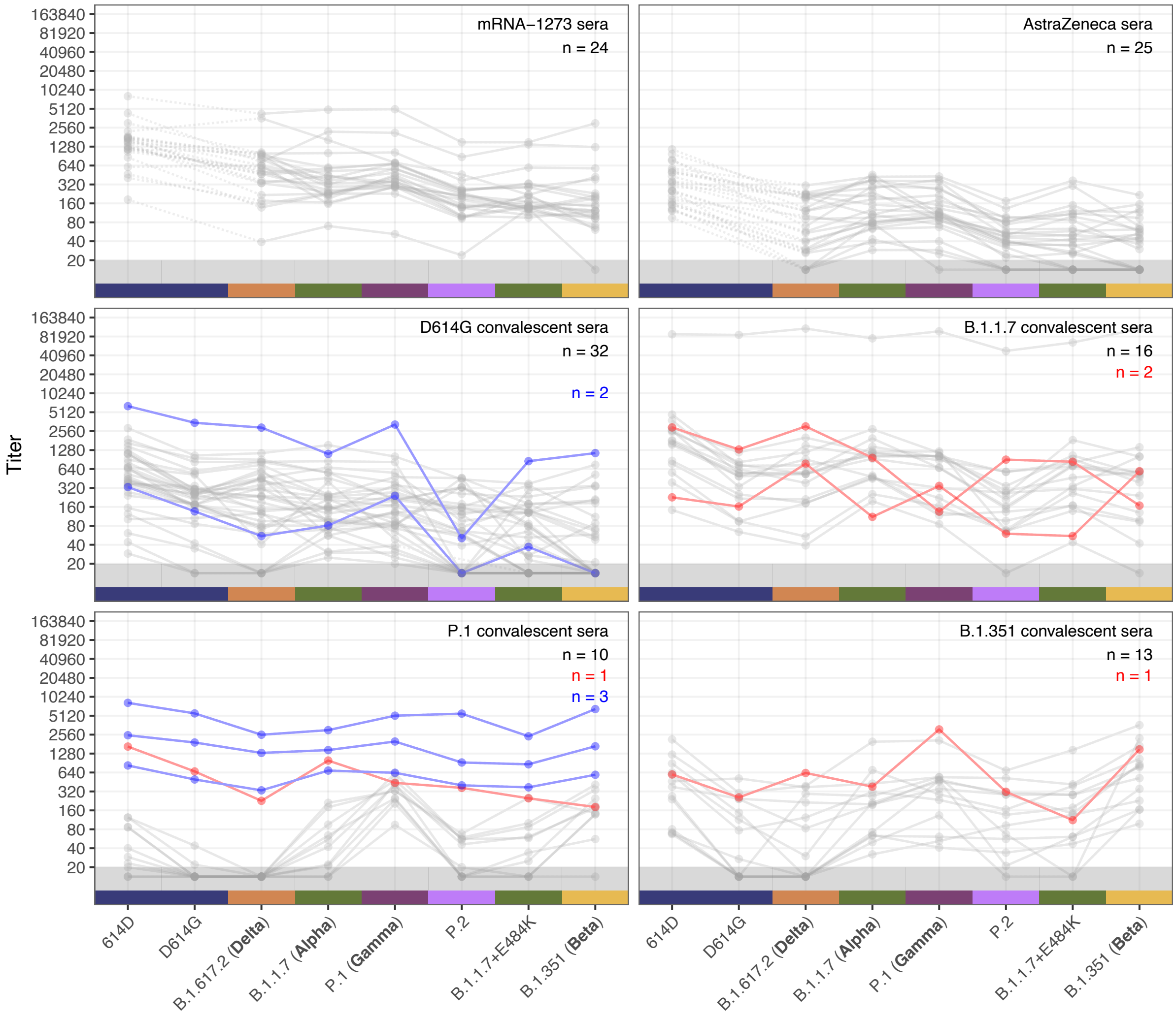


**Figure S9: Titerplots for the Oxford dataset, with outlier titers highlighted.** Sera excluded based on the presence of a titer that was >2-fold higher than the titer against the homologous variant for their serum group are shown in red, with the number of excluded sera given in red numbers at the top right. The 614D variant was not considered when determining the titers >2-fold higher than homologous. Sera excluded based on visual inspection of titer patterns are highlighted in blue, with the number of excluded sera given in red in the top right. Total number of non-excluded sera are shown in black in the top right. Points in the gray region at the bottom of the plots show titers and GMTs that fell below the detection threshold of 20.

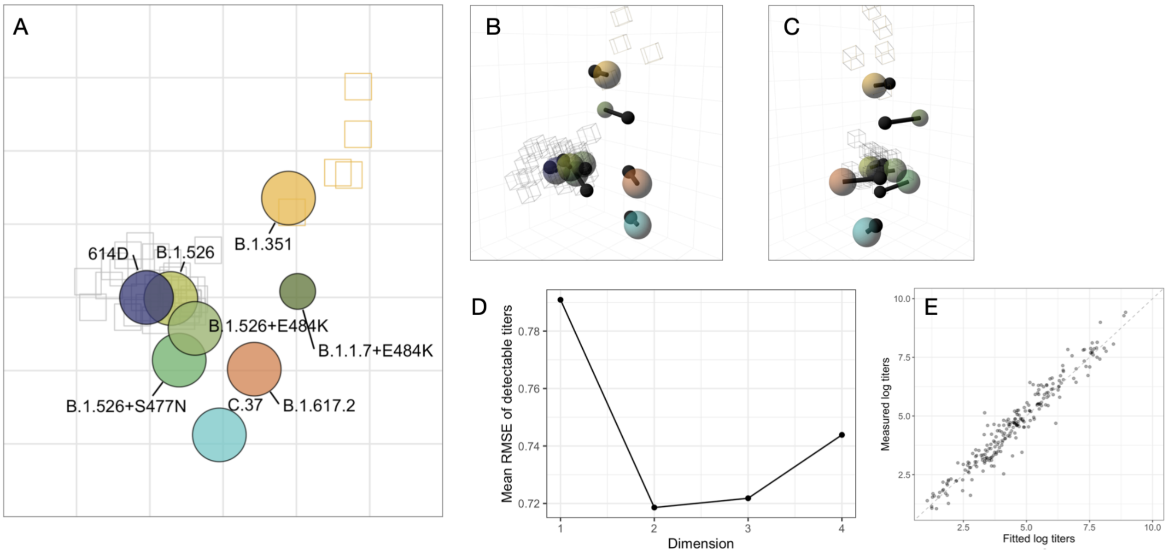


**Figure S10: Mt. Sinai antigenic map.** A) Detailed labeled figure of the Mt. Sinai antigenic map. B) Mt. Sinai antigenic map optimized in three dimensions, 'side' view. C) Mt. Sinai antigenic map optimized in three dimensions, 'front' view. D) Dimensionality test for Mt. Sinai antigenic map. Three-dimensional maps were constructed with 500 optimisations. Dimensionality tests were performed with 500 replicates per dimension and 500 optimisations. E) Scatter plot of fitted titers and measured titers. The fitted titers are determined from the distances in the antigenic map. Titers are plotted on the $\log_2$ scale. The dashed gray line is the line of best fit. Non-detectable measured titers were set to 1. In B and C, the black lines point to the position of the variant in the two dimensional map. All analyses were performed after removing outlier sera.

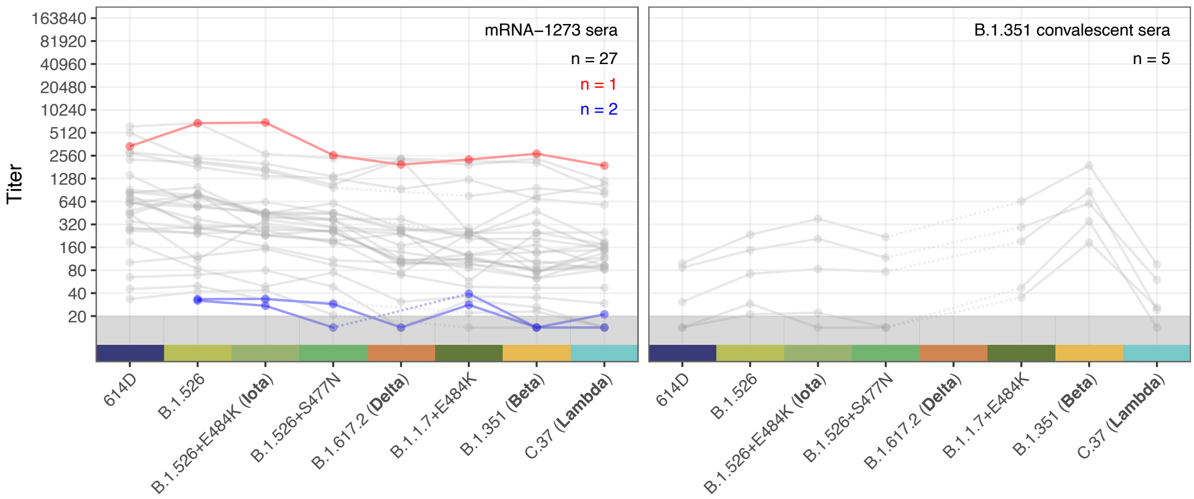


**Figure S11: Titerplots for the Mt. Sinai dataset, with outlier titers highlighted.** Sera excluded based on the presence of a titer that was >2-fold higher than the titer against the homologous variant for their serum group are shown in red, with the number of excluded sera given in red numbers at the top right. Sera excluded based on visual inspection of titer patterns are highlighted in blue, with the number of excluded sera given in red in the top right. Total number of non-excluded sera are shown in black in the top right. Points in the gray region at the bottom of the plots show titers and GMTs that fell below the detection threshold of 20.

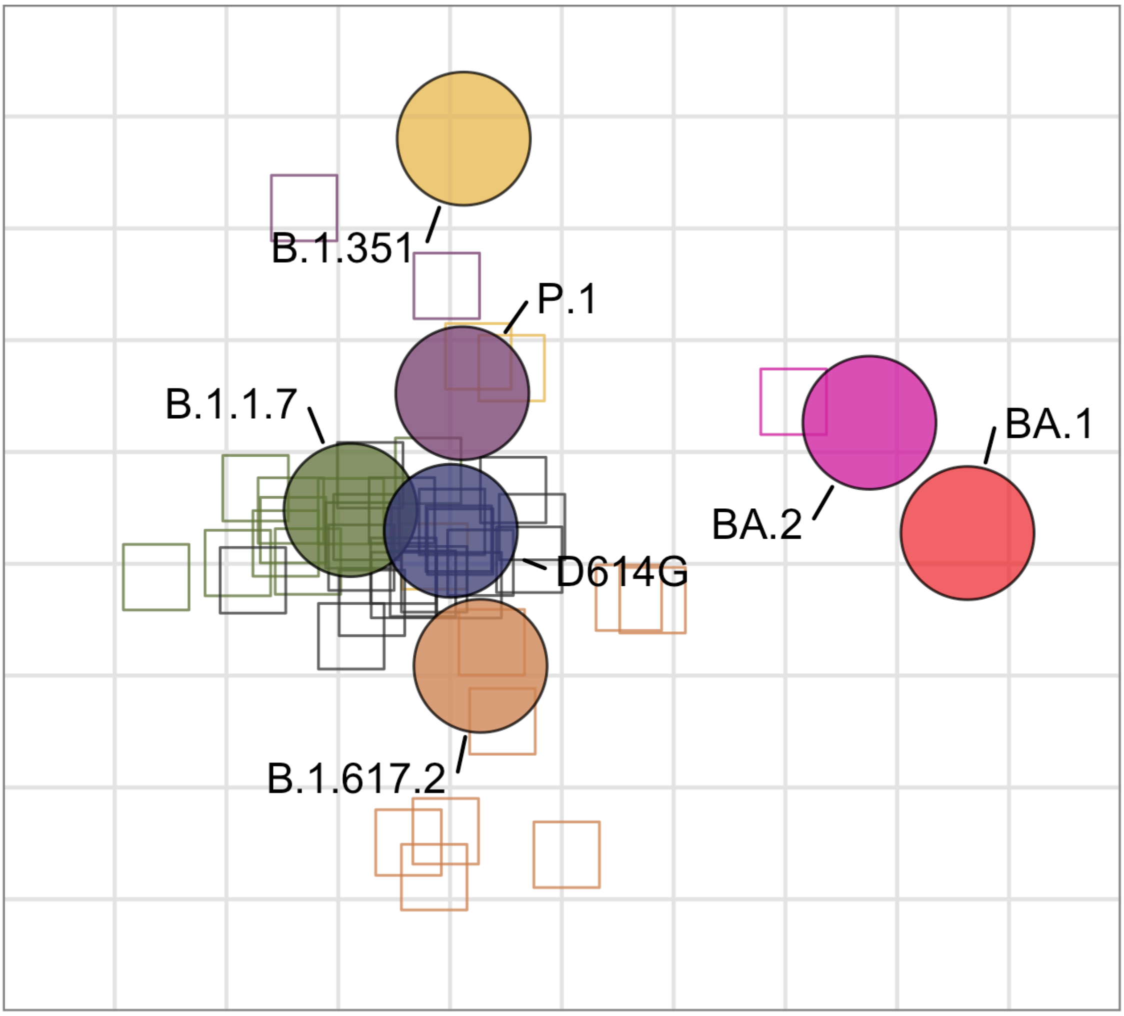


**Figure S12: Detailed figure of the AMC antigenic map.** Sensitivity analyses can be found in van der Straten et al., 2022 (*11*). The map is shown after removing outlier sera.

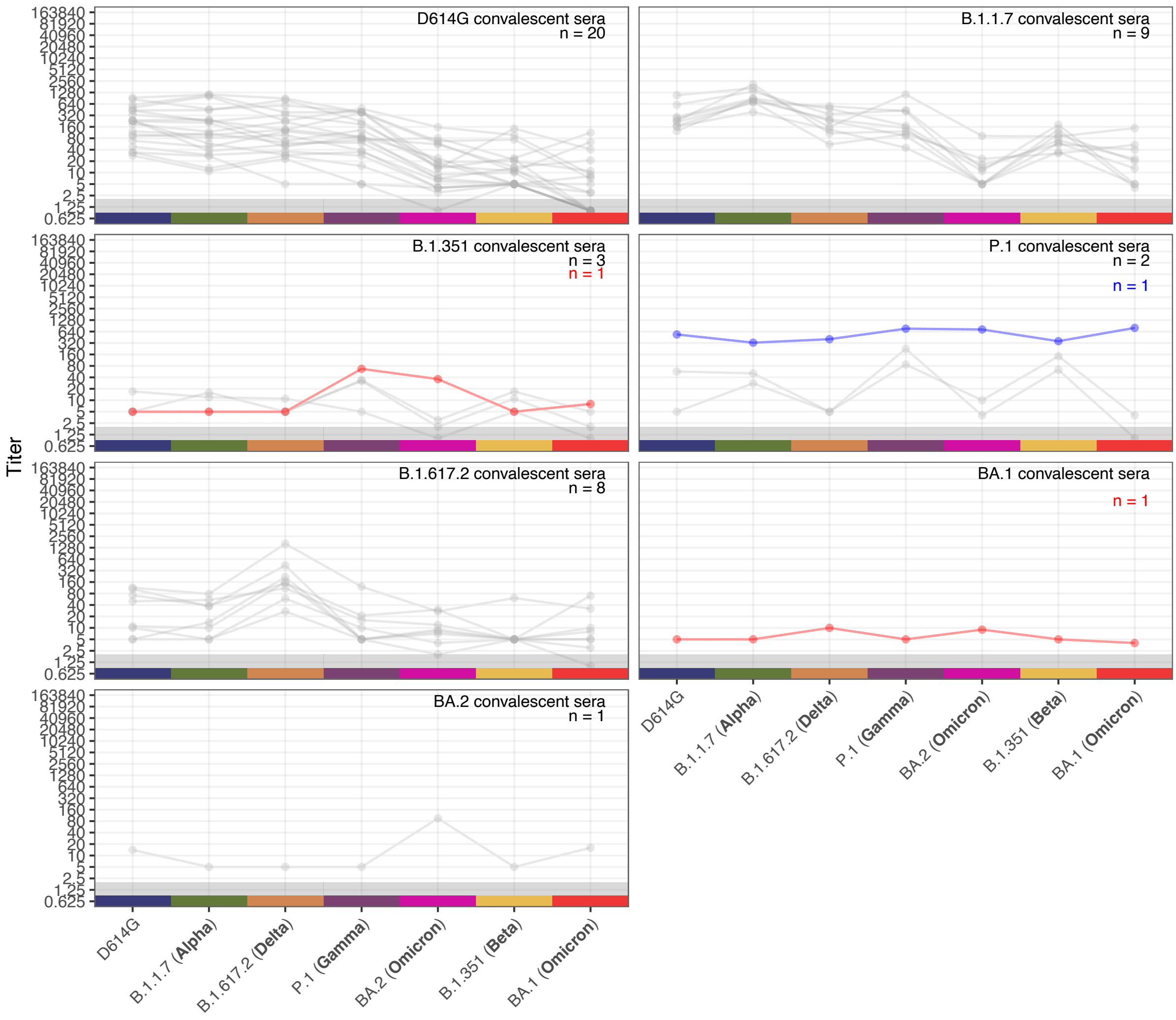


**Figure S13: Titerplots for the AMC dataset, with outlier titers highlighted.** Sera excluded based on the presence of a titer that was >2-fold higher than the titer against the homologous variant for their serum group are shown in red, with the number of excluded sera given in red numbers at the top right. The P.1 variant was not considered when determining the titers >2-fold higher than homologous. Sera excluded based on visual inspection of titer patterns are highlighted in blue, with the number of excluded sera given in red in the top right. Total number of non-excluded sera are shown in black in the top right. Points in the gray region at the bottom of the plots show titers and GMTs that fell below the detection threshold of 2.

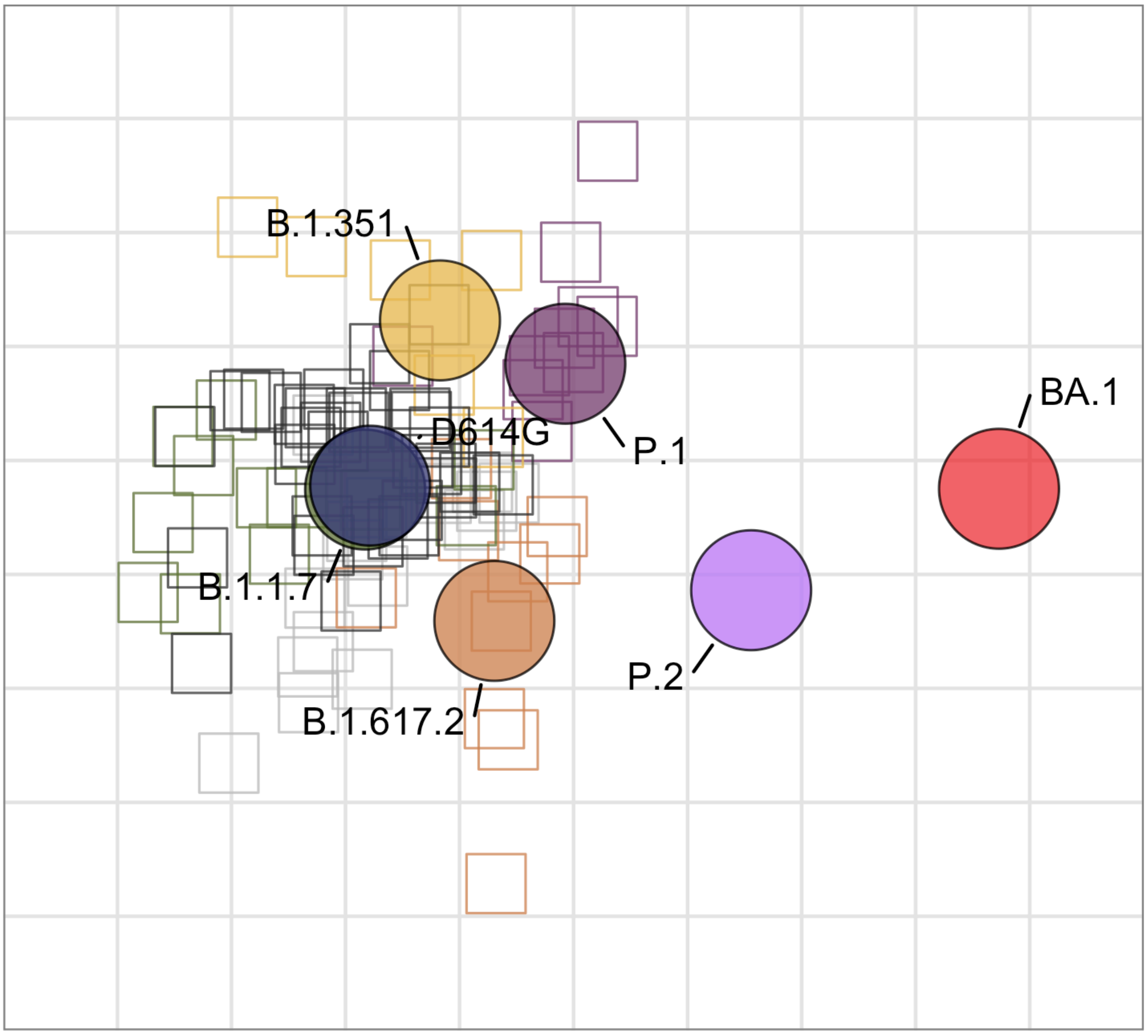


**Figure S14: Detailed figure of the Geneva antigenic map.** Sensitivity analyses can be found in Bekliz et al., 2022 (*7*).

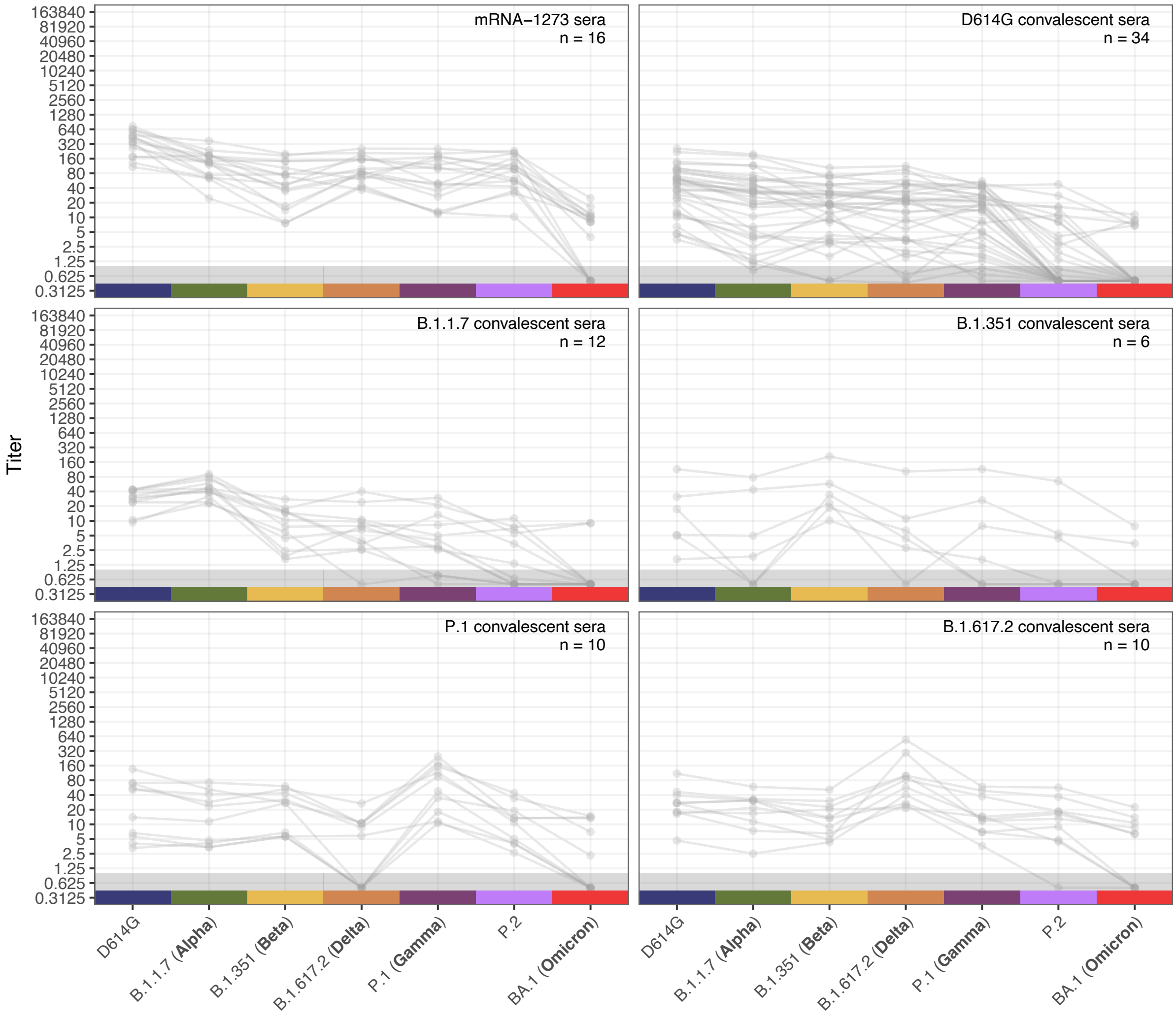


**Figure S15: Titerplots for the Geneva dataset.** No outlier sera were identified based on the presence of a titer that was >2-fold higher than the titer against the homologous variant for their serum group or based on visual inspection of titer patterns. Total number of sera is shown in black in the top right. Points in the gray region at the bottom of the plots show titers and GMTs that fell below the detection threshold of 1.

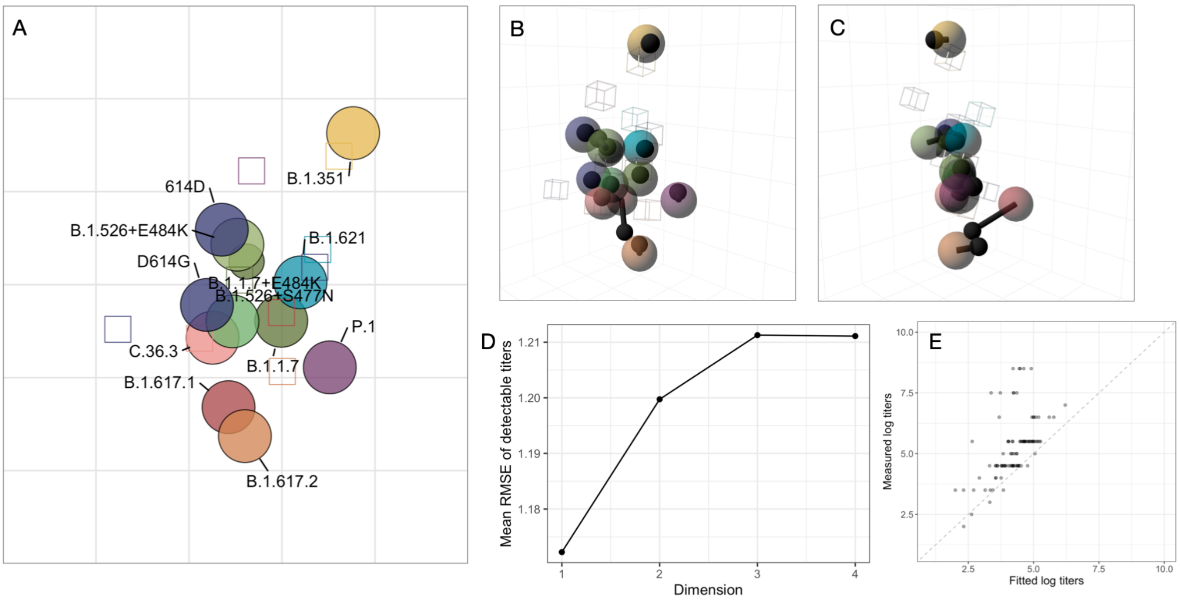


**Figure S16: Madison (pooled) antigenic map.** A) Detailed labeled figure of the Madison (pooled) antigenic map. B) Madison (pooled) antigenic map optimized in three dimensions, ‘side’ view. C) Madison (pooled) antigenic map optimized in three dimensions, ‘front’ view. D) Dimensionality test for the Madison (pooled) antigenic map. Three-dimensional maps were constructed with 500 optimisations. Dimensionality tests were performed with 500 replicates per dimension and 500 optimisations. E) Scatter plot of fitted titers and measured titers. The fitted titers are determined from the distances in the antigenic map. Titers are plotted on the $\log_2$ scale. The dashed gray line is the line of best fit. Non-detectable measured titers were set to 1. In B and C, the black lines point to the position of the variant in the two dimensional map.

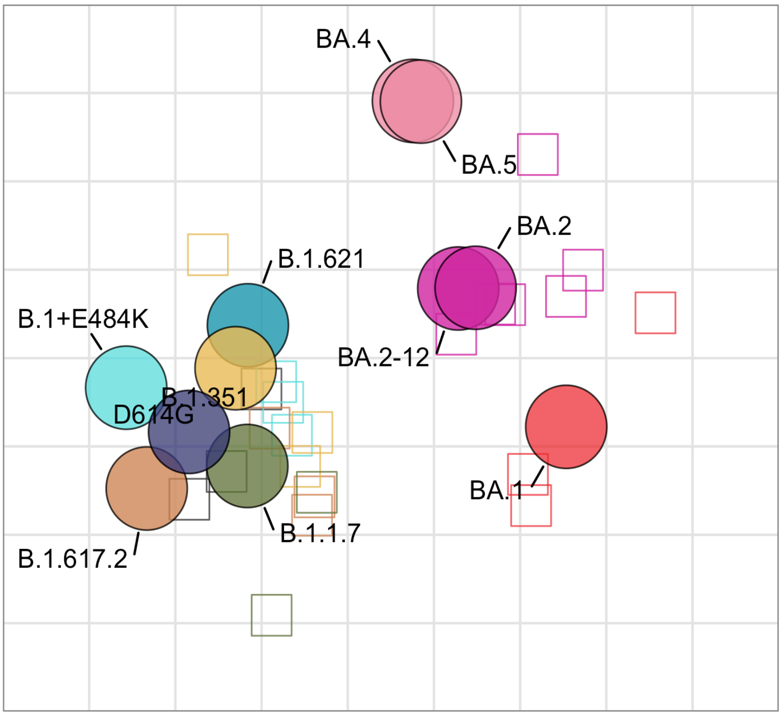


**Figure S17: Detailed figure of the Charité antigenic map.** Sensitivity analyses can be found in Mühlemann et al., 2023 (*25*).

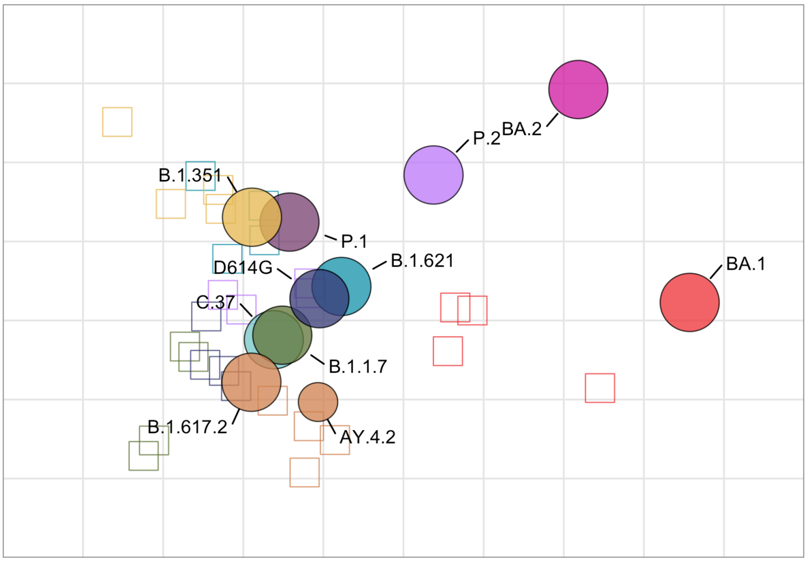


**Figure S18: Detailed figure of the EMC (PRNT) antigenic map.** Sensitivity analyses can be found in Mykytyn et al., 2022 (*4*).

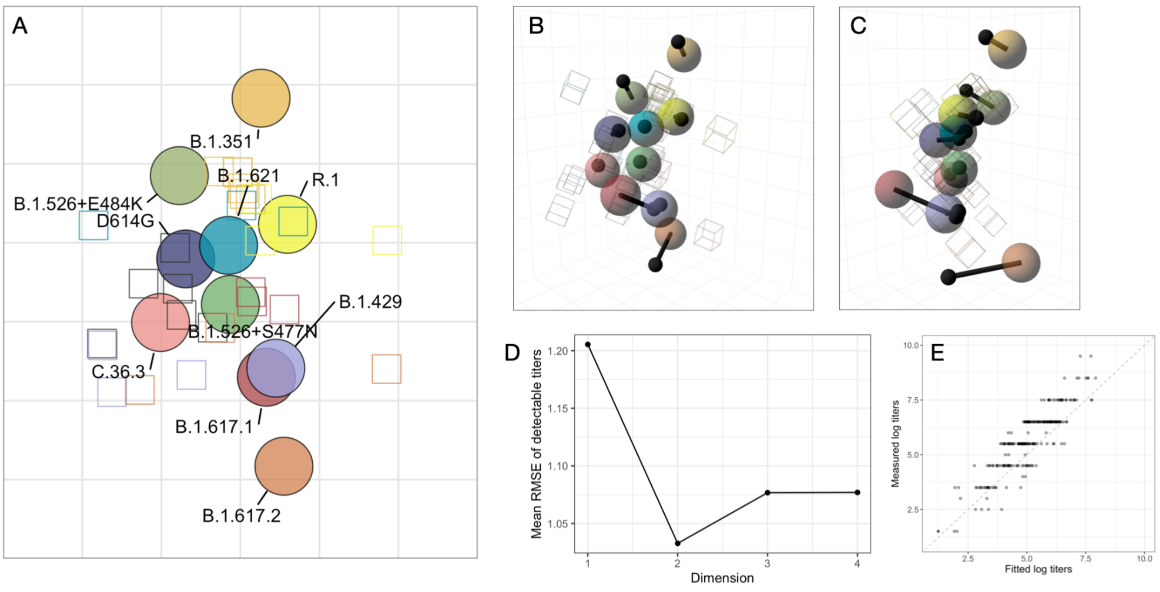


**Figure S19: Madison (unpooled) antigenic map.** A) Detailed labeled figure of the Madison (unpooled) antigenic map. B) Madison (unpooled) antigenic map optimized in three dimensions, 'side' view. C) Madison (unpooled) antigenic map optimized in three dimensions, 'front' view. D) Dimensionality test for Madison (unpooled) antigenic map. Three-dimensional maps were constructed with 500 optimisations. Dimensionality tests were performed with 500 replicates per dimension and 500 optimisations. E) Scatter plot of fitted titers and measured titers. The fitted titers are determined from the distances in the antigenic map. Titers are plotted on the $\log_2$ scale. The dashed gray line is the line of best fit. Non-detectable measured titers were set to 1. In B and C, the black lines point to the position of the variant in the two dimensional map.

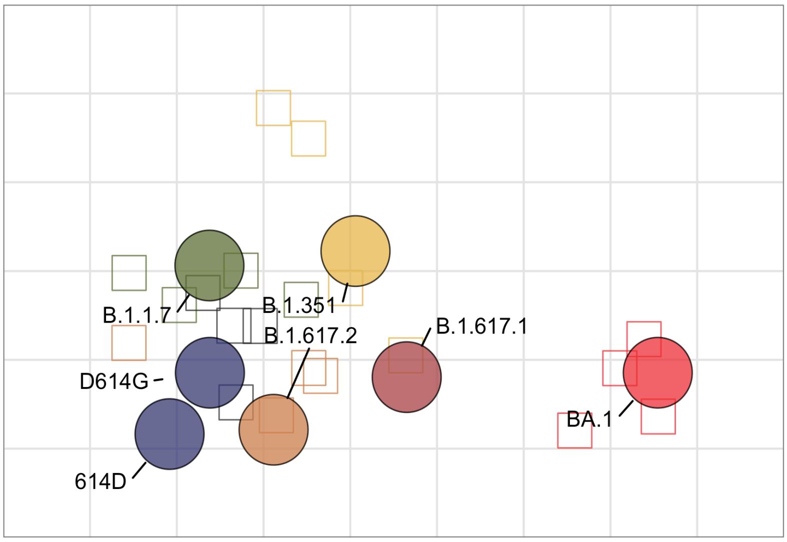


**Figure S20: Detailed figure of the EMC (VeroE6) antigenic map.** Sensitivity analyses can be found in Mykytyn et al., 2022 (*4*).

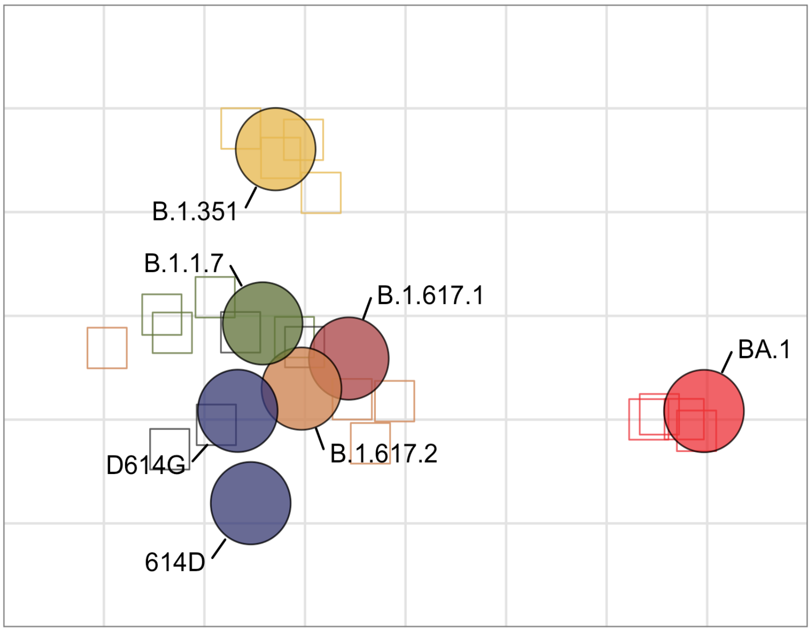


**Figure S21: Detailed figure of the EMC (Calu-3) antigenic map.** Sensitivity analyses can be found in Mykytyn et al., 2022 (*4*).

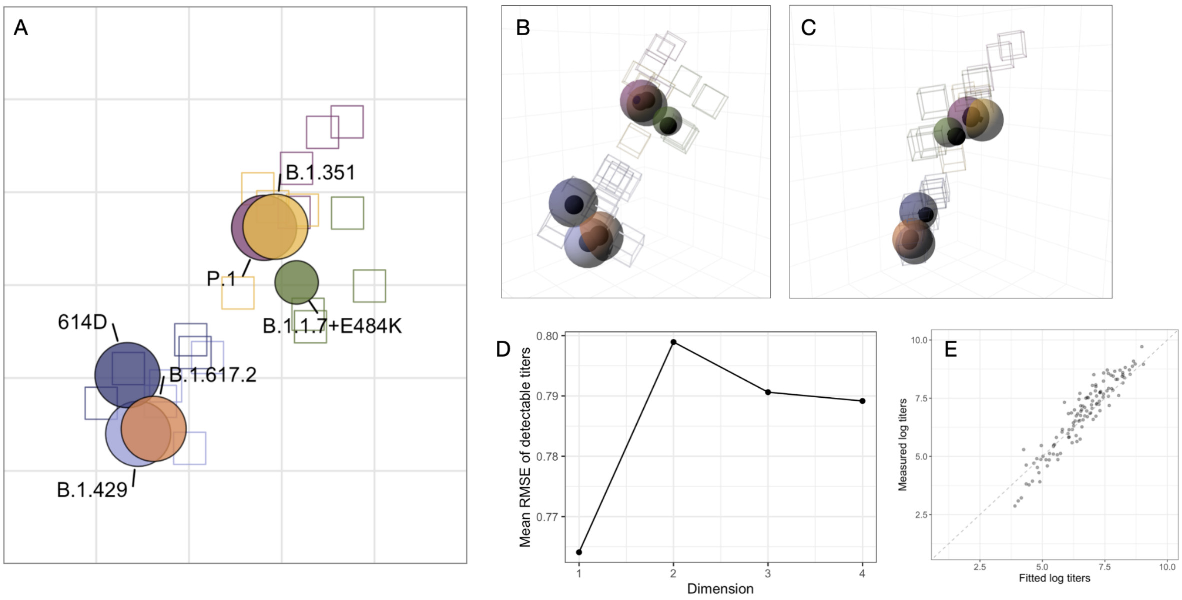


**Figure S22: Galveston antigenic map.** A) Detailed labeled figure of the Galveston antigenic map. B) Galveston antigenic map optimized in three dimensions, 'side' view. C) Galveston antigenic map optimized in three dimensions, 'front' view. D) Dimensionality test for Galveston antigenic map. Three-dimensional maps were constructed with 500 optimisations. Dimensionality tests were performed with 500 replicates per dimension and 500 optimisations. E) Scatter plot of fitted titers and measured titers. The fitted titers are determined from the distances in the antigenic map. Titers are plotted on the $\log_2$ scale. The dashed gray line is the line of best fit. Non-detectable measured titers were set to 1. In B and C, the black lines point to the position of the variant in the two dimensional map.

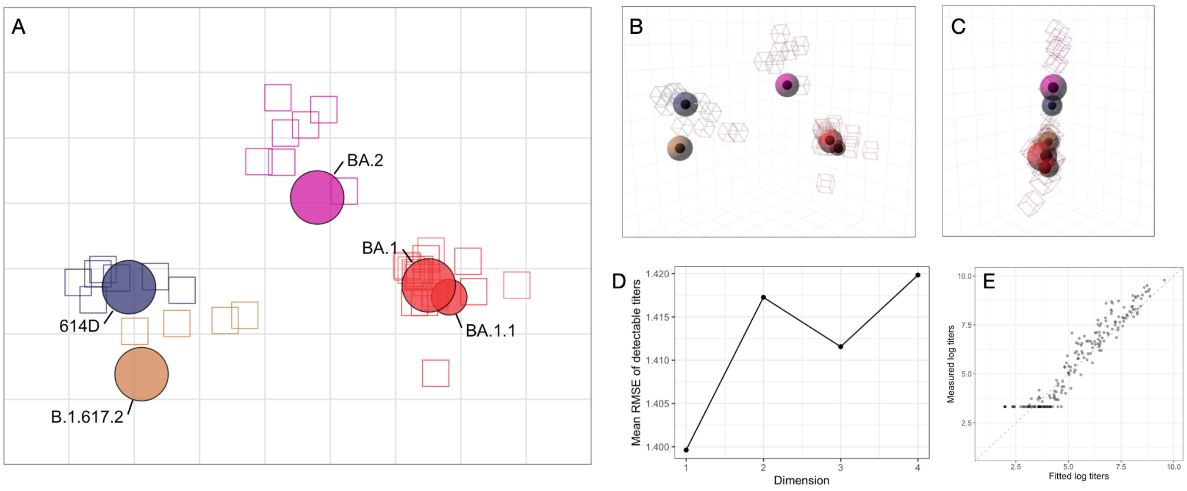


**Figure S23: Madison (FRNT) antigenic map.** A) Detailed labeled figure of the Madison (FRNT) antigenic map. B) Madison (FRNT) antigenic map optimized in three dimensions, ‘side’ view. C) Madison (FRNT) antigenic map optimized in three dimensions, ‘front’ view. D) Dimensionality test for Madison (FRNT) antigenic map. Three-dimensional maps were constructed with 500 optimisations. Dimensionality tests were performed with 500 replicates per dimension and 500 optimisations. E) Scatter plot of fitted titers and measured titers. The fitted titers are determined from the distances in the antigenic map. Titers are plotted on the $log_2$ scale. The dashed gray line is the line of best fit. Non-detectable measured titers were set to 1. In B and C, the black lines point to the position of the variant in the two dimensional map.

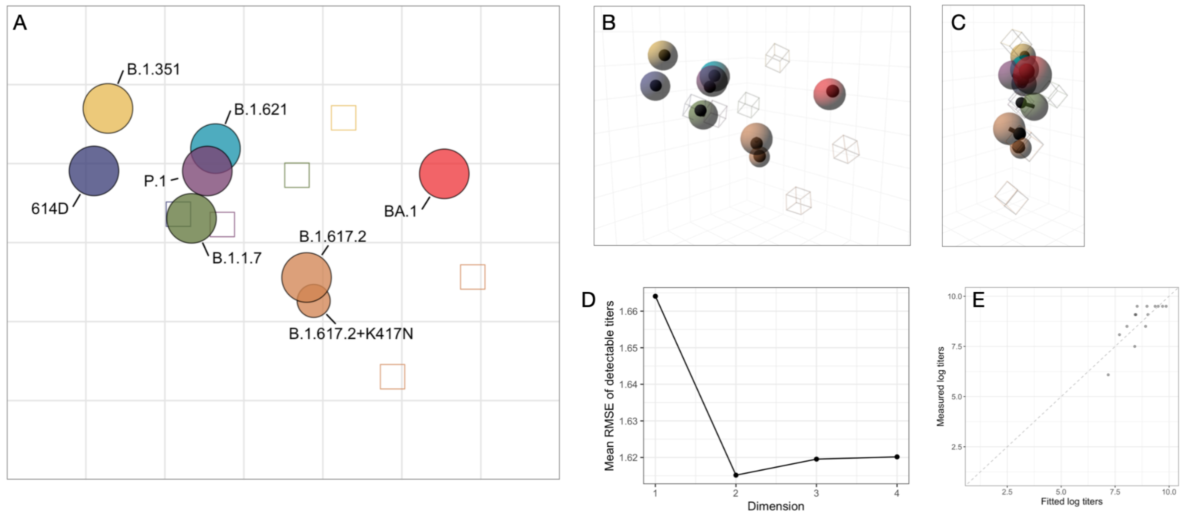


**Figure S24: Maryland antigenic map.** A) Detailed labeled figure of the Maryland antigenic map. B) Maryland antigenic map optimized in three dimensions, ‘side’ view. C) Maryland antigenic map optimized in three dimensions, ‘front’ view. D) Dimensionality test for Maryland antigenic map. Three-dimensional maps were constructed with 500 optimisations. Dimensionality tests were performed with 500 replicates per dimension and 500 optimisations. E) Scatter plot of fitted titers and measured titers. The fitted titers are determined from the distances in the antigenic map. Titers are plotted on the $\log_2$ scale. The dashed gray line is the line of best fit. Non-detectable measured titers were set to 1. In B and C, the black lines point to the position of the variant in the two dimensional map.

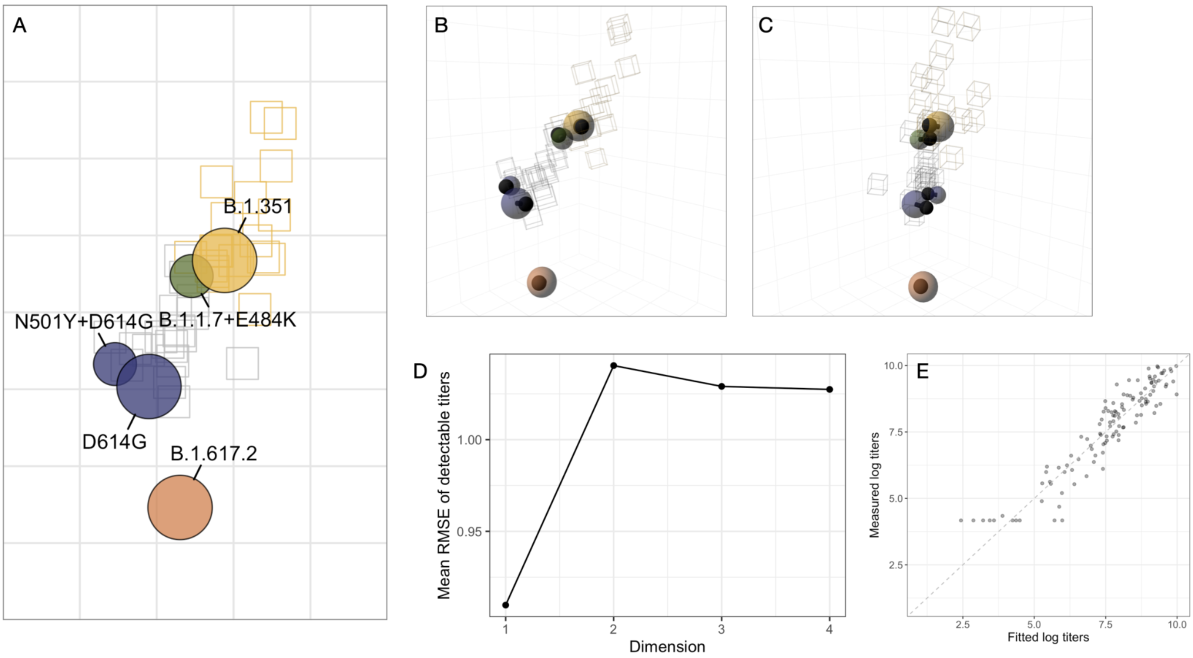


**Figure S25: WUSTL antigenic map.** A) Detailed labeled figure of the WUSTL antigenic map. B) WUSTL antigenic map optimized in three dimensions, 'side' view. C) WUSTL antigenic map optimized in three dimensions, 'front' view. D) Dimensionality test for the WUSTL antigenic map. Three-dimensional maps were constructed with 500 optimisations. Dimensionality tests were performed with 500 replicates per dimension and 500 optimisations. E) Scatter plot of fitted titers and measured titers. The fitted titers are determined from the distances in the antigenic map. Titers are plotted on the $\log_2$ scale. The dashed gray line is the line of best fit. Non-detectable measured titers were set to 1. In B and C, the black lines point to the position of the variant in the two dimensional map.

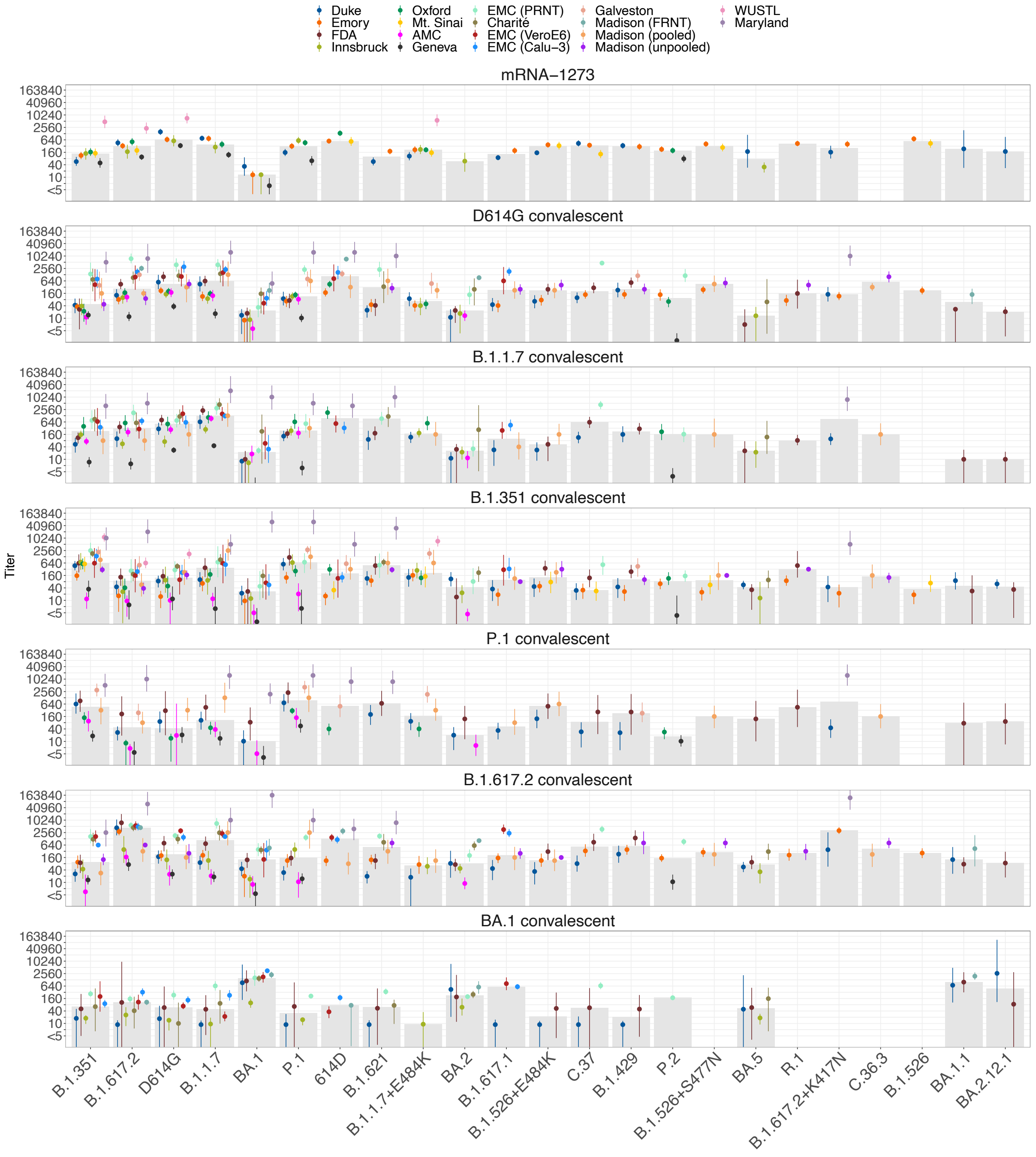


**Figure S26: Absolute titers for all datasets, colored by dataset.** Each dot corresponds to the estimated GMT of a variant titrated against all sera in a particular dataset and serum group. Bars correspond to the 95% highest posterior density interval. GMT and highest posterior density intervals were computed as described in the Materials and Methods. Dots are colored per dataset. The gray bars correspond to the median of the GMTs of the individual datasets. Antigens are ordered by the number of datasets they were titrated in. Within each antigen, datasets are ordered by animal model.

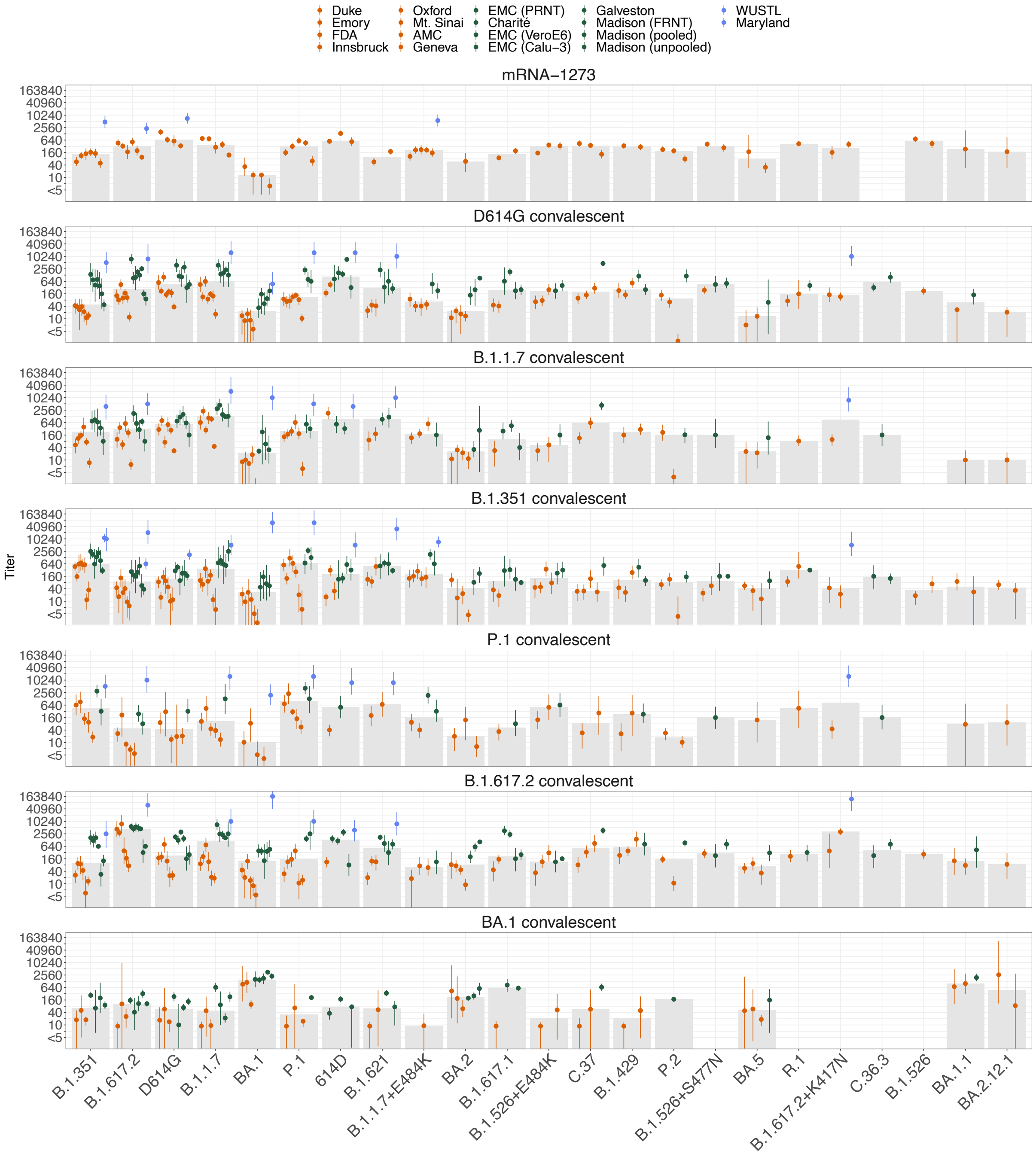


**Figure S27: Absolute titers for all datasets, colored by animal model.** Each dot corresponds to the GMT of a variant titrated against all sera in a serum group in a particular dataset. Bars correspond to the 95% highest posterior density interval. GMT and highest posterior density intervals were computed as described in the Methods. Dots are colored by the animal model (red: human, green: hamster, blue: mouse). The gray bars correspond to the median of the GMTs of the individual datasets. Antigens are ordered by the number of datasets they were titrated in. Within each antigen, datasets are ordered by animal model.

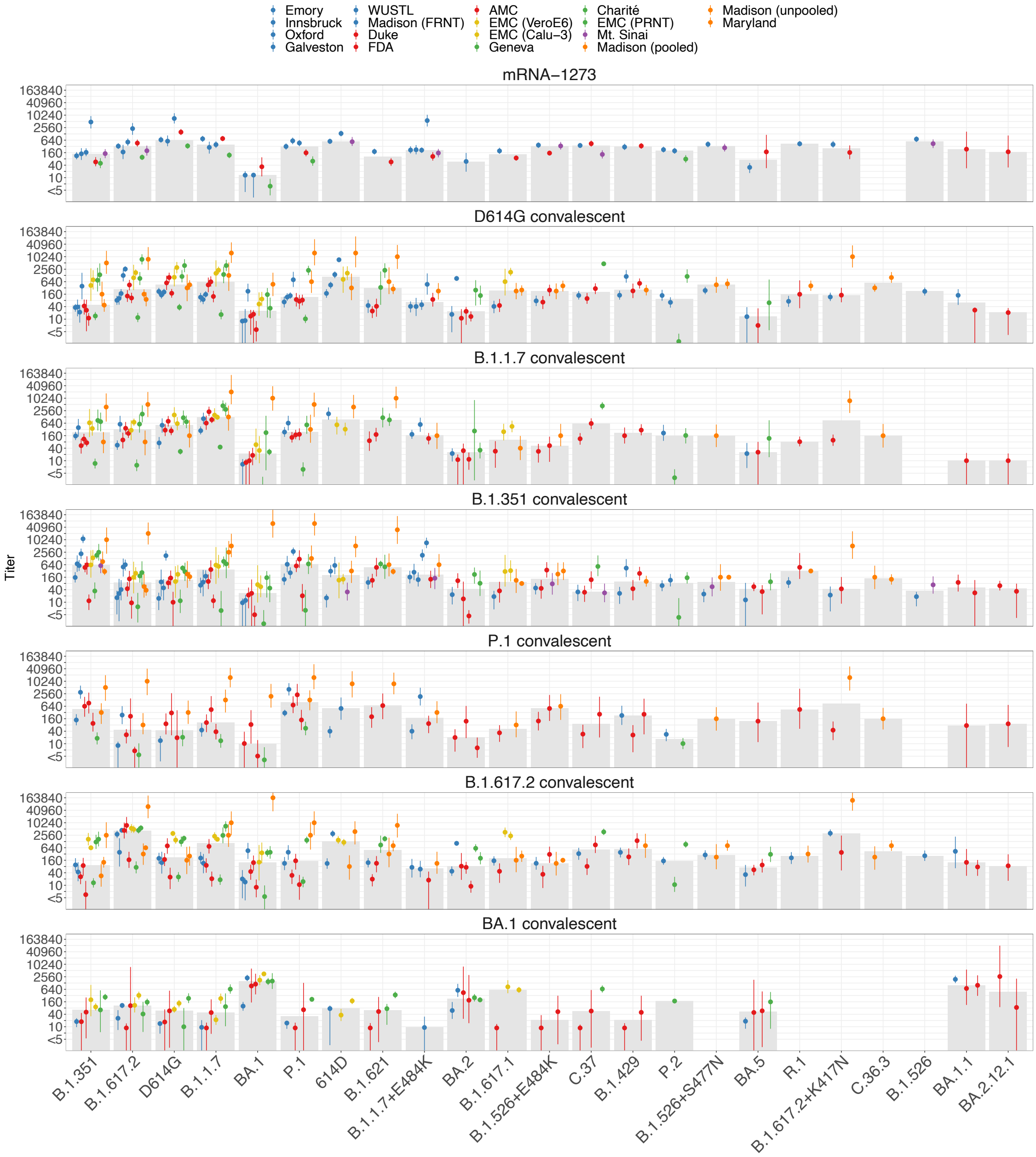


**Figure S28: Absolute titers for all datasets, colored by assay.** Blue: FRNT, red: lentivirus pseudotype neutralization, yellow: VSV pseudotype neutralization, green: PRNT, purple: Microneutralization, orange: CPE. Each dot corresponds to the GMT of a variant titrated against all sera in a particular serum group and dataset. Bars correspond to the 95% highest posterior density interval. GMT and highest posterior density intervals were computed as described in the Methods. The gray bars correspond to the median of the GMTs of the individual datasets. Antigens are ordered by the number of datasets they were titrated in. Within each antigen, datasets are ordered by assay.

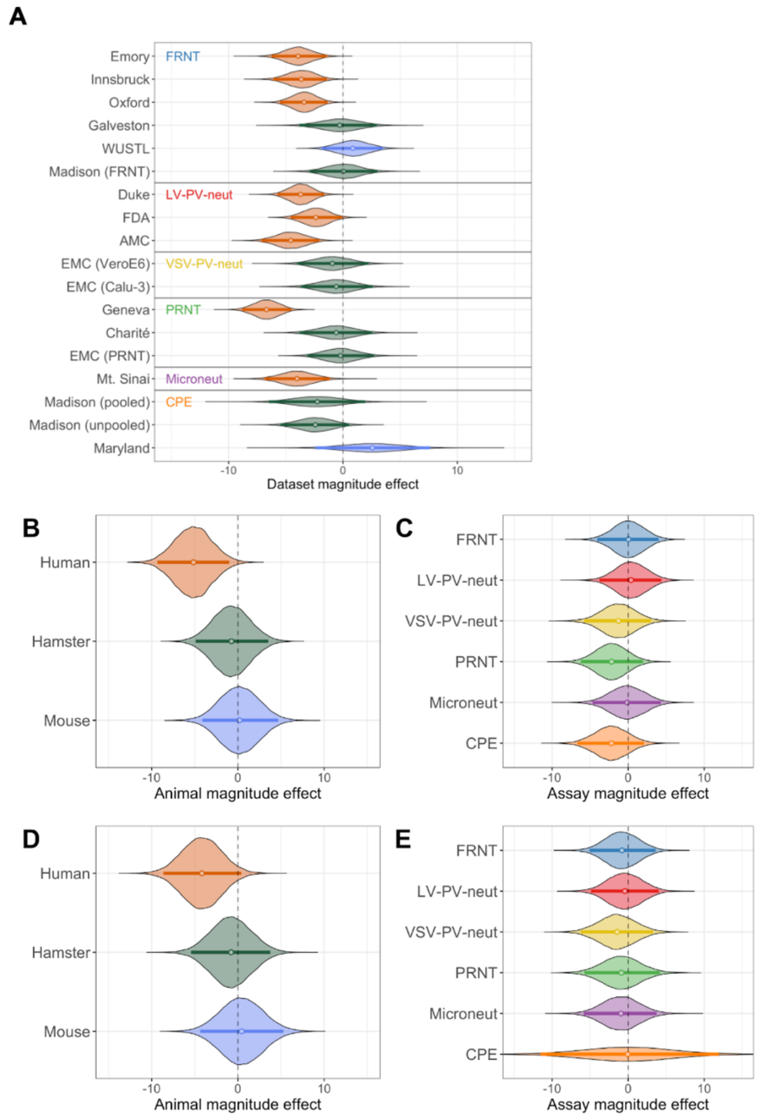


**Figure S29: Dataset magnitude effect.** A) Dataset magnitude effect highlighting differences by assay. B) Effect of the animal model on titer magnitude. C) Effect of the assay on titer magnitude. D and E) Analogous figures to B and C, but with the parameters inferred without datasets that did not use a neutralization cut-off of 50% (Charité, Geneva, AMC, Maryland, Madison (pooled), and Madison (unpooled)). In all figures, the bars show the 95% highest posterior density interval, the dot denotes the mean.

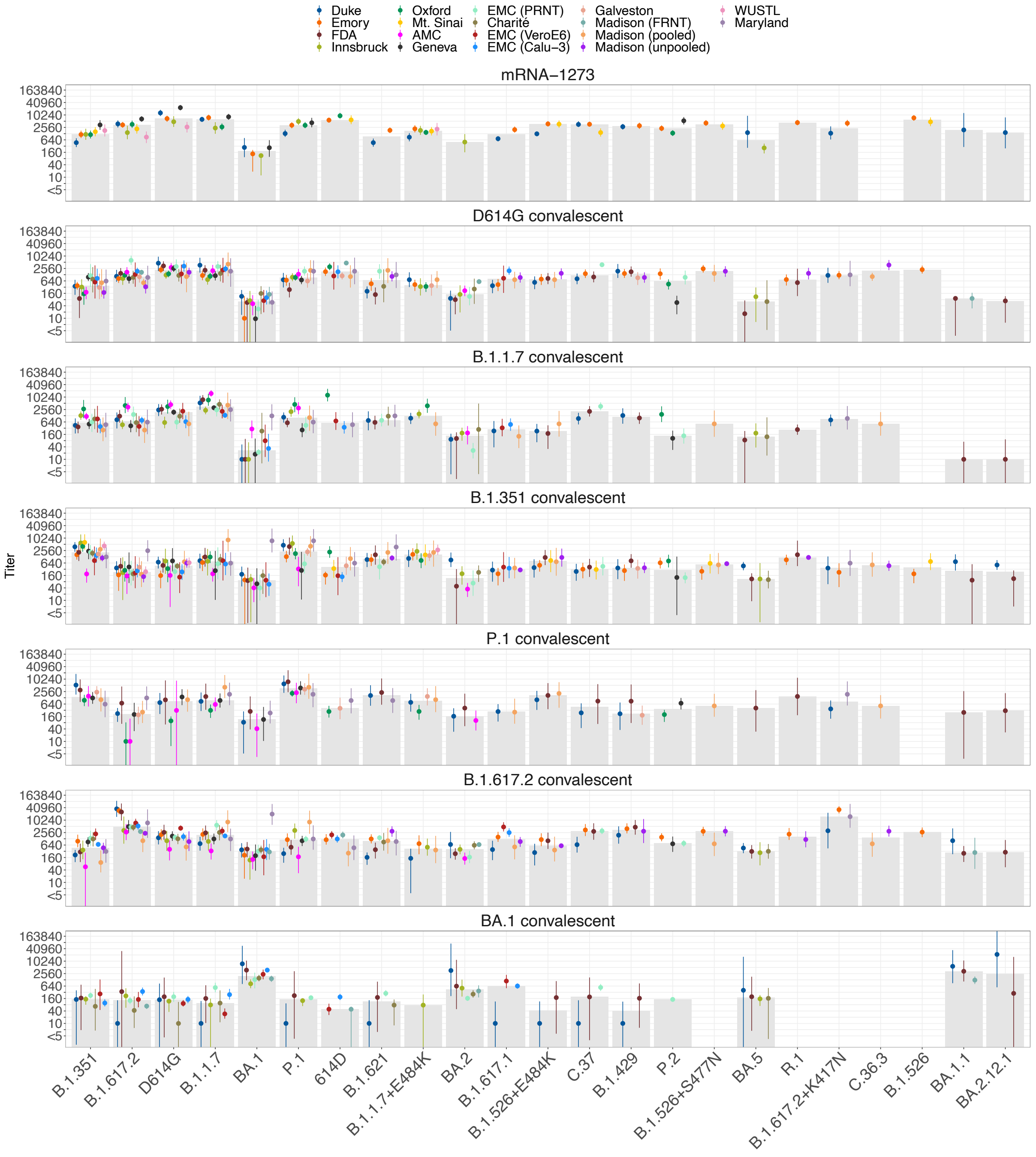


**Figure S30: Titers adjusted for dataset effect, colored by dataset.** Each dot corresponds to the estimated GMT of a variant titrated against all sera in a particular serum group and dataset. Bars correspond to the 95% highest posterior density interval. GMT and highest posterior density intervals were computed as described in the Methods. Dots are colored by the dataset. The gray bars correspond to the median of the GMTs of the individual datasets. Antigens are ordered by the number of datasets they were titrated in. Within each antigen, datasets are ordered by animal model.

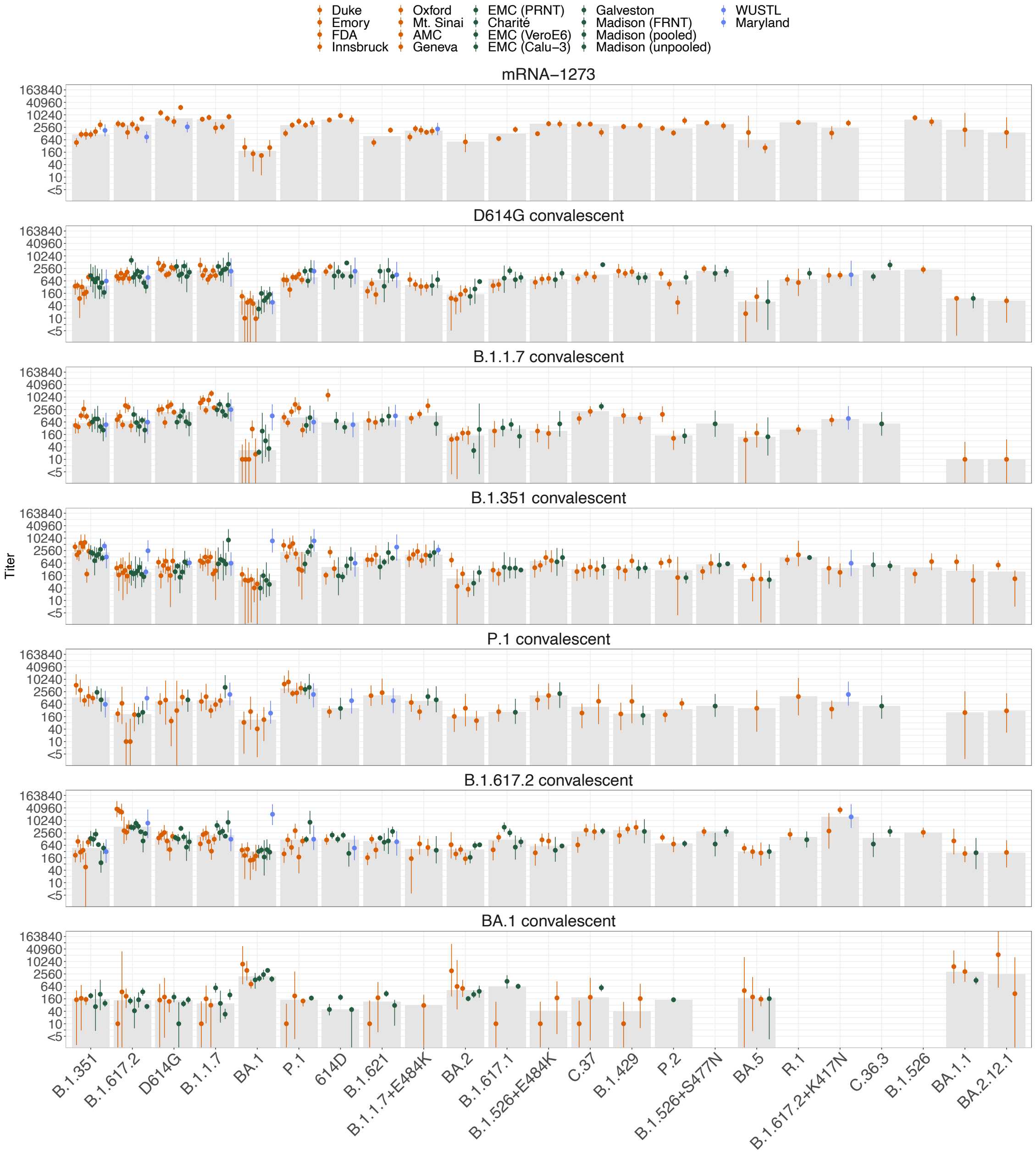


**Figure S31: Titers adjusted for dataset effect, colored by animal model.** Each dot corresponds to the estimated GMT of a variant titrated against all sera in a particular serum group and dataset. Bars correspond to the 95% highest posterior density interval. GMT and highest posterior density intervals were computed as described in the Methods. Dots are colored by the animal model. (red: human, green: hamster, blue: mouse). The gray bars correspond to the median of the GMTs of the individual datasets. Antigens are ordered by the number of datasets they were titrated in. Within each antigen, datasets are ordered by animal model.

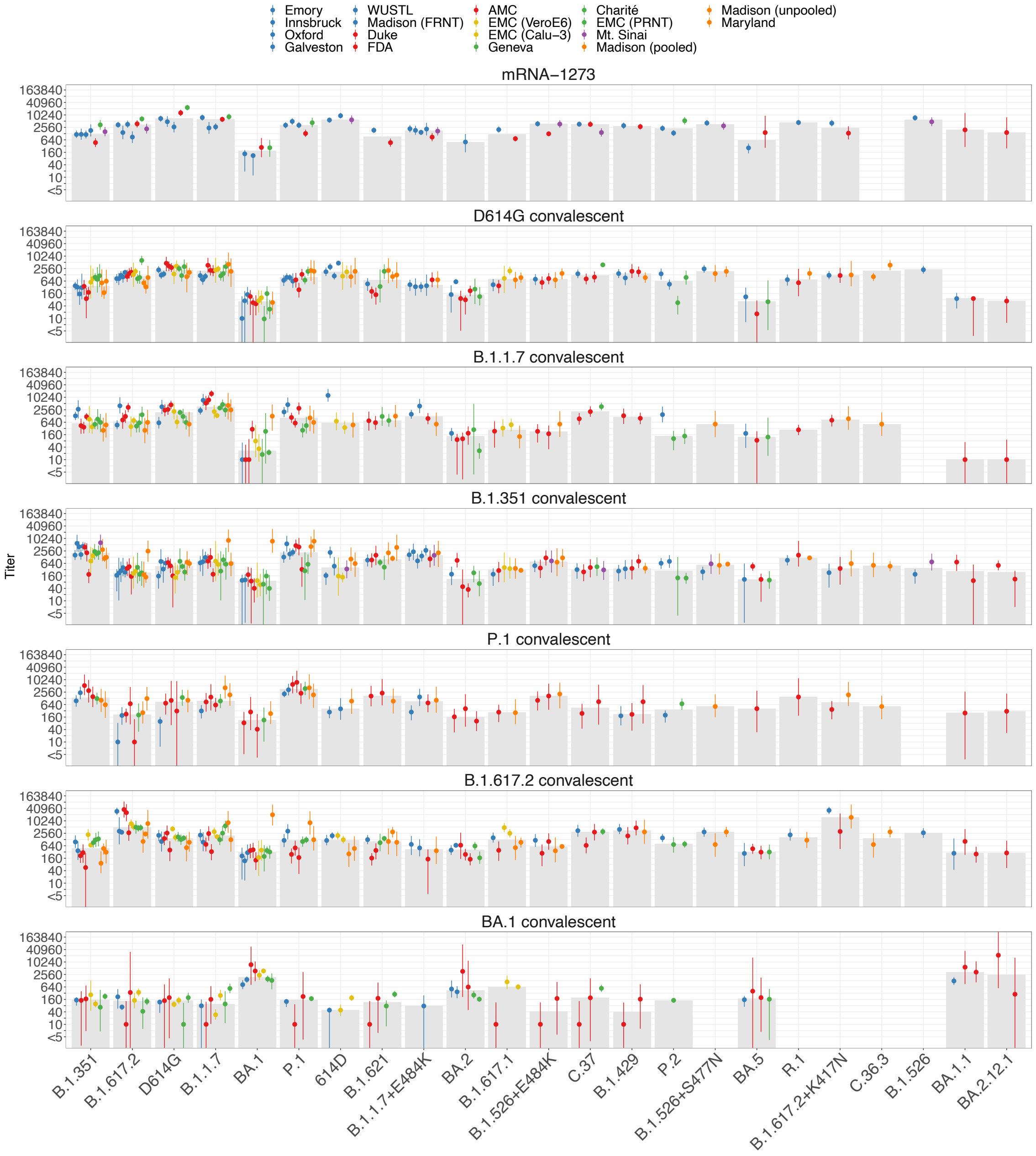


**Figure S32: Titers adjusted for dataset effect, colored by assay.** Each dot corresponds to the estimated GMT of a variant titrated against all sera in a particular serum group and dataset. Bars correspond to the 95% highest posterior density interval. GMT and highest posterior density intervals were computed as described in the Methods. Dots are colored by the assay (blue: FRNT, red: pseudovirus neutralization, green: PRNT, purple: Microneutralization, orange: CPE). The gray bars correspond to the median of the GMTs of the individual datasets. Antigens are ordered by the number of datasets they were titrated in. Within each antigen, datasets are ordered by assay.

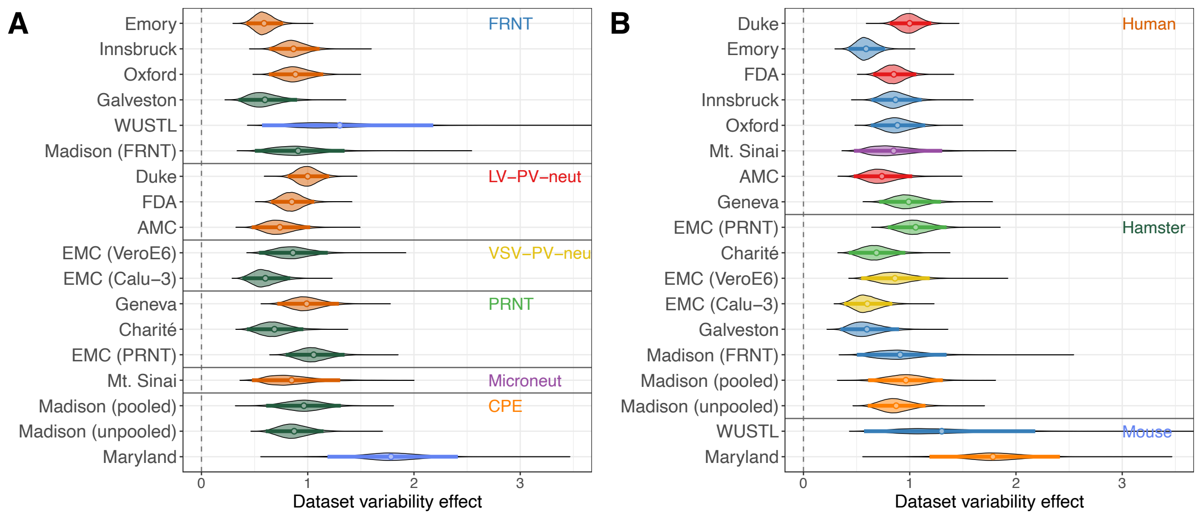


**Figure S33: Dataset variability effect.** A) Ordered by animal model, coloured by assay (blue: FRNT, red: LV-PV-neut, purple: Microneut, green: PRNT, orange: CPE). B) Ordered by assay, coloured by animal model (red: human, green: hamster, blue: mouse). In both figures, the bars show the 95% highest posterior density interval, the dot denotes the mean.

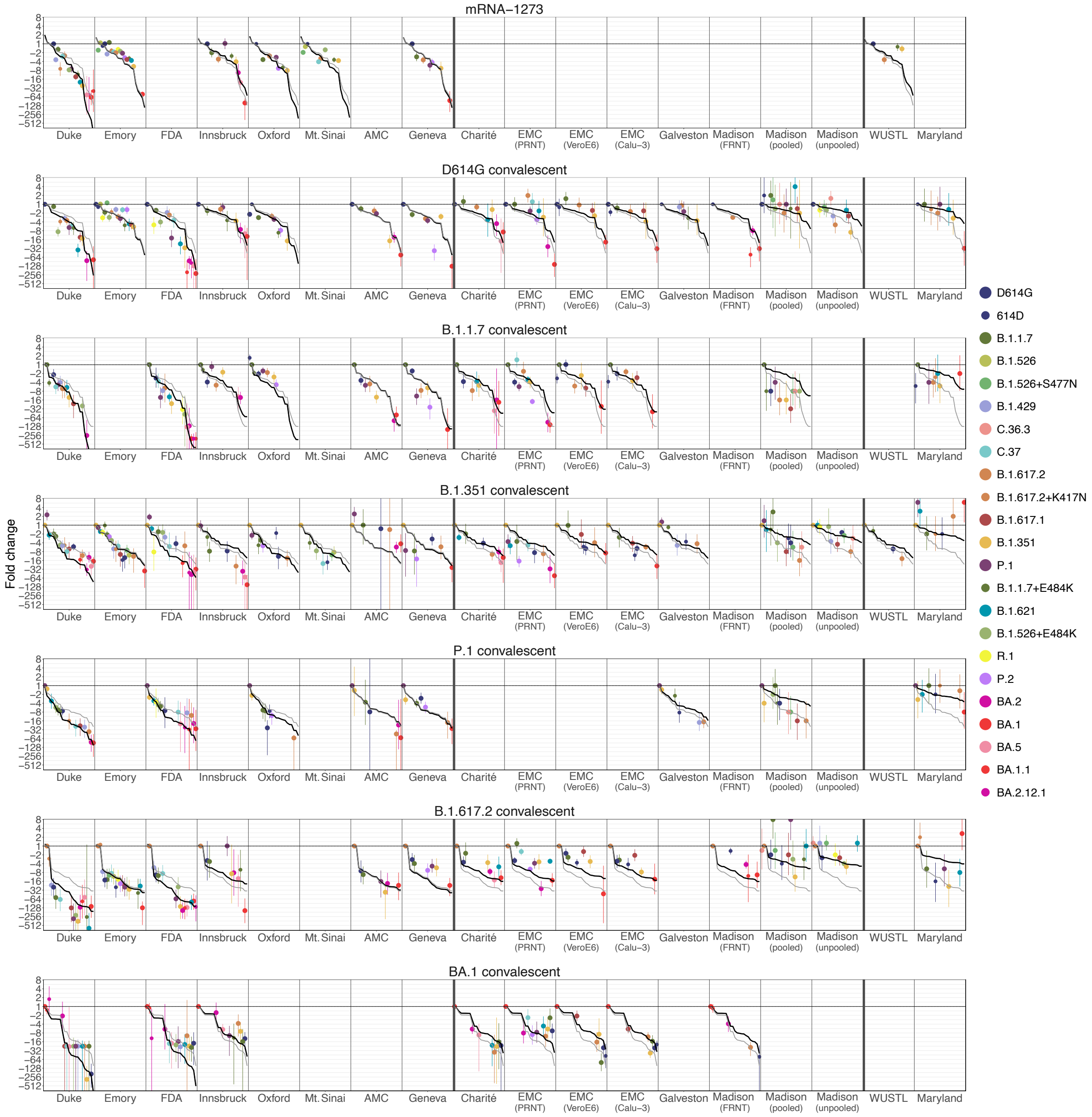


**Figure S34: Fold drop from homologous variant ordered by animal model.** Datasets are grouped on the x-axis with bold vertical lines separating the human, hamster, and mouse datasets. Dots show the mean estimated fold change for each variant and the bars show the estimated 95% highest posterior density intervals, with the colors corresponding to the variant. The light gray line in each panel indicates the estimated fold change per variant in a particular serum group in descending order, calculated across all datasets. The black line shows the estimated fold change in descending order after accounting for differences in fold change between datasets (Material and Methods). Variants are ordered on the x-axis by decreasing estimated fold change, calculated in a particular serum group across all datasets.

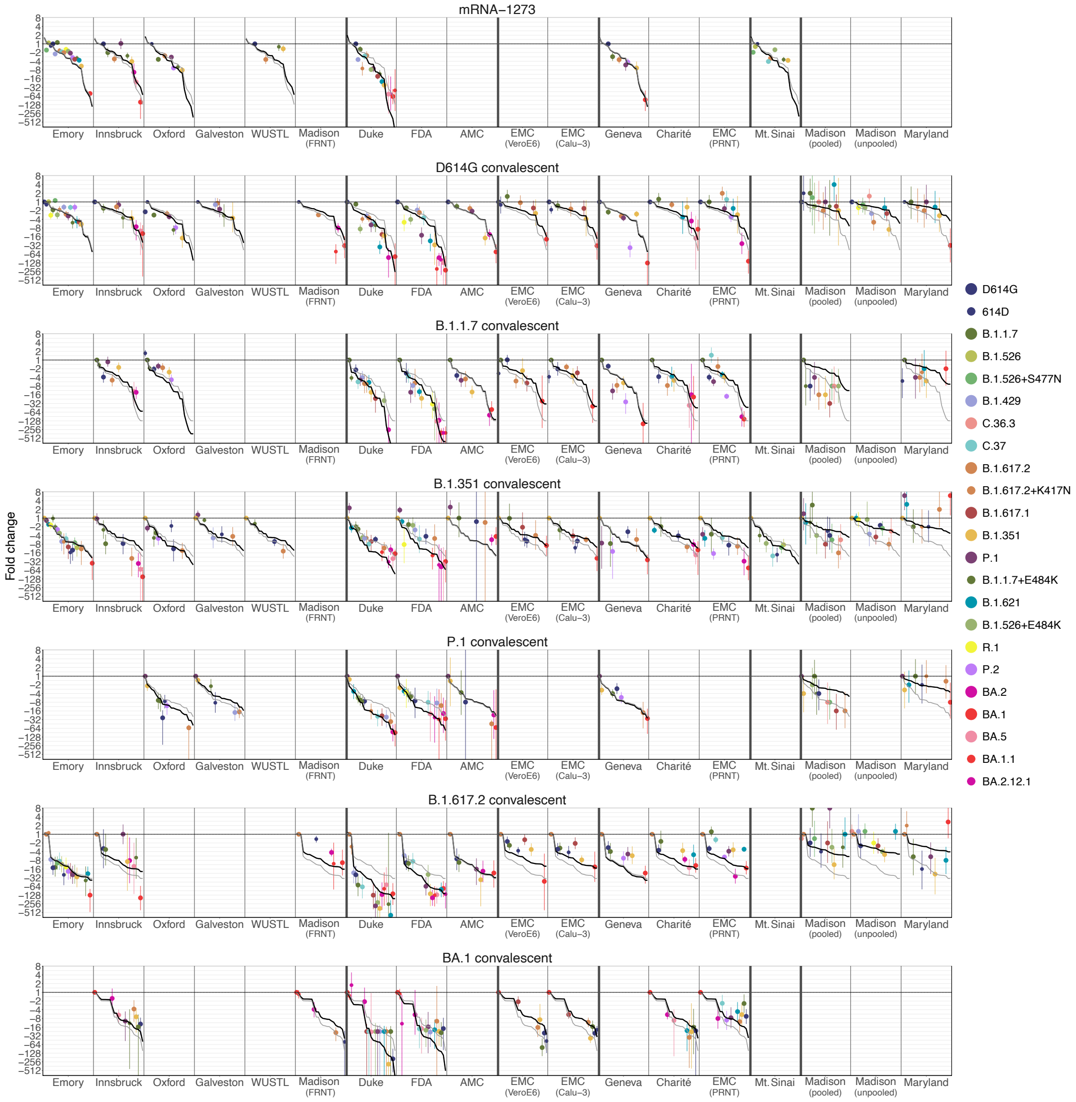


**Figure S35: Fold drop from homologous variant ordered by assay.** Datasets are grouped on the x-axis with bold vertical lines separating the FRNT, lentivirus pseudotype neutralization, VSV pseudotype neutralization, PRNT, Microneutralization, and CPE datasets. Dots show the mean estimated fold change for each variant and the bars show the estimated 95% highest posterior density intervals, with the colors corresponding to the variant. The light gray line in each panel indicates the estimated fold change per variant in a particular serum group in descending order, calculated across all datasets. The black line shows the estimated fold change in descending order after accounting for differences in fold change between datasets (Material and Methods). Variants are ordered on the x-axis by decreasing estimated fold change, calculated in a particular serum group across all datasets.

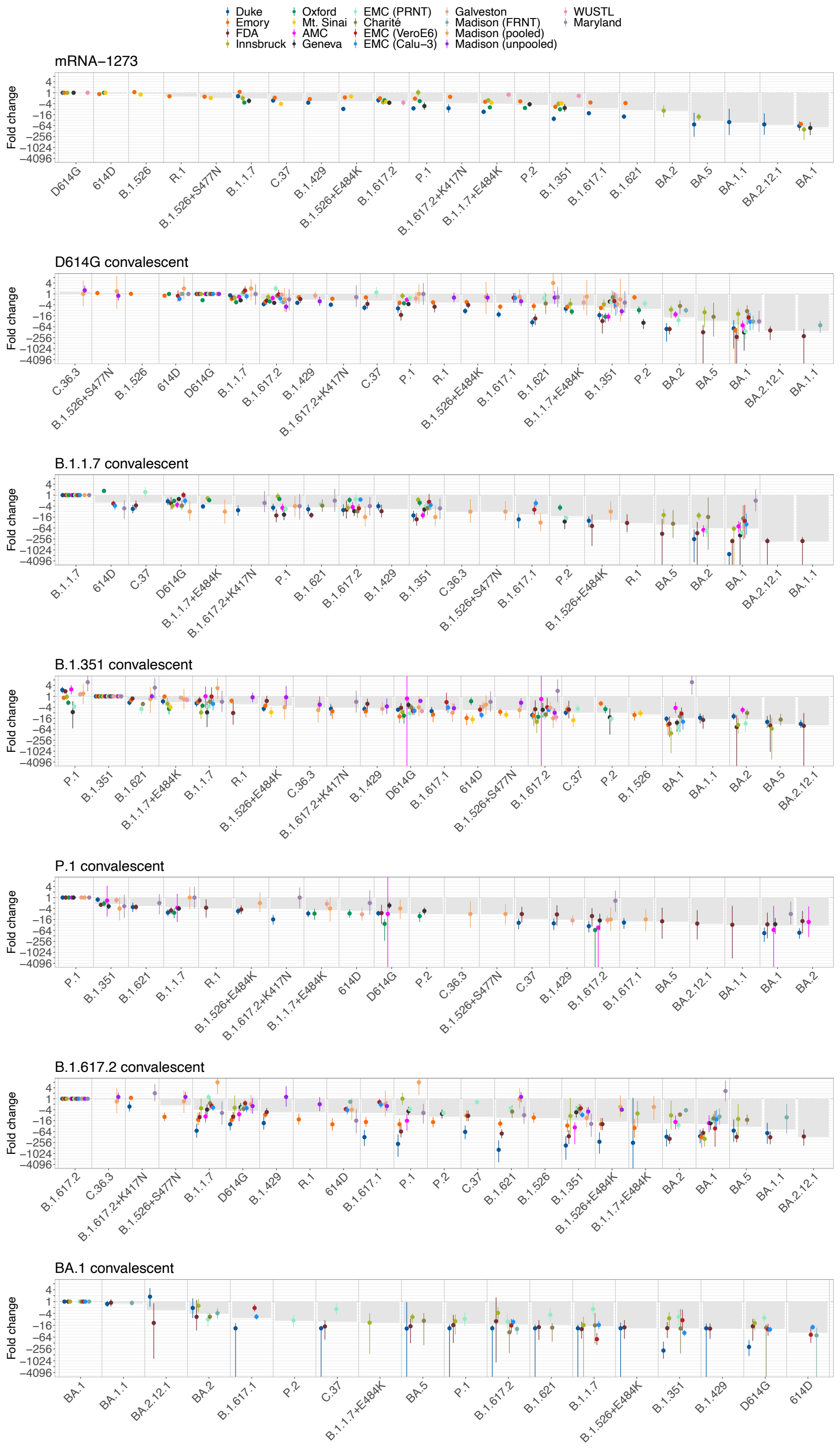


**Figure S36: Fold drop from homologous variant colored by dataset.** Within each variant and serum group combination, datasets are ordered by animal model. Gray bars show the mean fold change calculated from fold changes of all maps for a particular variant and serum group combination. Each dot and vertical line indicate the estimated mean fold drop for a particular dataset and serum group. Dots are colored by dataset.

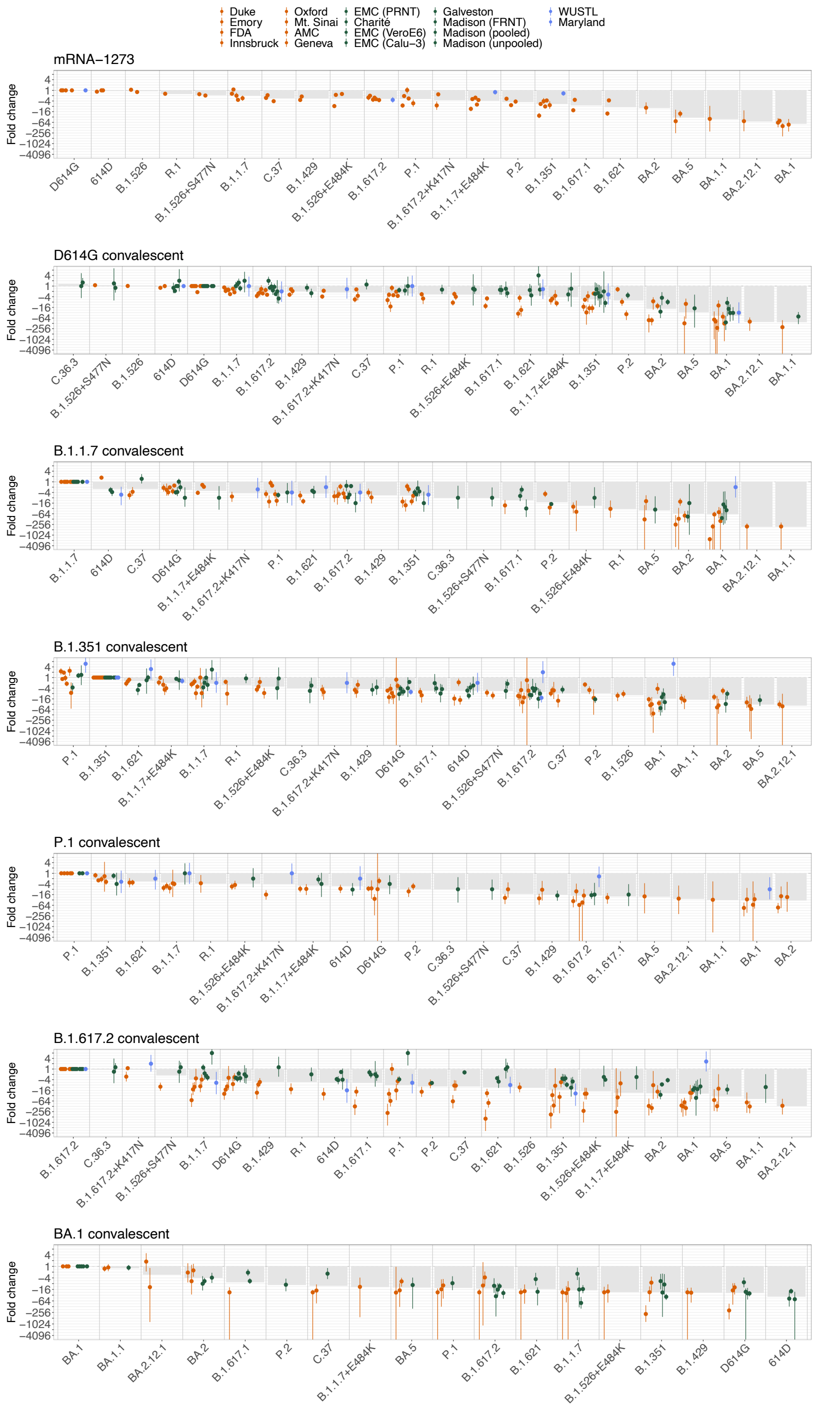


**Figure S37: Fold drop from homologous variant colored by animal model.** Within each variant and serum group combination, datasets are ordered by animal model. Gray bars show the mean fold change calculated from fold changes of all maps for a particular variant and serum group combination. Each dot and vertical line indicate the estimated mean fold drop for a particular dataset and serum group. Dots are colored by animal model (red: human, green: hamster, blue: mouse).

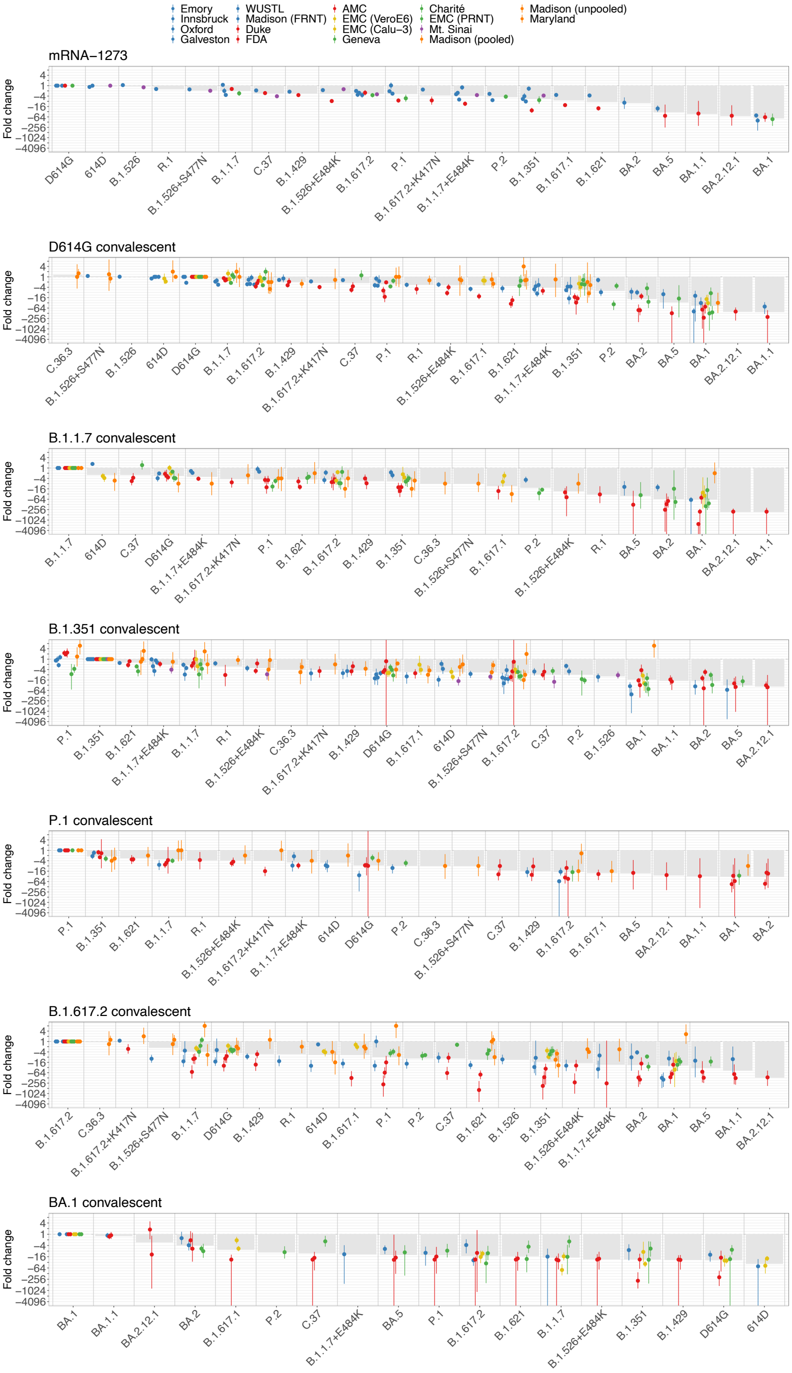


**Figure S38: Fold drop from homologous variant colored by assay.** Within each variant and serum group combination, datasets are ordered by assay. Gray bars show the mean fold change calculated from fold changes of all maps for a particular variant and serum group combination. Each dot and vertical line indicate the estimated mean fold drop for a particular dataset and serum group. Dots are colored by assay (blue: FRNT, red: lentivirus pseudotype neutralization, yellow: VSV pseudotype neutralization, green: PRNT, purple: Microneutralization, orange: CPE).

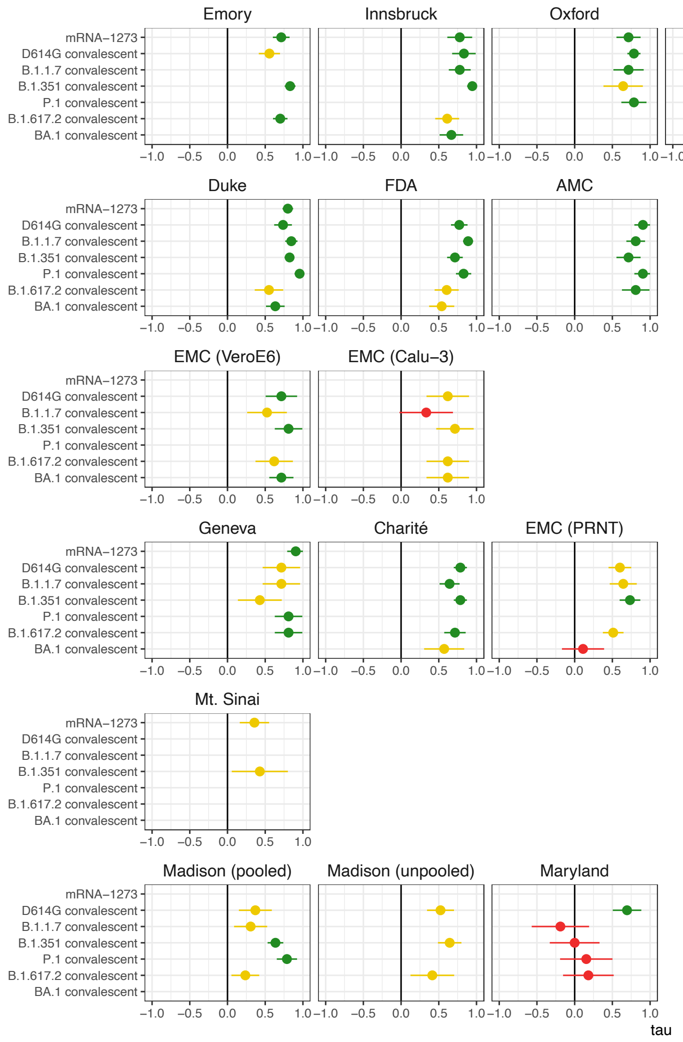


**Figure S39: Kendall rank correlation of mean fold change per serum group across all datasets and fold change per serum group for each individual dataset.** Mean fold change per serum group across all datasets is shown as a gray line in fig. S34. The estimated correlation coefficient is shown in red if the lower bound of the confidence interval is less than 0, yellow if the lower bound of the correlation coefficient is less than 0.5, and green if the lower bound of the correlation coefficient is greater than 0.5. Rows correspond to the different assays top to bottom: FRNT, Lentivirus pseudotype neutralization, VSV pseudotype neutralization, PRNT, Microneutralization, CPE.

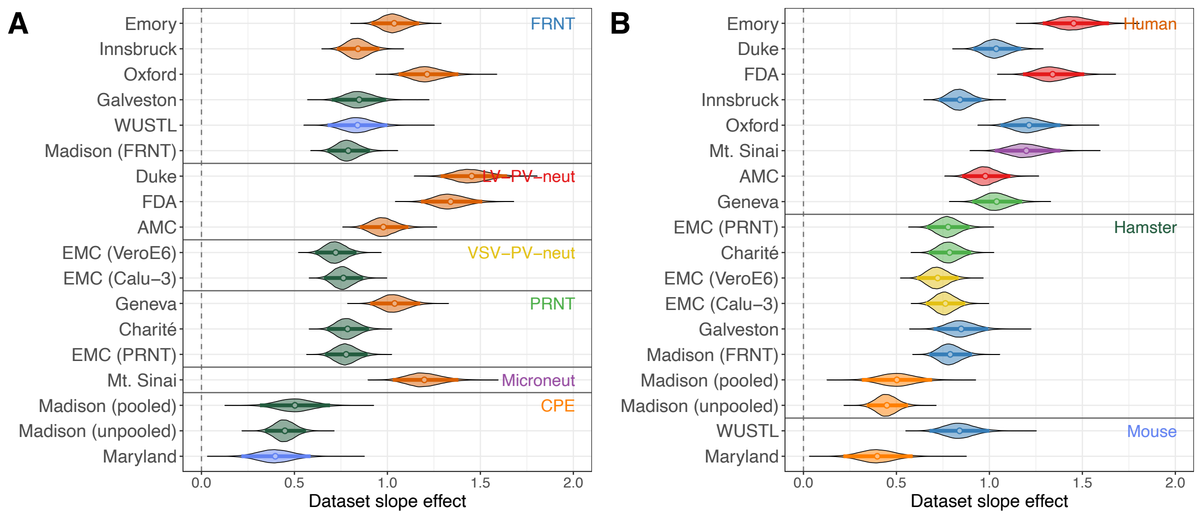


**Figure S40: Amount of fold change measured in each dataset.** Each titer was modeled as a combination of the fold change for a particular variant and serum group combination multiplied with a slope effect given by the dataset (Methods). The bars show the 95% highest posterior density interval, the dot denotes the mean. Panel A and B show the estimated slope effect, ordered by assay and colored by animal model (A), and ordered by animal model and coloured by assay (B) for all datasets.

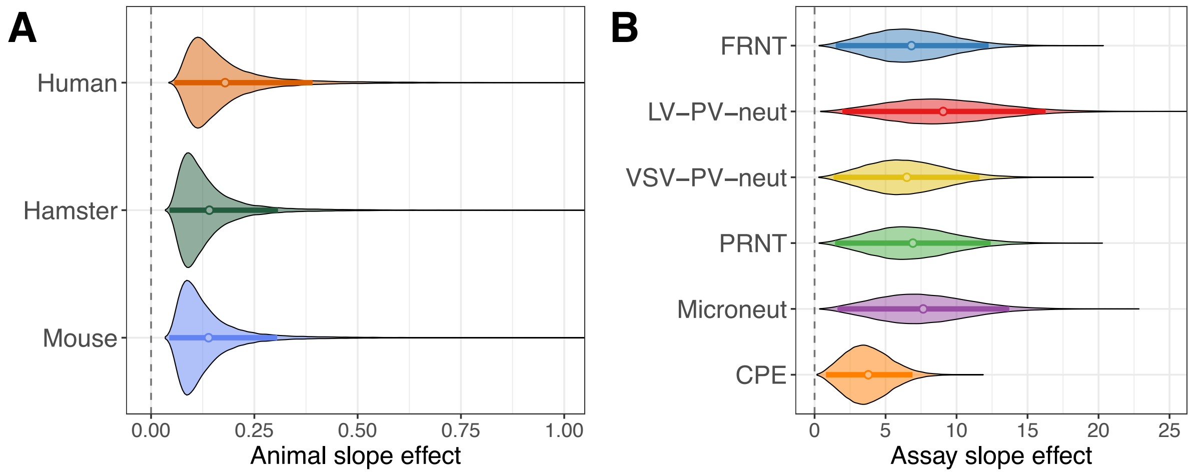


**Figure S41: Effect of different animal models and assays on the amount of fold change measured in each dataset.** A) Effect of the animal model. B) Effect of the assay. The bars show the 95% highest posterior density interval, the dot denotes the mean.

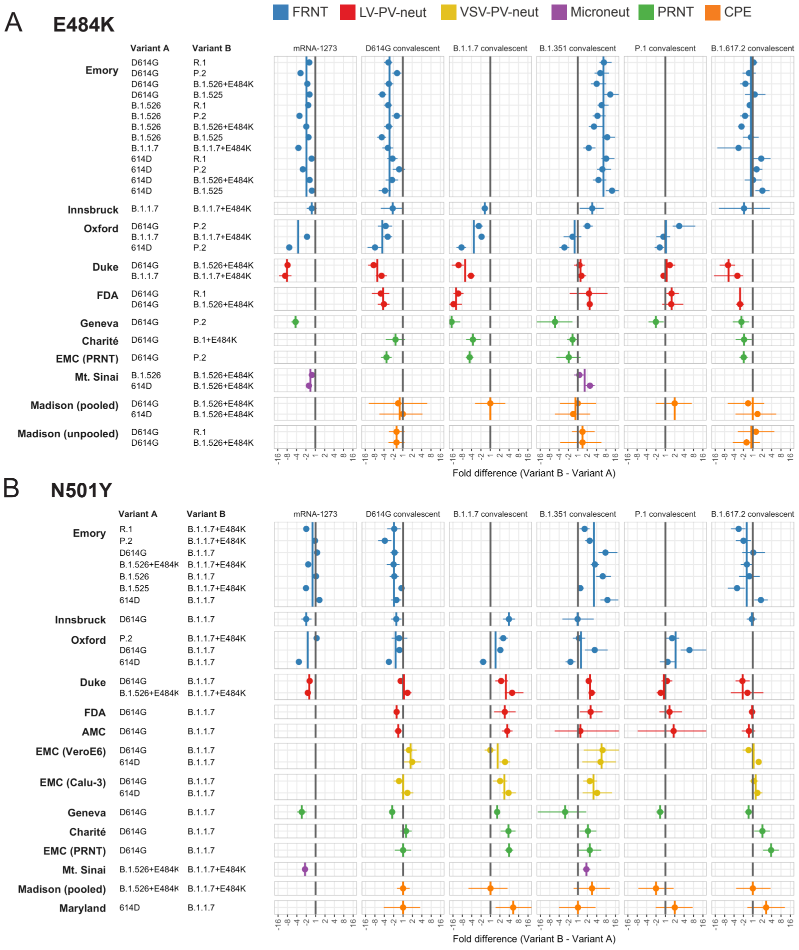


**Figure S42: Sensitivity of different serum groups to the E484K (A) and N501Y (B) substitutions, colored by assay.** Each row shows the average fold difference and 95% highest posterior density interval in titer between the two variants on the left, which differ by only the E484K or the N501Y substitution in the RBD (a positive fold difference corresponds to higher titers against variant with the substitution, while a negative fold difference corresponds to higher titers against the variant without the substitution). Values for each dataset are colored by assay (blue: FRNT, red: lentivirus pseudovirus neutralization, yellow: VSV pseudotype neutralization, green: PRNT, purple: Microneutralization, orange: CPE). Bold vertical lines indicate the average fold difference measured across datasets made with different assays (blue: FRNT, red: lentivirus pseudovirus neutralization, yellow: VSV pseudotype neutralization, green: PRNT, purple: Microneutralization, orange: CPE). The gray line indicates no difference in titers between the two variants.

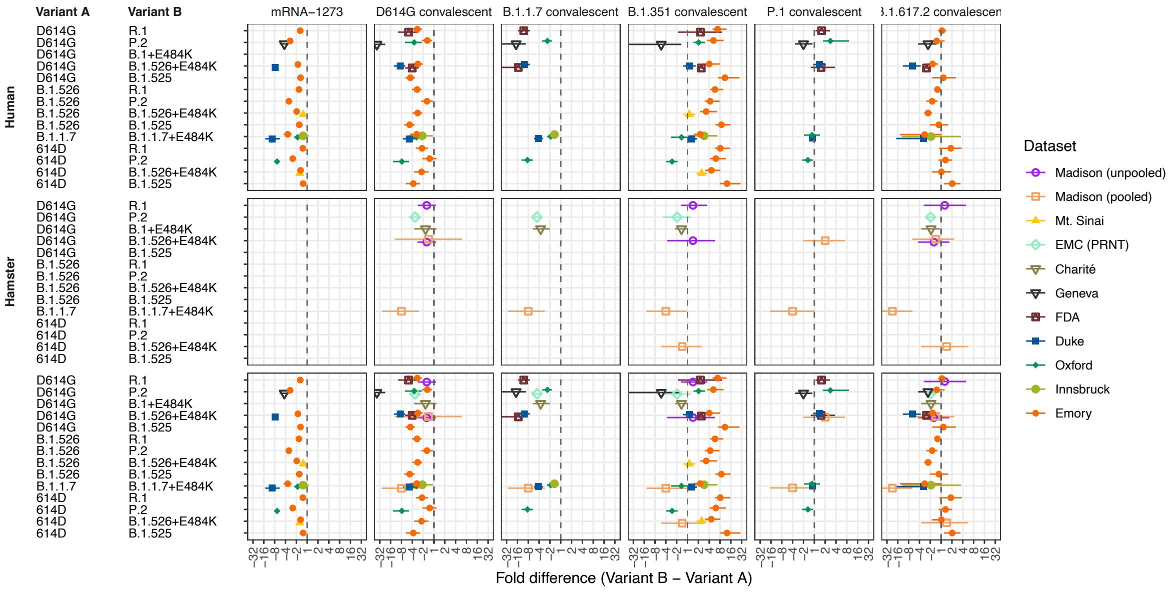


**Figure S43: Sensitivity of different serum groups to the E484K substitution, split by animal model.** Each row shows the average fold difference and 95% highest posterior density interval in titer between the two variants on the left, which differ by only the E484K substitution in the RBD (a positive fold difference corresponds to higher titers against variant with the substitution, while a negative fold difference corresponds to higher titers against the variant without the substitution). The dashed line indicates no difference in titers between the two variants.

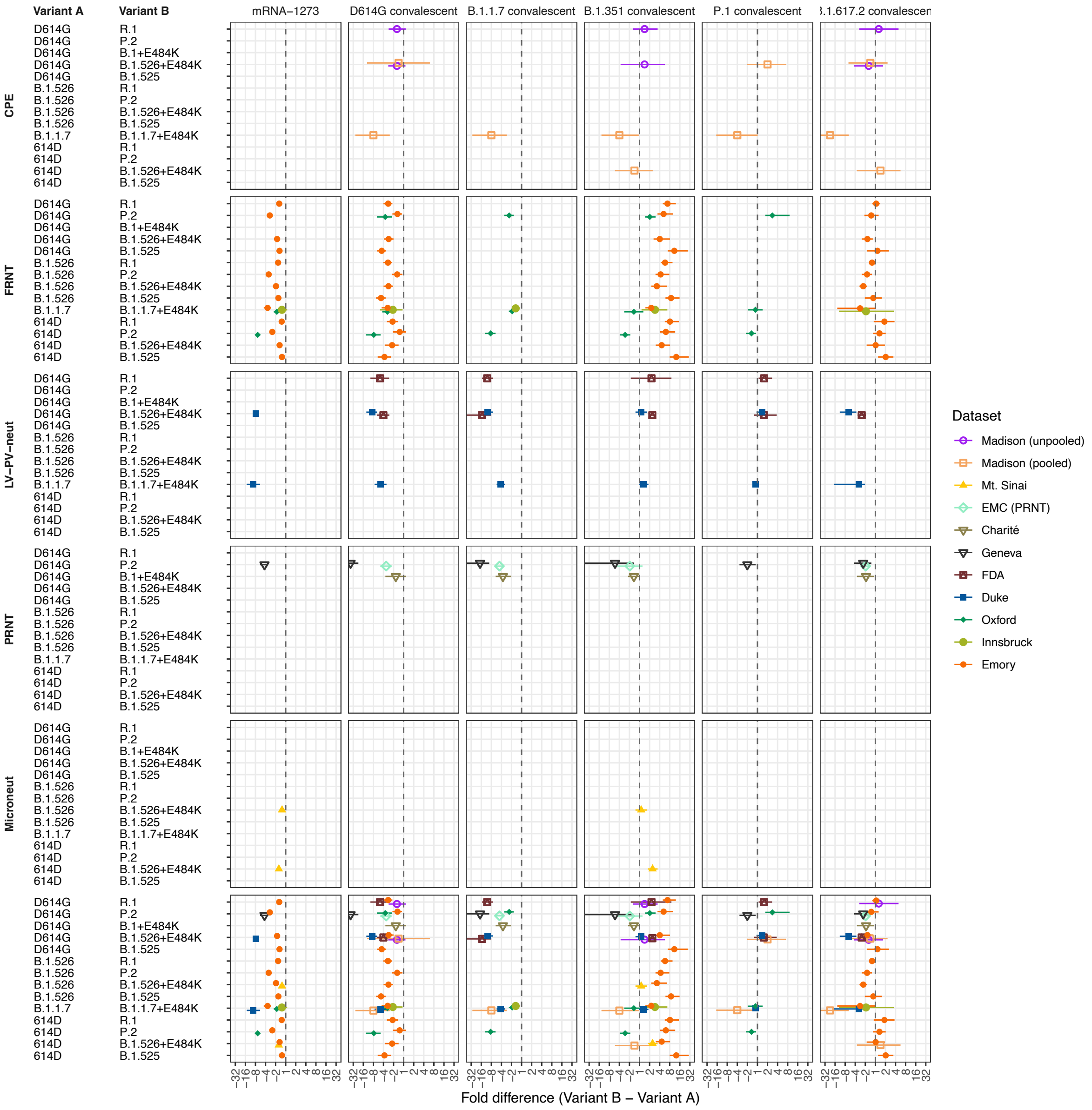


**Figure S44: Sensitivity of different serum groups to the E484K substitution, split by assay.** Each row shows the average fold difference and 95% highest posterior density interval in titer between the two variants on the left, which differ by only the E484K substitution in the RBD (a positive fold difference corresponds to higher titers against variant with the substitution, while a negative fold difference corresponds to higher titers against the variant without the substitution). The dashed line indicates no difference in titers between the two variants.

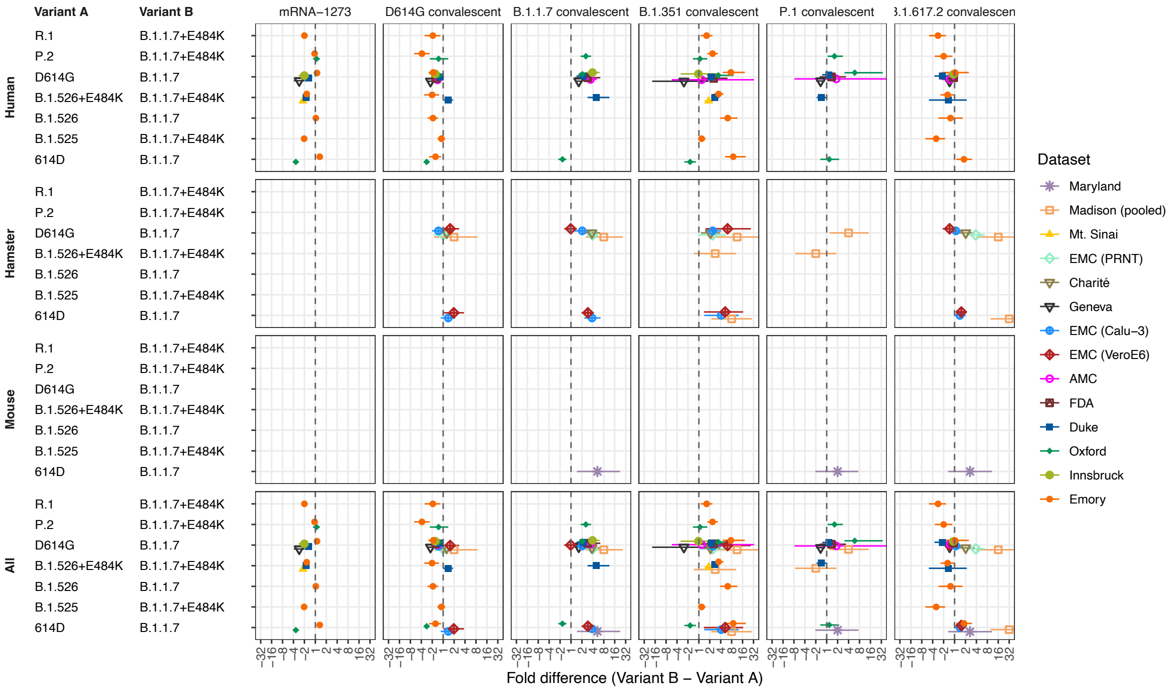


**Figure S45: Sensitivity of different serum groups to the N501Y substitution, split by animal model.** Each row shows the average fold difference and 95% highest posterior density interval in titer between the two variants on the left, which differ by only the N501Y substitution in the RBD (a positive fold difference corresponds to higher titers against variant with the substitution, while a negative fold difference corresponds to higher titers against the variant without the substitution). The dashed line indicates no difference in titers between the two variants.

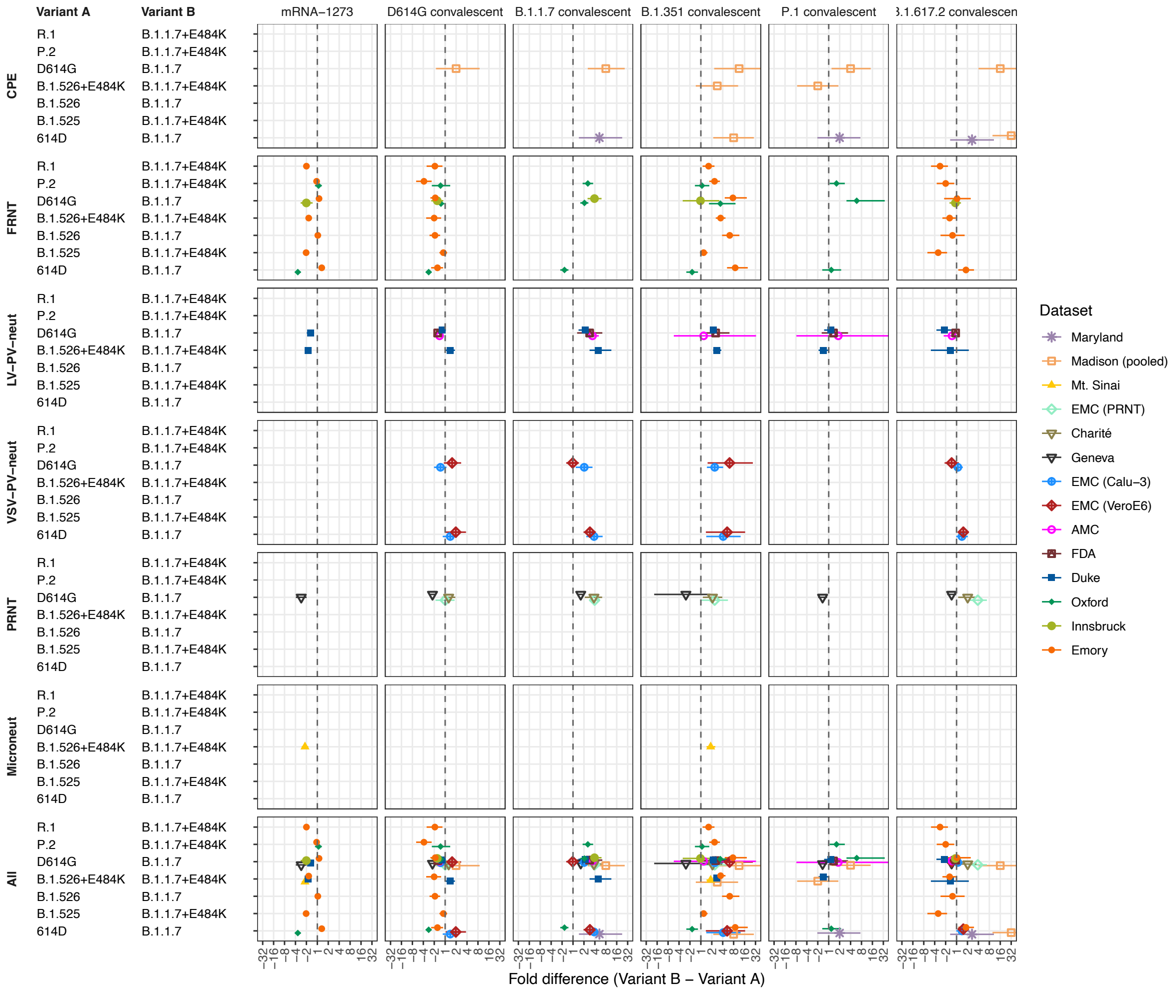


**Figure S46: Sensitivity of different serum groups to the N501Y substitution, split by assay.** Each row shows the average fold difference and 95% highest posterior density interval in titer between the two variants on the left, which differ by only the N501Y substitution in the RBD (a positive fold difference corresponds to higher titers against variant with the substitution, while a negative fold difference corresponds to higher titers against the variant without the substitution). The dashed line indicates no difference in titers between the two variants.

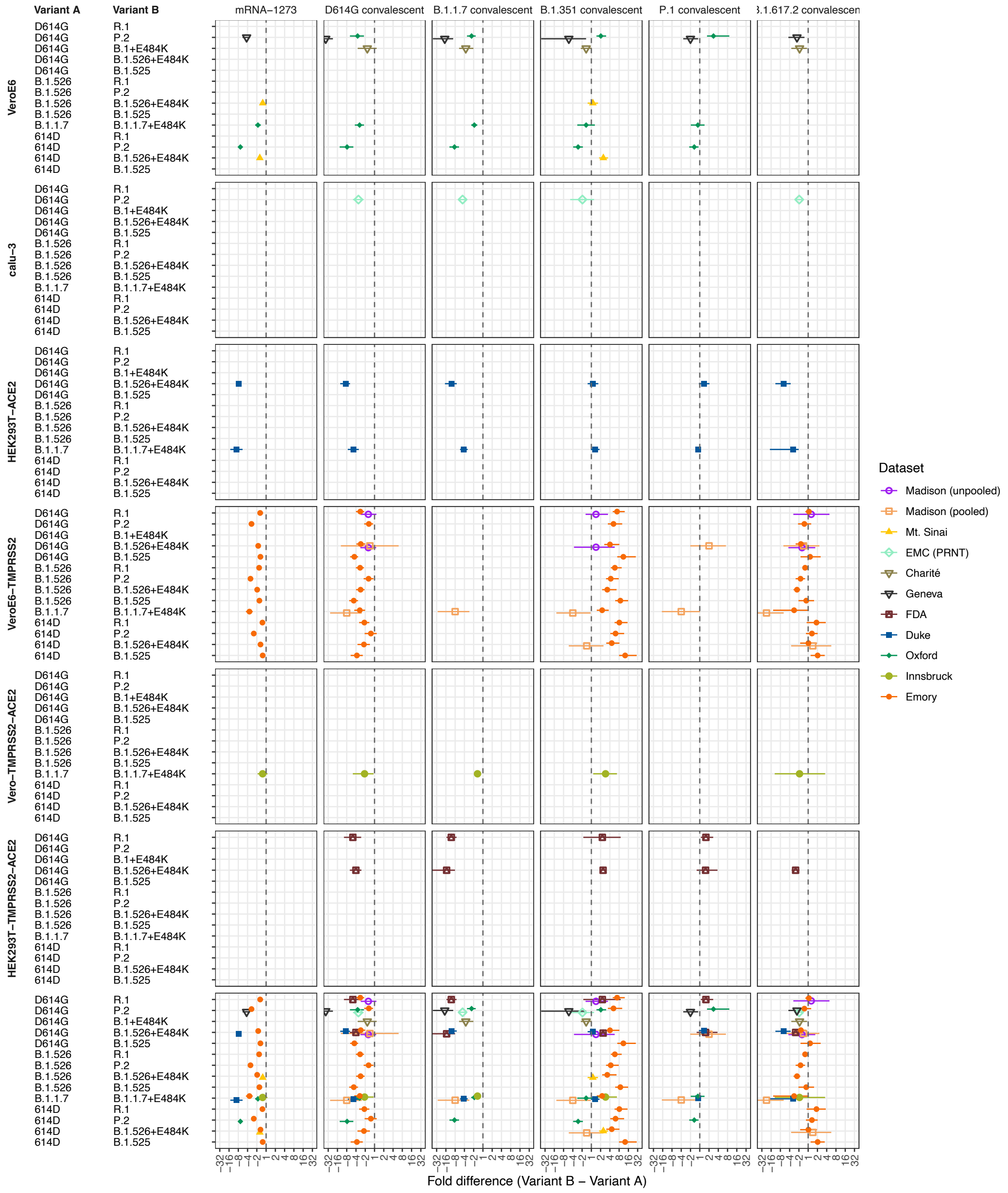


**Figure S47: Sensitivity of different serum groups to the E484K substitution, split by cell type.** Each row shows the average fold difference and 95% highest posterior density interval in titer between the two variants on the left, which differ by only the E484K substitution in the RBD (a positive fold difference corresponds to higher titers against variant with the substitution, while a negative fold difference corresponds to higher titers against the variant without the substitution). The dashed line indicates no difference in titers between the two variants.

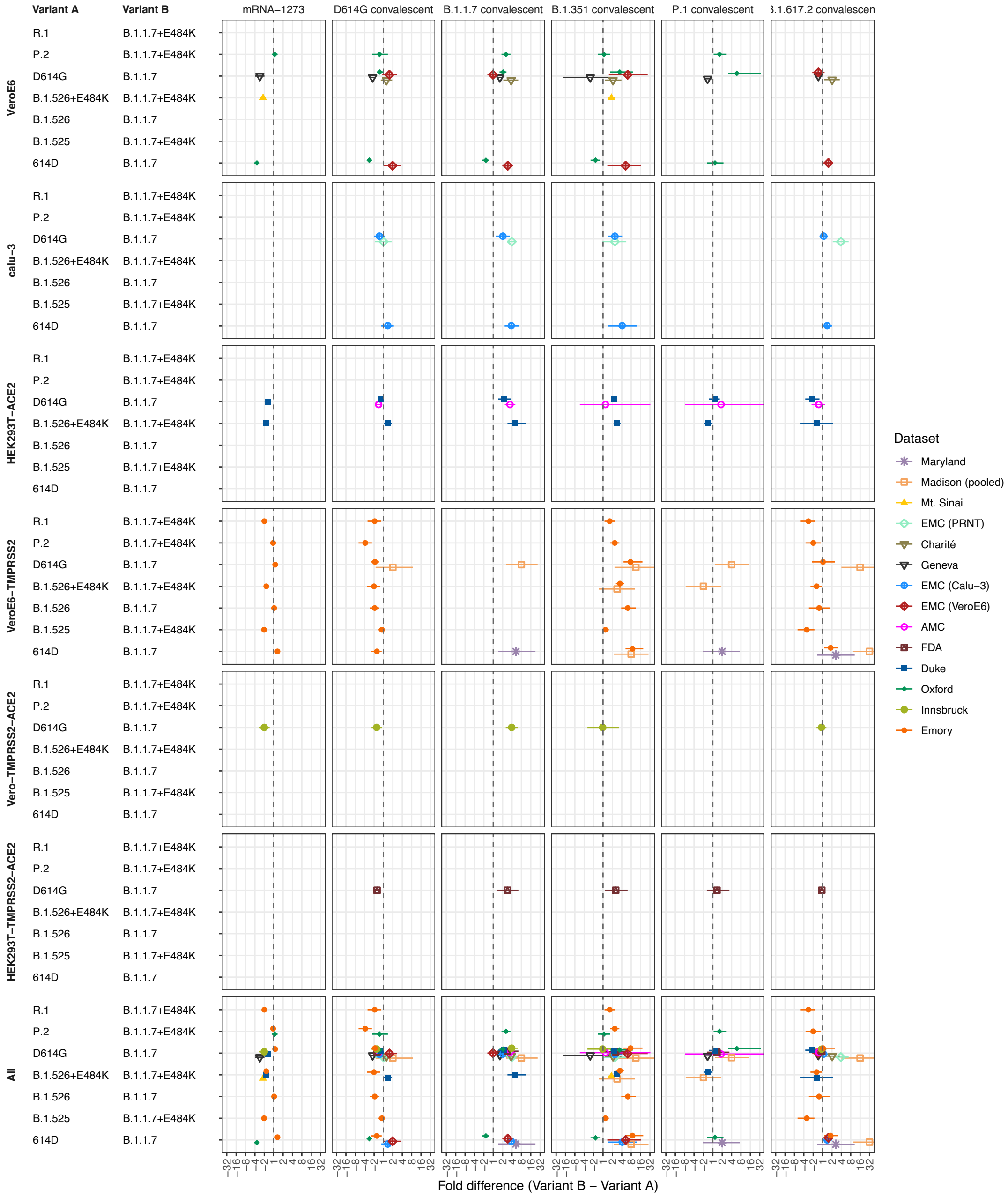


**Figure S48: Sensitivity of different serum groups to the N501Y substitution, split by cell type.** Each row shows the average fold difference and 95% highest posterior density interval in titer between the two variants on the left, which differ by only the N501Y substitution in the RBD (a positive fold difference corresponds to higher titers against variant with the substitution, while a negative fold difference corresponds to higher titers against the variant without the substitution). The dashed line indicates no difference in titers between the two variants.

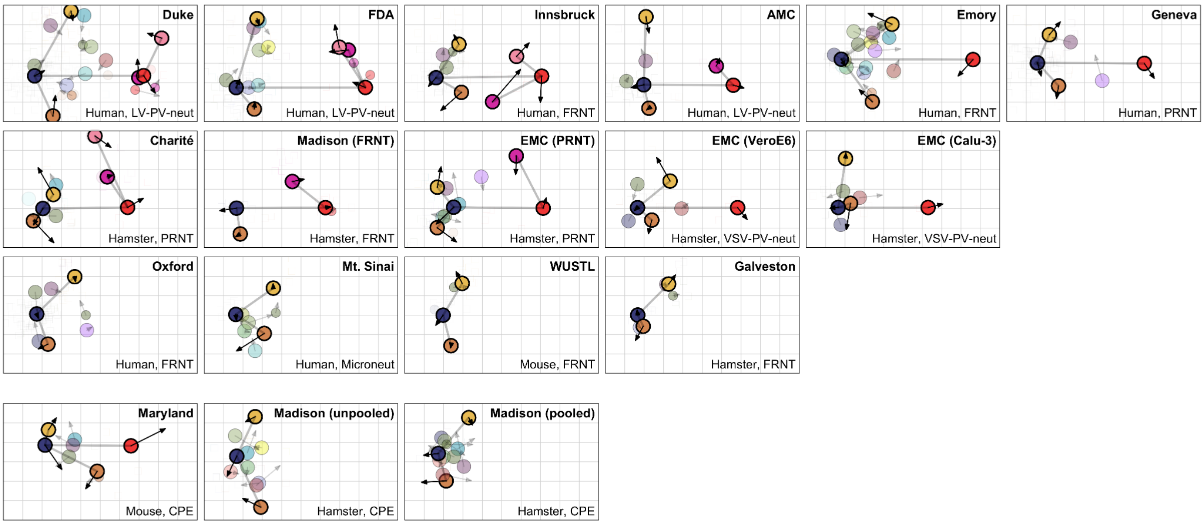


**Figure S49: Antigenic maps showing the relative positions of ancestral, B.1.351, B.1.617.2, BA.1, BA.2, and BA.4/5 variants.** Maps are colored by variant (see fig. S53). Ancestral, B.1.351, B.1.617.2, BA.1, BA.2, and BA.4/5 variants are highlighted with black outlines and faint gray lines are drawn to highlight their relative positions. Arrows show the position of the variants in the merged antigenic map constructed from all titers, shown in Fig. 4B. Maps are oriented such that the line from the ancestral variant (either D614G or 614D) to Omicron BA.1 is horizontal, with Omicron BA.1 on the right. Maps without Omicron BA.1 are oriented such that the ancestral variant (D614G or 614D) is on the left, the B.1.351 variant at the top and the B.1.617.2 variant at the bottom. Data for the maps in the bottom row were generated using CPE assays.

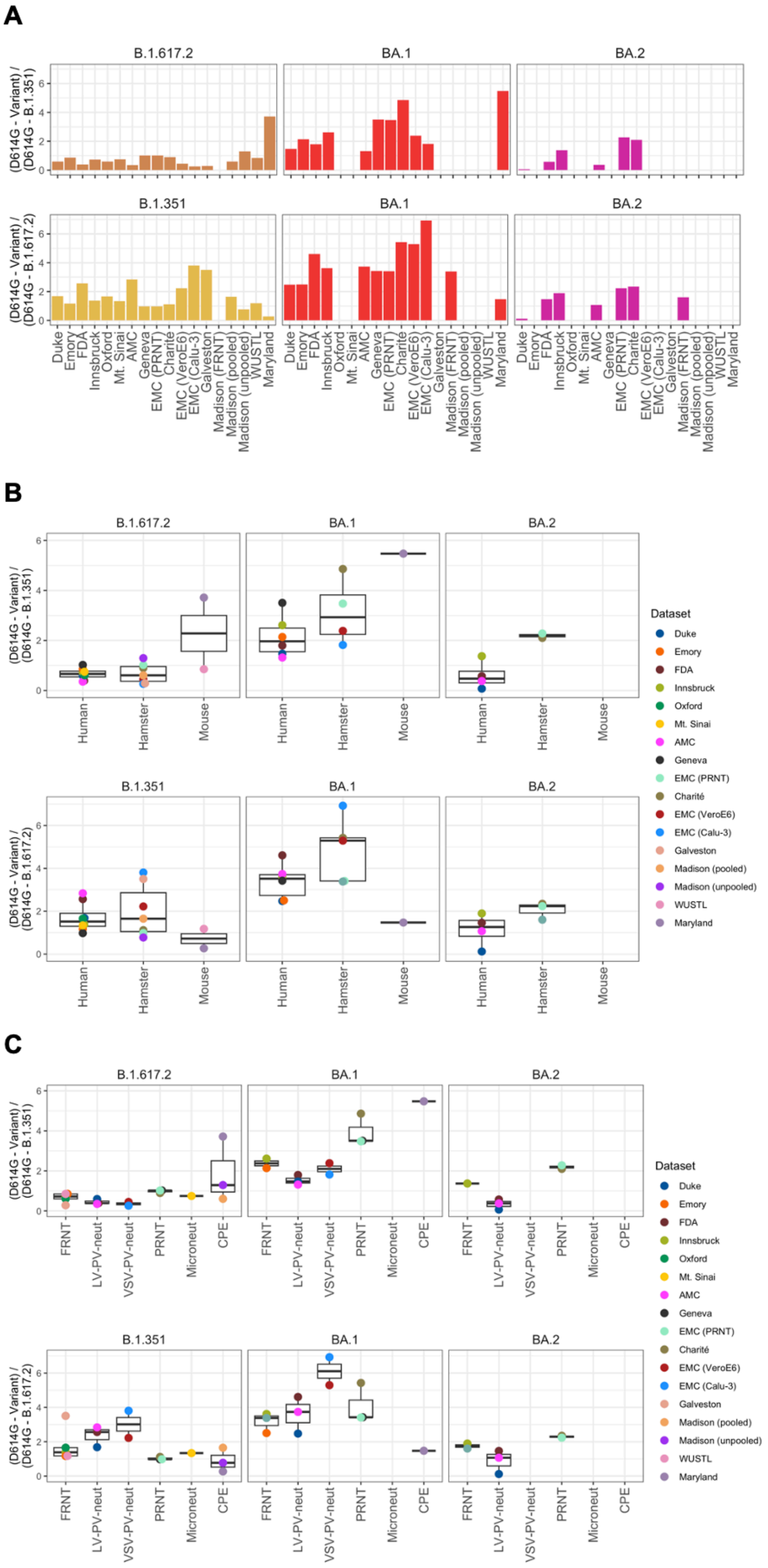
A
B.1.617.2
BA.1
BA.2
B.1.351
(D614G - Variant) / (D614G - B.1.351)
(D614G - Variant) / (D614G - B.1.617.2)
Duke
Emory
FDA
Innsbruck
Oxford
Mt. Sinai
AMC
Geneva
EMC (PRNT)
Charité
EMC (VeroE6)
EMC (Calu-3)
Galveston
Madison (FRNT)
Madison (pooled)
Madison (unpooled)
WUSTL
Maryland
B
Human
Hamster
Mouse
Dataset
C
FRNT
LV-PV-neut
VSV-PV-neut
PRNT
Microneut
CPE

**Figure S50: Relative distance within the pre-Omicron and Omicron variant clusters and between the pre-Omicron and Omicron variant clusters.** Distance within the pre-Omicron variants is measured by the following ratios of Euclidean distances between variants: (D614G - B.1.617.2) / (D614G - B.1.351), as well as (D614G - B.1.351) / (D614G - B.1.617.2). Relative distances between the pre-Omicron and Omicron variants were calculated as (D614G - BA.1) / (D614G - B.1.351), and (D614G - BA.1) / (D614G - B.1.617.2). Relative distances within the Omicron variants were calculated as (BA.1 - BA.2) / (D614G - B.1.351), and (BA.1 - BA.2) / (D614G - B.1.617.2). Ratios of distances were plotted A) per map, B) aggregated by animal model, C) aggregated by assay.

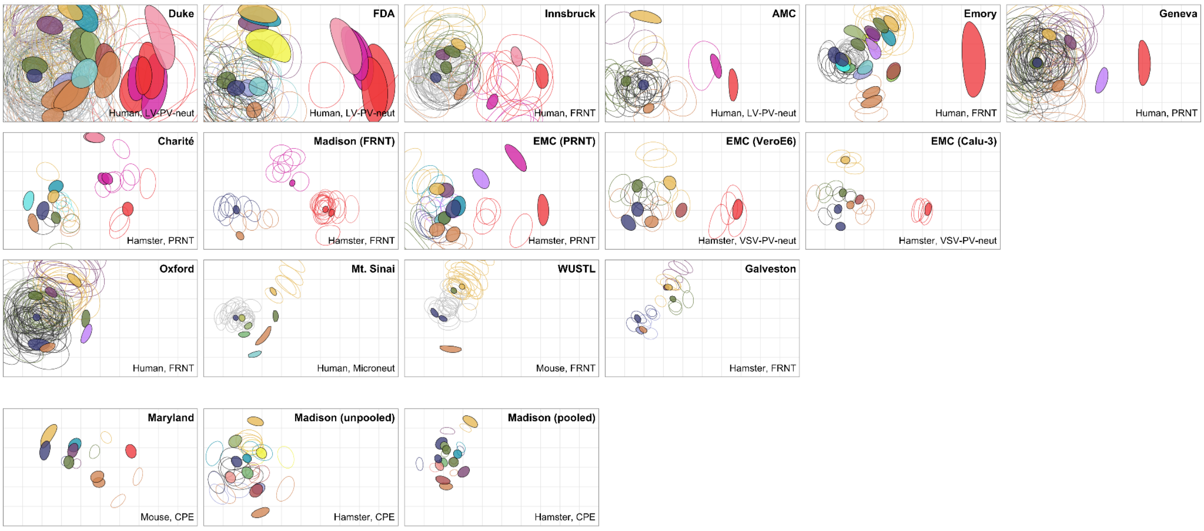


**Figure S51: Antigenic maps with triangulation regions.** Maps are colored by variant (see fig. S53). The colored region for each serum and variant point indicates the area which the variant or serum can occupy without the stress of the map increasing by more than 1% of the original map stress. Larger colored regions for a variant indicate that it is not coordinated well in antigenic space, possibly due to the small number of sera that resolve its position. Note the large colored regions for the Omicron variants in the Duke, FDA, Emory, and Geneva datasets, which also only have no or a relatively small number of sera in the space occupied by those variants.

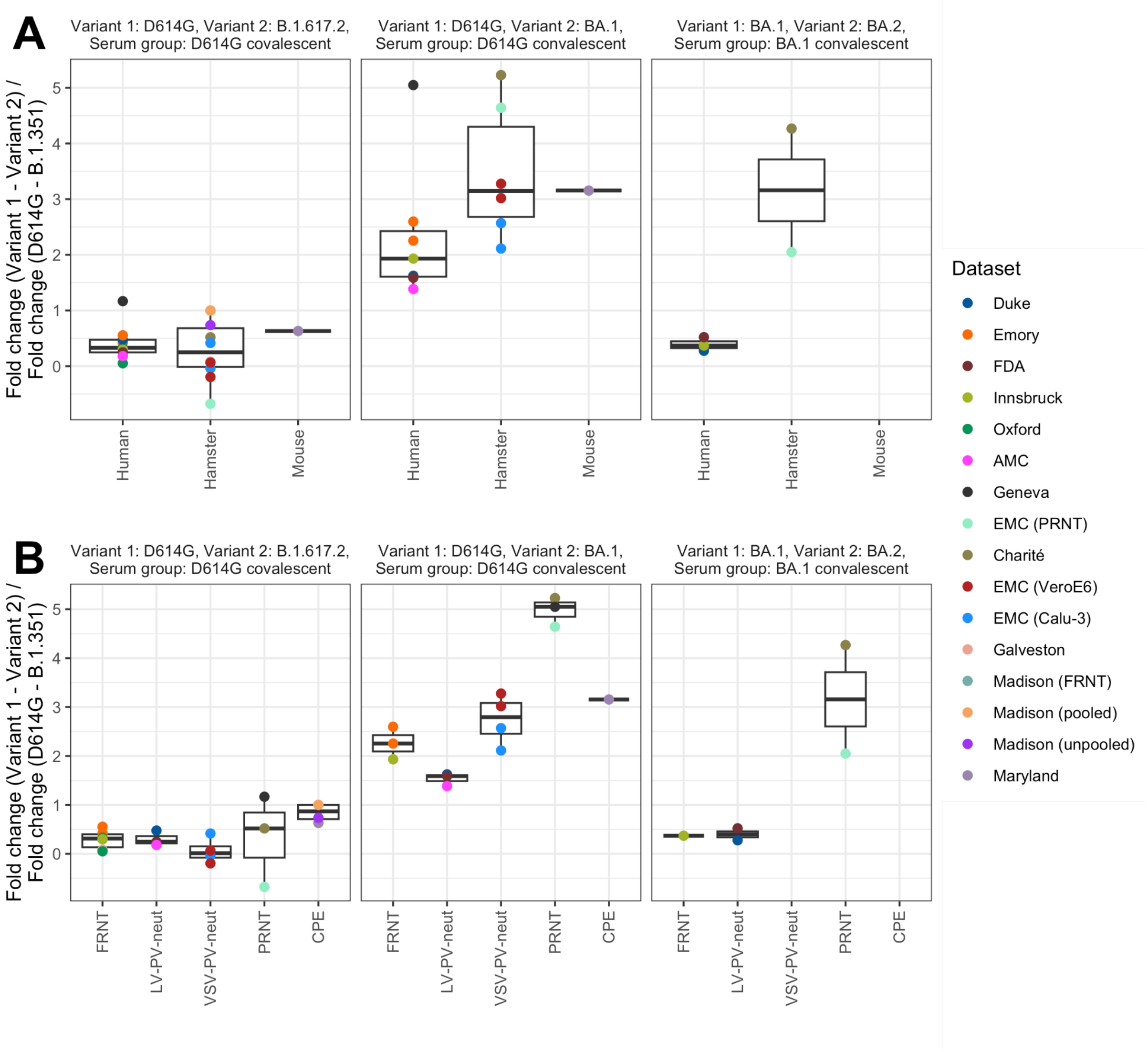


**Figure S52: Relative fold drop between pre-Omicron and Omicron variants.** Panels from left to right show the division of the fold change between the following pairs of variants: left: D614G and B.1.617.2 in D614G convalescent sera, divided by D614G and B.1.351 in D614G convalescent sera; middle: D614G and BA.1 in D614G convalescent sera and D614G and B.1.351 in D614G convalescent sera; right: BA.1 and BA.2 in BA.1 sera and D614G and B.1.351 in D614G convalescent sera. A) shows the results split by animal model, while B) shows the results split by assay.

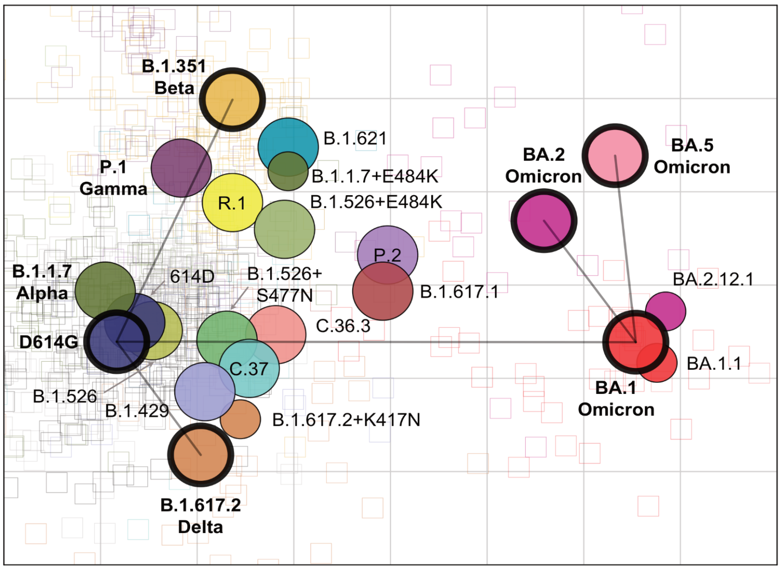


**Figure S53: Merged map colored by variant.** D614G, B.1.351, B.1.617.2, BA.1, BA.2 and BA.4/BA.5 are highlighted, and dashed lines are drawn to highlight their relative positions.

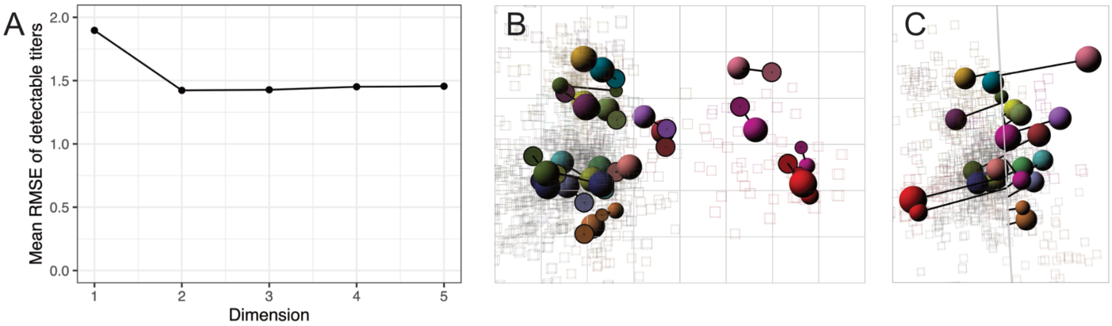


**Figure S54: Dimensionality tests and 3D map of the merged map.** A) RMSE of detectable titers in 1 to 5 dimensions compared to known titers. Per dimension, 100 repeats were performed wherein a map was constructed from 90% of all titers. For each run, the RMSE is calculated by comparing the titers predicted from the map against the known titers on the $\log_2$ scale. B, C) The map optimized in three dimensions, with arrows pointing to where each variant is placed in two dimensions. B) 'Side' view. C) 'Front' view. In panel A, no large reduction in error is observed when using more than two dimensions, suggesting that the two-dimensional representation fits the data acceptably well.

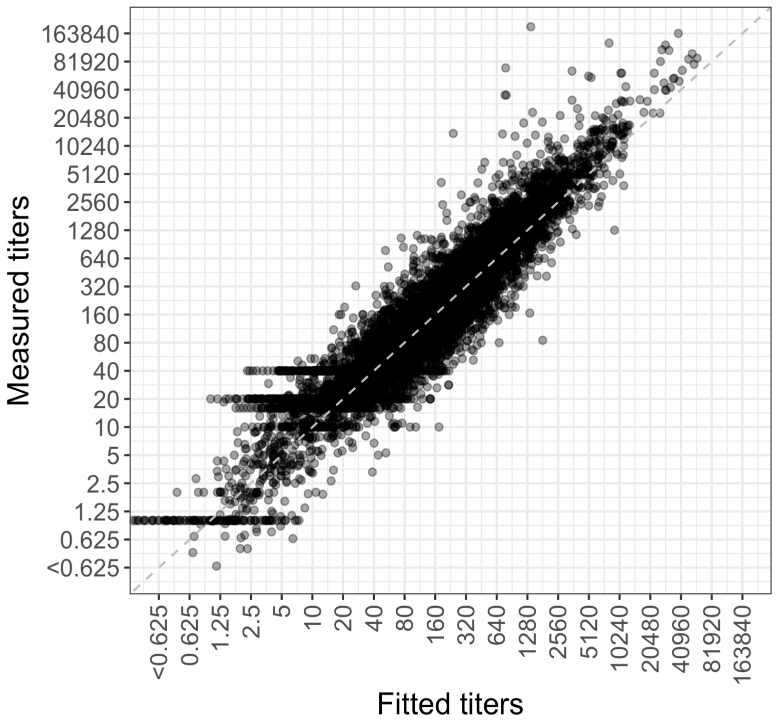


**Figure S55: Scatter plot of fitted titers and measured titers.** The fitted titers are determined from the distances in the two-dimensional antigenic map. Titers are plotted on the $\log_2$ scale. The dashed gray line is the line of best fit. Non-detectable measured titers were set to 1.

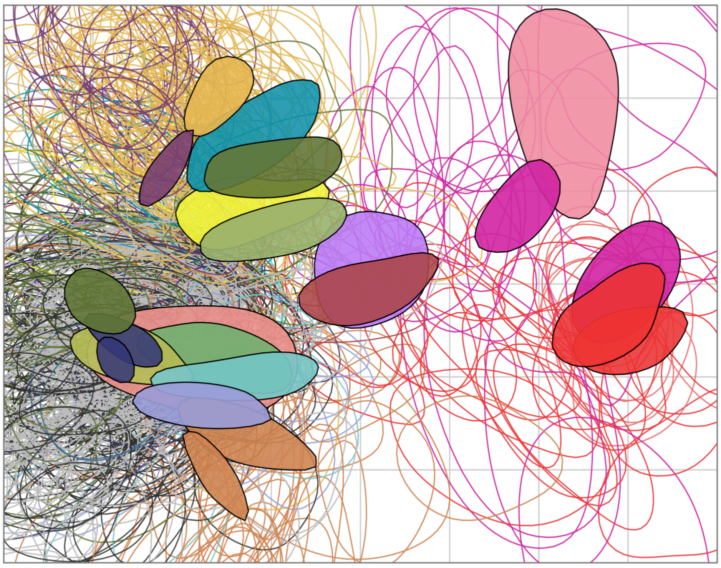

**Figure S56: Assessing the effect of the uncertainty in titers and variant reactivity for the antigenic map using 'smooth' bootstrap.** 100 bootstrap repeats were performed with 1000 optimisations per repeat. Noise was added to titers and to antigen reactivity. The noise added to the titers was normally distributed and had a standard deviation of 0.62, the noise added to the antigen reactivity a standard deviation of 0.4. Each colored region indicates the area in which 68% (one standard deviation) of the positional variation of a serum or variant position is captured. For color correspondence of variants and sera, refer to fig. S53.

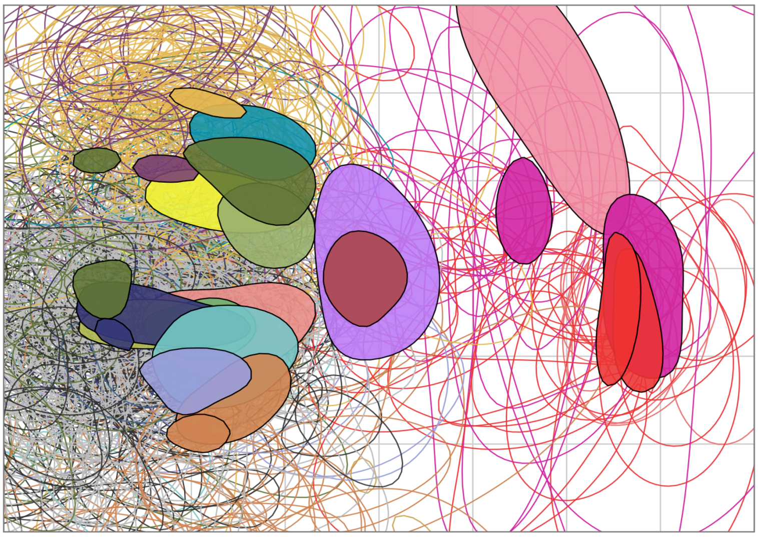

**Figure S57: Assessing the robustness of the antigenic map to the inclusion of particular titer measurements.** 100 bootstrap repeats were performed with 1000 optimisations each, where a random sample of titers were taken with replacement. Bootstrapping was applied to variants and sera. Each colored region indicates the area in which 68% (one standard deviation) of the positional variation of a serum or variant position is captured. For color correspondence of variants and sera, refer to fig. S53.

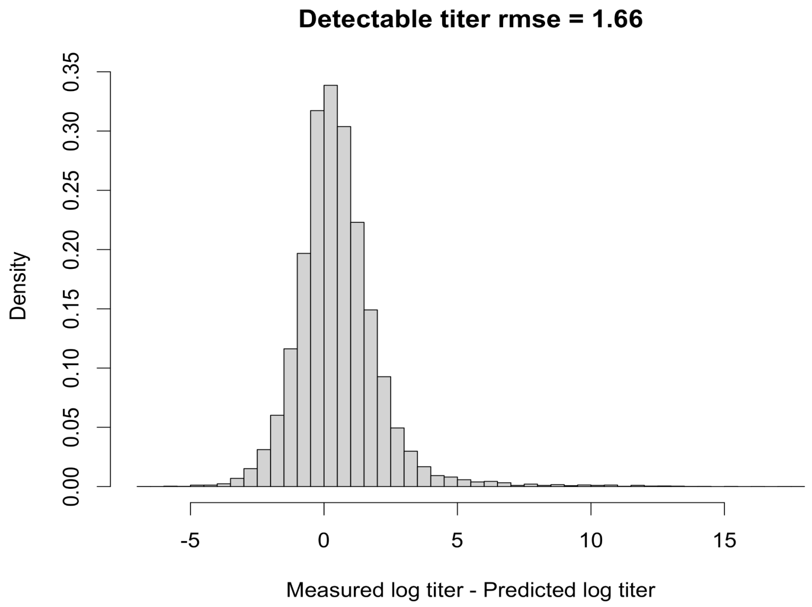


**Figure S58: Histogram of differences between measured and predicted $\log_2$ titers.** 100 replicates were performed where 10% of the data were omitted, a new map constructed with 1000 optimisations, and the missing titers were estimated from the new map. The difference between predicted and measured $\log_2$ titers is shown. Only detectable titers were considered. Predicted titers are on average 1.66 $\log_2$ lower than measured titers. Assuming a mean of 0 to account for the systematic bias of titrations between repeats, the standard deviation is 1.58 on the $\log_2$ scale.

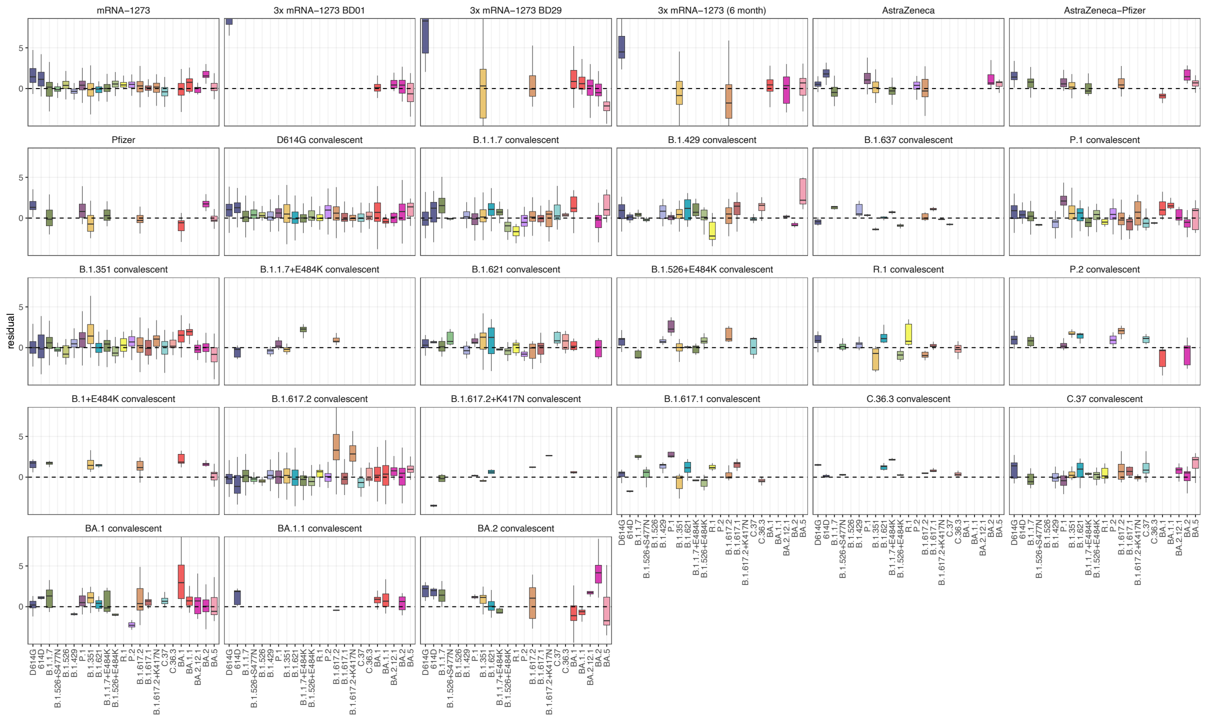


**Figure S59: Difference between measured and predicted $\log_2$ titers split by serum group and variant.** 100 replicates were performed where 10% of the data were omitted, a new map constructed with 1000 optimisations, and the missing titers were estimated from the new map. The difference between predicted and measured $\log_2$ titers is shown split by variant and serum group. Only detectable titers were considered. The box plot indicates the median and 25$^{th}$ and 75$^{th}$ percentile.

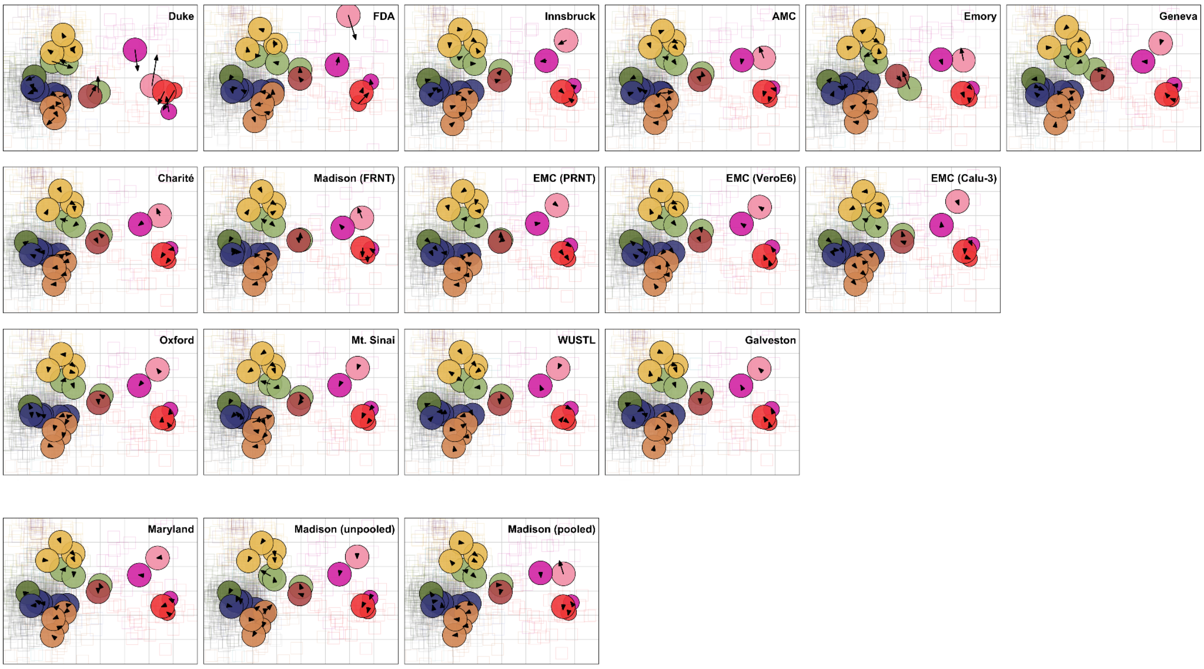


**Figure S60: Sensitivity of merged map to inclusion of different datasets.** In each panel, one of the 18 datasets was omitted from the construction of the merged map. The name of the omitted dataset is indicated in the top right corner of each panel. The arrows point from the position of a variant in the leave-one-out map to the corresponding position of the same variant in the merged map constructed from all 18 datasets.

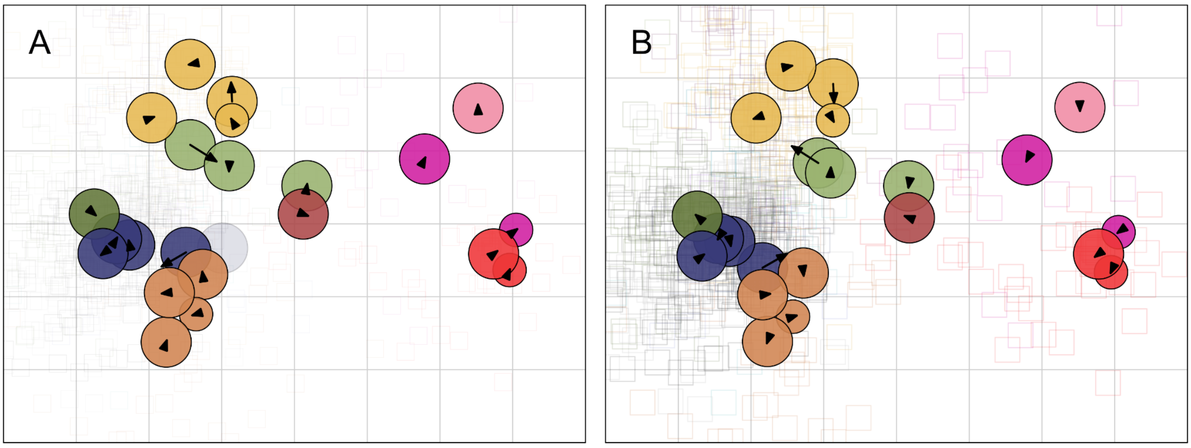


**Figure S61: Sensitivity of merged map to inclusion of different individual datasets, effect of removing datasets generated using CPE assays.** A) The arrows point from the position of a variant in the merged map without the CPE assay datasets to the corresponding position of the same variant in the merged map constructed from all 18 individual maps. B) The arrows point from the position of a variant in the merged map constructed from all datasets to the corresponding position of the same variant in the merged map constructed without the CPE assay datasets.

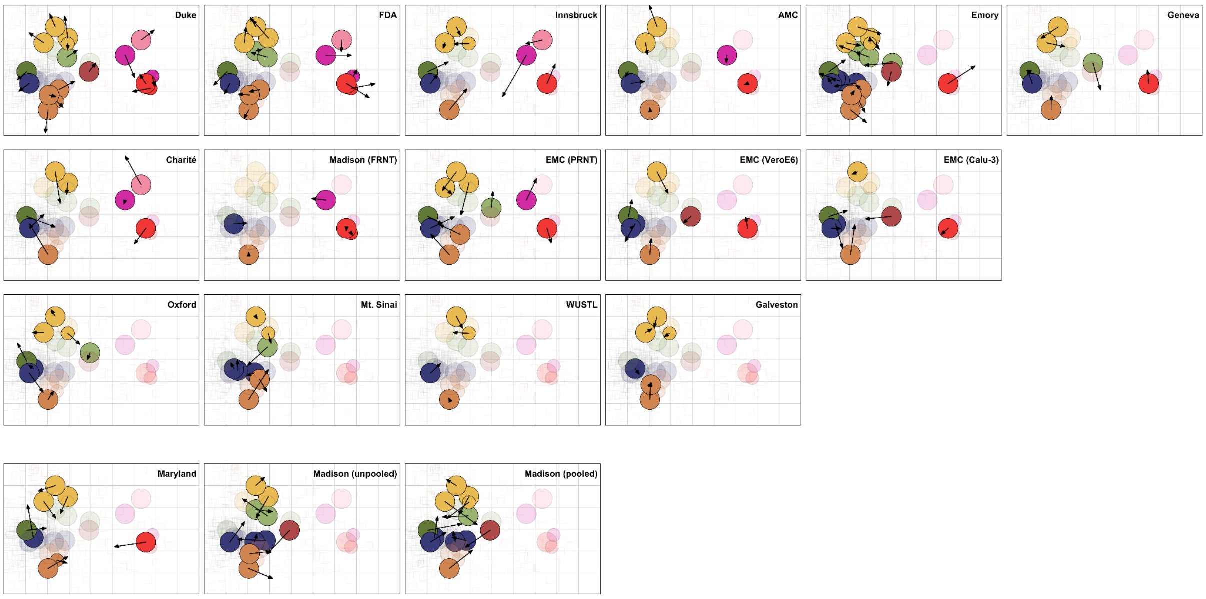


**Figure S62: Comparison of the merged map to individual maps.** Arrows point from the position of a variant in the merged map to the corresponding position of the same variant in each individual map. The individual map is indicated in the top right of the panel.

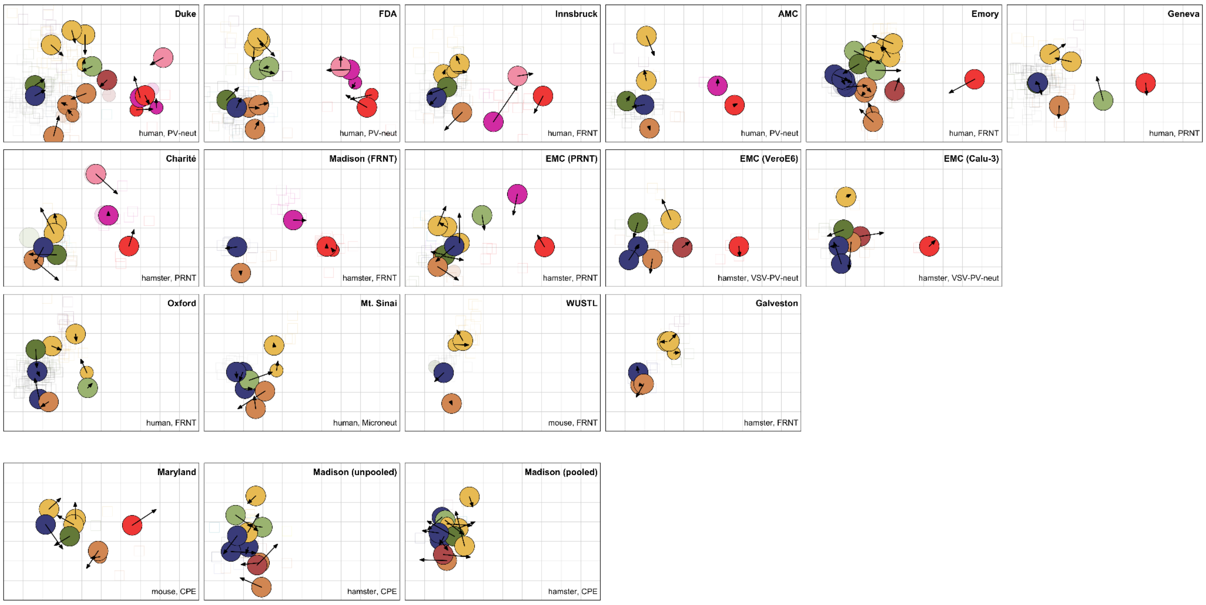


**Figure S63: Comparison of individual maps to the merged map.** Arrows point from the position of a variant in each individual map to the corresponding position of the same variant in the merged map. The individual map is indicated in the top right of the panel.

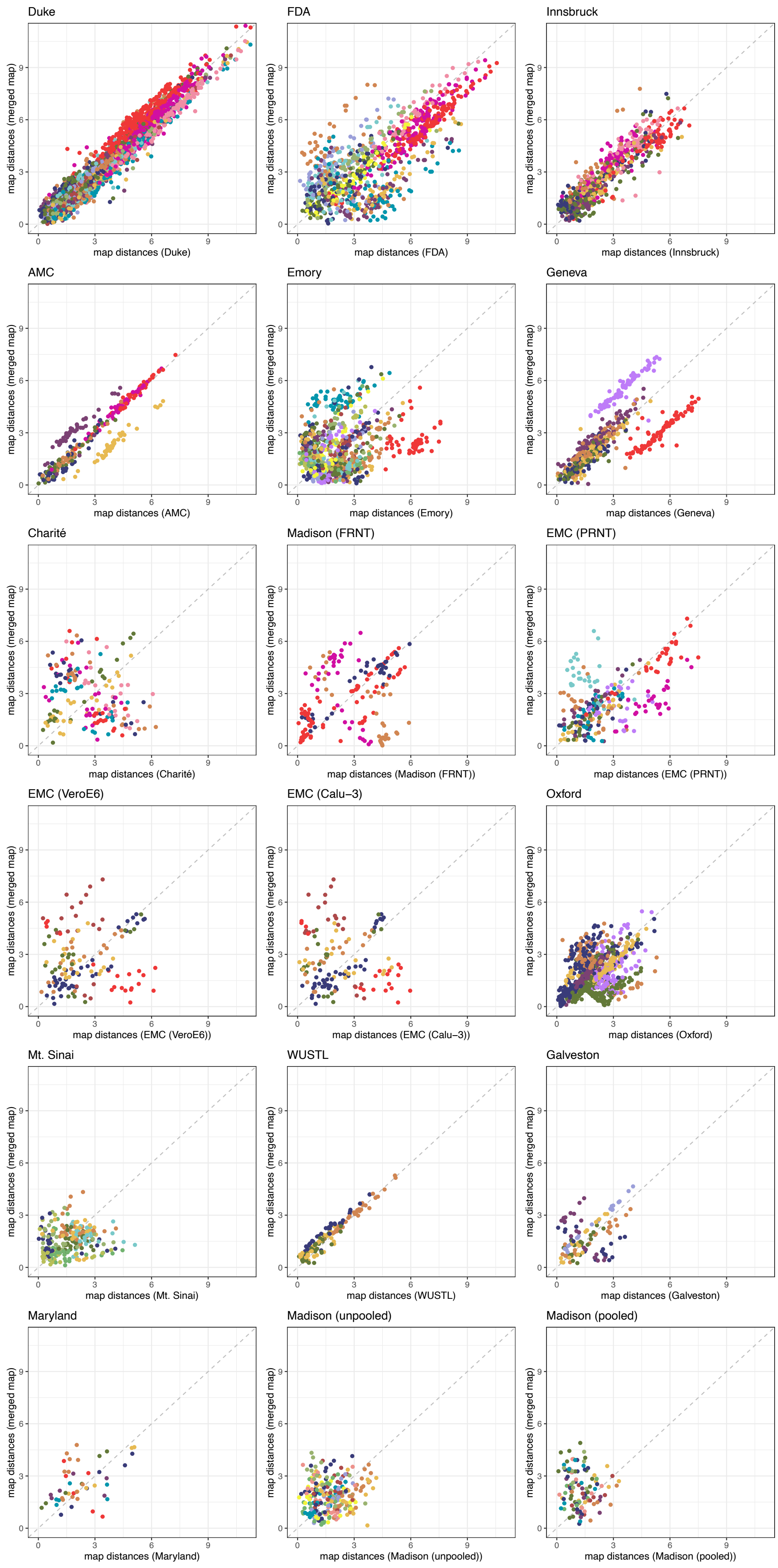
Duke
FDA
Innsbruck
AMC
Emory
Geneva
Charité
Madison (FRNT)
EMC (PRNT)
EMC (VeroE6)
EMC (Calu−3)
Oxford
Mt. Sinai
WUSTL
Galveston
Maryland
Madison (unpooled)
Madison (pooled)
map distances (merged map)
map distances (Duke)
map distances (FDA)
map distances (Innsbruck)
map distances (AMC)
map distances (Emory)
map distances (Geneva)
map distances (Charité)
map distances (Madison (FRNT))
map distances (EMC (PRNT))
map distances (EMC (VeroE6))
map distances (EMC (Calu−3))
map distances (Oxford)
map distances (Mt. Sinai)
map distances (WUSTL)
map distances (Galveston)
map distances (Maryland)
map distances (Madison (unpooled))
map distances (Madison (pooled))
0
3
6
9

**Figure S64: Scatter plot of map distances in the merged map and map distances in each individual map.** Scatter plots are colored by variant. Six maps (Duke, FDA, Innsbruck, AMC, Geneva, and WUSTL) show a good correlation between the map distances in the merged map compared to the individual maps. In six maps (Emory, Madison (FRNT), EMC (PRNT), EMC (VeroE6), Oxford, and Galveston), there is a within-variant correlation of distances in the merged map compared to the individual maps, possibly resulting from the differences in scale between the merged and the individual maps, that affect variants differently. The map distances in the remaining individual maps are not well correlated with the distances in the merged map. For the CPE assay datasets (Maryland, Madison (unpooled), and Madison (pooled), this is as expected, based on the different amount and rank order of fold change

| Dataset | Lab | Species | Assay (Antigen type / assay / cell type) | Sera | Serum information | Antigens | Titer units | Refer-ence |
|---|---|---|---|---|---|---|---|---|
| Duke | David Montefiori, Duke | Human | Lentivirus pseudotype Neutralization HEK293T-ACE2 | B.1.617.2 conv. (n=26, taken ≥28 days and <60 days post infection), B.1.1.7 conv. (n=13, taken ≥28 days and <60 days post infection), B.1.526+E484K conv. (n=6, taken ≥28 days and <60 days, or ≥60 days post infection), B.1.637 conv. (n=2, taken ≥28 days and <60 days post infection), P.1 conv. (n=13, taken ≥28 days and <60 days post infection), mRNA-1273 2x vax. (n=30, taken 4 weeks post vax.), D614G conv. (n=13, taken ≥28 days and <60 days post infection), B.1.351 conv. (n=24, taken ≥28 days and <60 days post infection), C.37 conv. (n=7, taken ≥28 days and <60 days post infection), BA.1 conv. (n=4, taken ≥28 days and <60 days post infection), mRNA-1273 3x vax (n=26, 4 weeks post vax). | N/A | D614G, B.1.1.7, B.1.1.7+E484K, P.1, B.1.351, B.1.429, B.1.526+E484K, B.1.617.1, B.1.617.2, B.1.617.2+K417N, B.1.617.2 (AY.1)+K417N, B.1.617.2 (AY.2)+K417N, B.1.617.2 (AY.3)+E484Q, B.1.621, C.37, BA.1, BA.1.1, BA.2, BA.2.12.1, BA.3, BA.4/BA.5 | NT50 | (*3*) |
| Emory | Mehul Suthar, Emory | Human | Live virus FRNT VeroE6-TMPRSS2 | D614G conv. (31 to 60 days after symptom onset, n=17), B.1.351 conv. (28 days after diagnosis, n=17), B.1.617.2 conv. (2 to 15 days after symptom onset, n=14), mRNA-1273 2x vax. (35 to 51 days post second dose) | N/A | 614D, B.1.351, R.1, A.23.1, C.37, D614G, B.1.617.2, B.1.617.2+K417N, B.1.617.1, B.1.525, B.1.526, B.1.526+E484K, B.1.1.7, B.1.1.7+E484K, B.1.621, P.1, B.1.630, B.1.526+S477N, P.2, B.1.429, BA.1 | NT50 | Unpubl ished |

| | | | | | | | | |
|---|---|---|---|---|---|---|---|---|
| FDA | Carol Weiss, FDA | Human | Lentivirus pseudotype Neutralization 293T-ACE2-TMPRSS2 | D614G conv. (n=25), B.1.429 conv. (n=5), B.1.1.7 conv. (n=10), C.37 conv. (n=10), P.1 conv. (n=4), B.1.351 conv. (n=7), B.1.617.2 conv. (n=15), BA.1 conv. (n=2), B.1.526+E484K conv (n=1). Sera taken 8-51 days post infection, mean=28 days). | N/A | B.1.1.7, B.1.351, B.1.429, B.1.617.2, P.1, C.37, B.1.621, B.1.526+E484K, D614G, BA.1, BA.1.1, BA.2, BA.2.12.1, BA.5, R.1 | NT50 | (*8*) |
| Oxford | Gavin Screaton, Oxford | Human | Live virus FRNT Vero | BNT162b2 2x vax (n=24, taken 7 to cccc15 days post second dose), AstraZeneca 2x (n=25, taken 14 or 28 days past second dose), D614G conv. (n=34, taken at least 4 to 9 weeks post symptom onset), B.1.351 conv. (n=14), B.1.1.7 conv. (n=18, taken at least 14 days post symptom onset), P.1 conv (n=14). | N/A | 614D, B.1.1.7, B.1.351, P.1, B.1.617.2, P.2, B.1.1.7+E484K, D614G | NT50 | (*35*) |
| Mt. Sinai | Florian Krammer, Mt. Sinai School of Medicine | Human | Live-virus Microneutralization VeroE6 | BNT162b2 2x vax (n=15) and mRNA-1273 2x vax (n=15), taken 2 to 29 days post second dose, B.1.351 conv (n=5), taken on average 28 days post-diagnosis (min: 19 days, max: 46 days). | N/A | C.37, B.1.351, B.1.1.7+E484K, B.1.617.2, B.1.526+E484K, B.1.526, B.1.526+S477N, 614D | NT50 | (*24*) |
| Innsbruck | Janine Kimpel, Medical University of Innsbruck | Human | Live-virus FRNT Vero-TMPRSS2-ACE2 | B.1.617.2 conv. (n=5, mean 10 weeks post infection), B.1.1.7 conv. (n=9, mean 22 weeks post infection), B.1.351 conv. (n=6, 1-4 weeks post infection), mRNA-1273 2x vax. (n=10, mean 20 weeks post vax), AstraZeneca 2x vax. (n=10, mean 4.3 weeks post vax), AstraZeneca-BNT162b2 vax. (n=10, mean 4.3 weeks post vax), BNT162b2 2x vax. (n=11, mean 4.3 weeks post vax), BA.1 conv. (n=8, mean 2.4 weeks post infection), BA.2 conv. (n=9, mean 2.3 weeks post infection) | N/A | D614G, B.1.1.7, B.1.1.7+E484K, P.1, B.1.351, B.1.617.2, BA.1, BA.2, BA.5 | NT50 | (*12*) |

| | | | | | | | | |
|---|---|---|---|---|---|---|---|---|
| Geneva | Isabella Eckerle, University of Geneva | Human | Live-virus PRNT VeroE6 | D614G conv. (n=35, 25-37 days post symptom onset), B.1.1.7 conv. (n=12, 8-42 days post symptom onset), B.1.351 conv. (n=8, 3-98 days post symptom onset), P.1 conv. (n=10, 7-137 days post symptom onset), B.1.617.2 conv. (n=10, 9-118 days post symptom onset), mRNA-1273 2x vax. (n=16, 8 weeks post second dose) | N/A | D614G, B.1.1.7, B.1.351, P.1, B.1.617.2, P.2, BA.1 | NT90 | (*7*) |
| AMC | Rogier Sanders, Amsterdam Medical Centre | Human | Lentivirus pseudotype Neutralization HEK293T/ACE2 | D614G conv. (n=21, 30-45 days post symptom onset), B.1.1.7 conv. (n=8, 24-58 days post symptom onset), B.1.351 conv. (n=4, 30-63 days post symptom onset), P.1 conv. (n=3, 38-55 days post symptom onset), B.1.617.2 conv. (n=8, 36-52 days post symptom onset), BA.1 conv. (n=1, 75 days post symptom onset), BA.2 conv. (n=1, 41 days post symptom onset) | N/A | D614G, B.1.1.7, B.1.351, P.1, B.1.617.2, BA.1, BA.2 | IU/ml | (*11*) |
| WUSTL | Michael Diamond, Washington University in St. Louis | Mouse (129S2, female, 7-9 weeks) | Live-virus FRNT VeroE6-TMPRSS2 | mRNA-1273 vax. (n=18), mRNA-1273.351 vax (n=18). Sera taken at 6 weeks post infection. | Inoculated and boosted 3 weeks apart, 0.25 µg. | D614G, N501Y+D614G, B.1.1.7+E484K, B.1.351, B.1.617.2 | NT50 | (*10*) |
| Maryland | Matthew Frieman, University of Maryland | Mouse (female BALB/c, 7-9 weeks) | Live-virus CPE VeroE6-TMPRSS2 | D614G vax., B.1.1.7 vax., B.1.351 vax., P.1 vax., B.1.617.2 vax., B.1.617.2+K417N vax.<br>1 serum pool per serum group. Sera taken at day 21. | Inoculated and boosted 14 days apart. Inoculation with spike nanoparticles with 5 µg Matrix-M adjuvant. Sera from 20 animals | 614D, B.1.1.7, B.1.351, P.1, B.1.621, B.1.617.2, B.1.617.2+K417N, BA.1 | NT99 | (*26*) |

| | | | | | | | | |
|---|---|---|---|---|---|---|---|---|
| | | | | | were pooled. | | | |
| Madison (pooled) | Yoshihior Kawaoka, University of Wisconsin-Madison | Hamster | Live-virus CPE VeroE6-TMPRSS2 | B.1.617.1 vax., B.1.617.2 vax., D614G vax., P.1 vax., B.1.351 vax., B.1.1.7 vax., C.36.3 vax., B.1.621 vax., 614D vax. 1 serum pool per serum group, except for D614G conv., where 2 pools were used. Sera taken at 4 weeks post infection. | Inoculated with $10^3$ pfu live virus. Sera pooled from 3 individuals. | B.1.617.1, B.1.617.2, D614G, P.1, B.1.351, B.1.1.7, B.1.1.7+E484K, B.1.526+E484K, C.36.3, B.1.526+S477N, 614D, B.1.621 | NT99 | Unpublished |
| Madison (unpooled) | Yoshihior Kawaoka, University of Wisconsin-Madison | Hamster | Live-virus CPE VeroE6-TMPRSS2 | B.1.351 vax. (n=6), D614G vax. (n=6), B.1.617.1 vax. (n=3), B.1.429 vax. (n=3), B.1.617.2 vax. (n=3), B.1.621 vax. (n=3), R.1 vax. (n=3). Sera taken at 4 weeks post infection. | Inoculated with $10^3$ pfu live virus. | B.1.351, D614G, B.1.617.2, B.1.621, B.1.617.1, B.1.526+E484K, C.36.3, B.1.526+S477N, B.1.429, R.1 | NT99 | Unpublished |
| Madison (FRNT) | Yoshihior Kawaoka, University of Wisconsin-Madison | Hamster | Live-virus FRNT VeroE6-TMPRSS2 | D614G conv. (n=7), B.1.617.2 conv. (n=4), BA.1 conv. (n=9), BA.1.1 conv. (n=10), BA.2 conv. (n=7). Sera taken at 4 weeks post infection. | Inoculated with $10^3$ pfu live virus. | 614D, B.1.617.2, BA.1, BA.1.1, BA.2 | NT50 | Unpublished |
| Galveston | Pei-Yong Shi, University of Texas, Galveston | Hamster (Syrian golden hamster, male, 4-6 weeks) | Live-virus FFRNT VeroE6 | D614G conv., B.1.1.7+E484K conv., B.1.351 conv., P.1 conv., B.1.429 conv. 4 sera per serum group. Sera taken at 28 days post infection. | $10^6$ pfu live virus | 614D, B.1.1.7+E484K, B.1.351, P.1, B.1.429, B.1.617.2 | NT50 | (*9*) |
| EMC (PRNT) | Bart Haagmans, Erasmus Medical Centre (EMC) | Hamster (Syrian golden hamster, female, 6 weeks) | Live-virus PRNT Calu-3 | D614G conv., B.1.1.7 conv., B.1.351 conv., P.2 conv., B.1.617.2 conv., B.1.621 conv., BA.1 conv. 4 sera per serum group. Sera taken at 26 days post infection. | $1.0 \times 10^5$ pfu (614G, Alpha, Beta, Gamma, Zeta, Mu) or $5.0 \times 10^4$ pfu (Delta, Omicron), nasally. | D614G, B.1.1.7, B.1.351, P.1, P.2, B.1.617.2, B.1.621, BA.1, C.37, AY.4.2, BA.2 | NT50 | (*4*) |

| | | | | | | | | |
|---|---|---|---|---|---|---|---|---|
| EMC (Calu-3) | Bart Haagmans, Erasmus Medical Centre (EMC) | Hamster (Syrian golden hamster, female, 6 weeks) | VSV pseudotypes Neutralization Calu-3 | D614G conv., B.1.1.7 conv., B.1.351 conv., B.1.617.2 conv., BA.1 conv. | Same sera as ‘EMC (PRNT)’ dataset | 614D, D614G, B.1.1.7, B.1.351, B.1.617.2, B.1.617.1, BA.1 | NT50 | (*4*) |
| EMC (VeroE6) | Bart Haagmans, Erasmus Medical Centre (EMC) | Hamster (Syrian golden hamster, female, 6 weeks) | VSV pseudotypes Neutralization VeroE6 | D614G conv., B.1.1.7 conv., B.1.351 conv., B.1.617.2 conv., BA.1 conv. | Same sera as ‘EMC (PRNT)’ dataset | 614D, D614G, B.1.1.7, B.1.351, B.1.617.2, B.1.617.1, BA.1 | NT50 | (*4*) |
| Charité | Christian Drosten/Victor Corman, Charité - Universitätsmedizin Berlin | Hamster (Syrian golden hamster, female, 6 weeks) | Live-virus PRNT VeroE6 | BA.2 conv., B.1.1.7 conv., B.1.351 conv., B.1.617.2 conv., BA.1 conv., B.1+E484K conv., D614G conv. 3 sera per serum group. Sera taken at day 36. | Infection with 1000 pfu at day 0, infection with $1x10^6$ pfu at day 21. | B.1.1.7, BA.1, BA.2-12, BA.2, BA.4, BA.5, B.1.351, B.1.617.2, B.1.621, D614G, B.1+E484K | NT90 | (*25*) |

**Table S1: Detailed description of datasets.** Abbreviations for the ‘Sera’ column are: vax.: vaccinated, conv.: convalescent. Abbreviations for the ‘Assay’ column are: FRNT: Focus forming neutralization assay, PRNT: Plaque reduction neutralization assay, CPE: Cytopathic effect assays (limiting dilution assay).

4. A. Z. Mykytyn, M. Rissmann, A. Kok, M. E. Rosu, D. Schipper, T. I. Breugem, P. B. van den Doel, F. Chandler, T. Bestebroer, M. de Wit, M. E. van Royen, R. Molenkamp, B. B. O. Munnink, R. D. de Vries, C. GeurtsvanKessel, D. J. Smith, M. P. G. Koopmans, B. Rockx, M. M. Lamers, R. Fouchier, B. L. Haagmans, Antigenic cartography of SARS-CoV-2 reveals that Omicron BA.1 and BA.2 are antigenically distinct. *Science Immunology*. **0**, eabq4450.

5. W. Dejnirattisai, J. Huo, D. Zhou, J. Zahradník, P. Supasa, C. Liu, H. M. E. Duyvesteyn, H. M. Ginn, A. J. Mentzer, A. Tuekprakhon, R. Nutalai, B. Wang, A. Dijokaite, S. Khan, O. Avinoam, M. Bahar, D. Skelly, S. Adele, S. A. Johnson, A. Amini, T. G. Ritter, C. Mason, C. Dold, D. Pan, S. Assadi, A. Bellass, N. Omo-Dare, D. Koeckerling, A. Flaxman, D. Jenkin, P. K. Aley, M. Voysey, S. A. Costa Clemens, F. G. Naveca, V. Nascimento, F. Nascimento, C. Fernandes da Costa, P. C. Resende, A. Pauvolid-Correa, M. M. Siqueira, V. Baillie, N. Serafin, G. Kwatra, K. Da Silva, S. A. Madhi, M. C. Nunes, T. Malik, P. J. M. Openshaw, J. K. Baillie, M. G. Semple, A. R. Townsend, K.-Y. A. Huang, T. K. Tan, M. W. Carroll, P. Klenerman, E. Barnes, S. J. Dunachie, B. Constantinides, H. Webster, D. Crook, A. J. Pollard, T. Lambe, OPTIC Consortium, ISARIC4C Consortium, N. G. Paterson, M. A. Williams, D. R. Hall, E. E. Fry, J. Mongkolsapaya, J. Ren, G. Schreiber, D. I. Stuart, G. R. Screaton, SARS-CoV-2 Omicron-B.1.1.529 leads to widespread escape from neutralizing antibody responses. *Cell*. **185**, 467–484.e15 (2022).

doi:10.3390/vaccines9010013.

20. C. Liu, H. M. Ginn, W. Dejnirattisai, P. Supasa, B. Wang, A. Tuekprakhon, R. Nutalai, D. Zhou, A. J. Mentzer, Y. Zhao, H. M. E. Duyvesteyn, C. López-Camacho, J. Slon-Campos, T. S. Walter, D. Skelly, S. A. Johnson, T. G. Ritter, C. Mason, S. A. C. Clemens, F. G. Naveca, V. Nascimento, F. Nascimento, C. F. da Costa, P. C. Resende, A. Pauvolid-Correa, M. M. Siqueira, C. Dold, N. Temperton, T. Dong, A. J. Pollard, J. C. Knight, D. Crook, T. Lambe, E. Clutterbuck, S. Bibi, A. Flaxman, M. Bittaye, S. Belij-Rammerstorfer, S. C. Gilbert, T. Malik, M. W. Carroll, P. Klenerman, E. Barnes, S. J. Dunachie, V. Baillie, N. Serafin, Z. Ditse, K. Da Silva, N. G. Paterson, M. A. Williams, D. R. Hall, S. Madhi, M. C. Nunes, P. Goulder, E. E. Fry, J. Mongkolsapaya, J. Ren, D. I. Stuart, G. R. Screaton, Reduced neutralization of SARS-CoV-2 B.1.617 by vaccine and convalescent serum. *Cell*. **184** (2021), pp. 4220–4236.e13.

21. J. M. Fonville, S. H. Wilks, S. L. James, A. Fox, M. Ventresca, M. Aban, L. Xue, T. C. Jones, N. M. H. Le, Q. T. Pham, N. D. Tran, Y. Wong, A. Mosterin, L. C. Katzelnick, D. Labonte, T. T. Le, G. van der Net, E. Skepner, C. A. Russell, T. D. Kaplan, G. F. Rimmelzwaan, N. Masurel, J. C. de Jong, A. Palache, W. E. P. Beyer, Q. M. Le, T. H.

Nguyen, H. F. L. Wertheim, A. C. Hurt, A. D. M. E. Osterhaus, I. G. Barr, R. A. M. Fouchier, P. W. Horby, D. J. Smith, Antibody landscapes after influenza virus infection or vaccination. *Science*. **346**, 996–1000 (2014).

22. D. J. Smith, A. S. Lapedes, J. C. de Jong, T. M. Bestebroer, G. F. Rimmelzwaan, A. D. M. Osterhaus, R. A. M. Fouchier, Mapping the Antigenic and Genetic Evolution of Influenza Virus. *Science*. **305** (2004), pp. 371–376.

23. M. M. DeGrace, E. Ghedin, M. B. Frieman, F. Krammer, A. Grifoni, A. Alisoltani, G. Alter, R. R. Amara, R. S. Baric, D. H. Barouch, J. D. Bloom, L.-M. Bloyet, G. Bonenfant, A. C. M. Boon, E. A. Boritz, D. L. Bratt, T. L. Bricker, L. Brown, W. J. Buchser, J. M. Carreño, L. Cohen-Lavi, T. L. Darling, M. E. Davis-Gardner, B. L. Dearlove, H. Di, M. Dittmann, N. A. Doria-Rose, D. C. Douek, C. Drosten, V.-V. Edara, A. Ellebedy, T. P. Fabrizio, G. Ferrari, W. M. Fischer, W. C. Florence, R. A. M. Fouchier, J. Franks, A. García-Sastre, A. Godzik, A. S. Gonzalez-Reiche, A. Gordon, B. L. Haagmans, P. J. Halfmann, D. D. Ho, M. R. Holbrook, Y. Huang, S. L. James, L. Jaroszewski, T. Jeevan, R. M. Johnson, T. C. Jones, A. Joshi, Y. Kawaoka, L. Kercher, M. P. G. Koopmans, B. Korber, E. Koren, R. A. Koup, E. B. LeGresley, J. E. Lemieux, M. J. Liebeskind, Z. Liu, B. Livingston, J. P. Logue, Y. Luo, A. B. McDermott, M. J. McElrath, V. A. Meliopoulos, V. D. Menachery, D. C. Montefiori, B. Mühlemann, V. J. Munster, J. E. Munt, M. S. Nair, A. Netzl, A. M. Niewiadomska, S. O'Dell, A. Pekosz, S. Perlman, M. C. Pontelli, B. Rockx, M. Rolland, P. W. Rothlauf, S. Sacharen, R. H. Scheuermann, S. D. Schmidt, M. Schotsaert, S. Schultz-Cherry, R. A. Seder, M. Sedova, A. Sette, R. S. Shabman, X. Shen, P.-Y. Shi, M. Shukla, V. Simon, S. Stumpf, N. J. Sullivan, L. B. Thackray, J. Theiler, P. G. Thomas, S. Trifkovic, S. Türeli, S. A. Turner, M. A. Vakaki, H. van Bakel, L. A. VanBlargan, L. R. Vincent, Z. S. Wallace, L. Wang, M. Wang, P. Wang, W. Wang, S. C. Weaver, R. J. Webby, C. D. Weiss, D. E. Wentworth, S. M. Weston, S. P. J. Whelan, B. M. Whitener, S. H. Wilks, X. Xie, B. Ying, H. Yoon, B. Zhou, T. Hertz, D. J. Smith, M. S. Diamond, D. J. Post, M. S. Suthar, Defining the risk of SARS-CoV-2 variants on immune protection. *Nature*. **605**, 640–652 (2022).

24. J. M. Carreño, H. Alshammary, G. Singh, A. Raskin, F. Amanat, A. Amoako, A. S. Gonzalez-Reiche, A. van de Guchte, P. Study Group, K. Srivastava, E. M. Sordillo, D. N. Sather, H. van Bakel, F. Krammer, V. Simon, Evidence for retained spike-binding and neutralizing activity against emerging SARS-CoV-2 variants in serum of COVID-19 mRNA vaccine recipients. *EBioMedicine*. **73**, 103626 (2021).

25. B. Mühlemann, J. Trimpert, F. Walper, M. L. Schmidt, S. Schroeder, L. M. Jeworowski, J. Beheim-Schwarzbach, T. Bleicker, D. Niemeyer, J. M. Adler, R. M. Vidal, C. Langner, D. Vladimirova, D. J. Smith, M. Voß, L. Paltzow, C. M. Christophersen, R. Rose, A. Krumbholz, T. C. Jones, V. M. Corman, C. Drosten, Antigenic cartography using variant-specific hamster sera reveals substantial antigenic variation among Omicron subvariants. *bioRxiv* (2023), p. 2023.07.02.547076.

26. J. Logue, R. M. Johnson, N. Patel, B. Zhou, S. Maciejewski, B. Foreman, H. Zhou, A. D. Portnoff, J.-H. Tian, A. Rehman, M. E. McGrath, R. E. Haupt, S. M. Weston, L. Baracco, H. Hammond, M. Guebre-Xabier, C. Dillen, M. Madhangi, A. M. Greene, M. J. Massare, G. M. Glenn, G. Smith, M. B. Frieman, Immunogenicity and protection of a variant nanoparticle vaccine that confers broad neutralization against SARS-CoV-2 variants.

*Nat. Commun.* **14**, 1130 (2023).

35. C. Liu, H. M. Ginn, W. Dejnirattisai, P. Supasa, B. Wang, A. Tuekprakhon, R. Nutalai, D. Zhou, A. J. Mentzer, Y. Zhao, H. M. E. Duyvesteyn, C. López-Camacho, J. Slon-Campos, T. S. Walter, D. Skelly, S. A. Johnson, T. G. Ritter, C. Mason, S. A. Costa Clemens, F. Gomes Naveca, V. Nascimento, F. Nascimento, C. Fernandes da Costa, P. C. Resende, A. Pauvolid-Correa, M. M. Siqueira, C. Dold, N. Temperton, T. Dong, A. J. Pollard, J. C. Knight, D. Crook, T. Lambe, E. Clutterbuck, S. Bibi, A. Flaxman, M. Bittaye, S. Belij-Rammerstorfer, S. C. Gilbert, T. Malik, M. W. Carroll, P. Klenerman, E. Barnes, S. J. Dunachie, V. Baillie, N. Serafin, Z. Ditse, K. Da Silva, N. G. Paterson, M. A. Williams, D. R. Hall, S. Madhi, M. C. Nunes, P. Goulder, E. E. Fry, J. Mongkolsapaya, J. Ren, D. I. Stuart, G. R. Screaton, Reduced neutralization of SARS-CoV-2 B.1.617 by vaccine and convalescent serum. *Cell*. **184**, 4220–4236.e13 (2021).

36. S. Chiba, S. J. Frey, P. J. Halfmann, M. Kuroda, T. Maemura, J. E. Yang, E. R. Wright, Y. Kawaoka, R. S. Kane, Multivalent nanoparticle-based vaccines protect hamsters against SARS-CoV-2 after a single immunization. *Commun Biol*. **4**, 597 (2021).

37. E. Takashita, N. Kinoshita, S. Yamayoshi, Y. Sakai-Tagawa, S. Fujisaki, M. Ito, K. Iwatsuki-Horimoto, P. Halfmann, S. Watanabe, K. Maeda, M. Imai, H. Mitsuya, N. Ohmagari, M. Takeda, H. Hasegawa, Y. Kawaoka, Efficacy of Antiviral Agents against the SARS-CoV-2 Omicron Subvariant BA.2. *N. Engl. J. Med.* **386**, 1475–1477 (2022).

38. S. Wilks, *titertools: Tools for maximum-likelihood based titer analysis, dealing with non-detectable titers* (Github; https://github.com/shwilks/titertools).

39. Racmacs, (available at https://acorg.github.io/Racmacs/).
